# Nuclear β-actin–dependent chromatin accessibility governs stem cell pluripotency and extracellular matrix gene programs to maintain cellular biomechanics for cell lineage decisions

**DOI:** 10.64898/2026.04.15.718829

**Authors:** Colin R. Campbell, Nadine Hosny El Said, Amani Ghassan Al Nuairi, Palanikumar Loganathan, Christopher Breunig, Muhammedin Deliorman, Zahraa Alraies, Esha Sadiq, Sabrina C. Desbordes, Liaqat Ali, Muhammad Shiraz Ali, Giuseppe A Saldi, Martin J. Lohse, Mazin Magzoub, Mohammad A Qasaimeh, Piergiorgio Percipalle

## Abstract

Pluripotency requires coordinated regulation of chromatin state, transcription and extracellular matrix (ECM) mechanics, but how these layers are integrated remains unclear. Here, using β-actin knockout mouse embryonic stem cells (mESCs) and a nuclear-targeted β-actin rescue, we identify nuclear β-actin as a key regulator linking chromatin accessibility to mechanosensitive control of cell fate. β-actin loss reduced OCT4, SOX2 and NANOG, broadly rewired the transcriptome and proteome and decreased accessibility at pluripotency regulatory regions. Integrated RNA-seq and ATAC-seq revealed coordinated dysregulation of stemness, ECM, mechanotransduction and early-lineage programs. These changes were accompanied by fibronectin and collagen upregulation, altered nuclear morphology, reduced Lamin A/C and mechanosensing proteins, increased nuclear YAP1 and greater ECM stiffness heterogeneity. Functionally, knockout cells displayed biased lineage specification, failed neuronal differentiation, ectopic cardiomyocyte-like differentiation, and markedly reduced teratoma growth with diminished ectodermal representation. Nuclear β-actin re-expression restored many molecular, mechanical and differentiation defects, although chromatin rescue remained incomplete. Together, these findings establish nuclear β-actin as an integrator of chromatin regulation and ECM-dependent mechanotransduction that preserves pluripotency and developmental competence.

## Introduction

The three-dimensional (3D) organization of the genome within the nucleus plays a pivotal role in orchestrating gene expression^1^. The genome’s spatial configuration brings regulatory elements such as enhancers, silencers, and insulators into physical proximity with gene promoters, enabling dynamic and context-specific control of transcription^2^. In pluripotent stem cells, this architectural flexibility is marked by a relatively open chromatin state, where chromatin compartments, topologically associating domains (TADs) and chromatin loops are more plastic, allowing for rapid access to genes required for maintaining pluripotency or initiating lineage commitment^3,4^. Key pluripotency transcription factors such as Oct4, Sox2, and Nanog not only regulate gene expression through direct binding to promoters and enhancers but also help shape chromatin conformation by recruiting chromatin remodelers and architectural proteins like CTCF and cohesin, thereby influencing chromatin architecture and 3D genome topology^5-7^. In this context, Oct4 and Sox2 are known to influence extracellular matrix (ECM) related processes through their control of upstream transcription factors and signaling pathways contributing to the degree of ECM stiffness during early developmental stages and differentiation. For instance, *Sox2* KO cells exhibit enrichment of integrin receptor-ECM interactions, which play an essential role in the epithelialization of the post-implantation epiblast^8,9^. As cells begin to differentiate, this chromatin landscape undergoes extensive reorganization: compartments switch from transcriptionally active (A) to repressive (B) states^10,11^, TAD intra-domain contacts change dynamically based on the compartment switching^10^, and new loops form between cell type-specific enhancers, super-enhancers and promoters, solidifying gene expression programs necessary for specialized functions^10-13^. Importantly, lineage-specific transcription factors cooperate with epigenetic modifiers to restructure the chromatin environment, stabilizing the identity of differentiated cells through long-range interactions and nuclear compartmentalization^14-17^. Thus, understanding the interplay between 3D genome organization, chromatin architecture and gene expression is essential for elucidating the molecular basis of pluripotency and differentiation^18-20^.

Recent studies have uncovered a critical role for nuclear actin, particularly β-actin, in the organization of the 3D genome, with profound implications for the regulation of gene expression programs that govern cell fate decisions. Once thought to function exclusively in the cytoplasm, β-actin has now emerged as a dynamic nuclear component essential for the spatial arrangement of chromatin^21^, including heterochromatin segregation at the nuclear envelope^22-24^, for transcription by all three eukaryotic RNA polymerases^21,25^ and it is required during co-transcriptional assembly of ribonucleoprotein complexes^26-28^. Recent work in mouse embryonic fibroblasts (MEFs) has shown that as a component of the SWI/SNF (BAF) chromatin remodeling complex regulating genomic deposition of the ATPase subunit Brg1, nuclear β-actin is required for the 3D genome organization at compartment level^23,24^. Nuclear actin depletion leads to recruitment of EZH2, the catalytic subunit of the polycomb repressive complex, leading to compartment switching, a general instability of the genome due to loss of promoter-enhancer interactions, and a significant increase in acetylation levels of histone H3 on lysine 27 (H3K27ac) specifically at gene regulatory regions^23,24,29^. Recently, we showed that in the absence of nuclear actin, increased H3K27ac at gene regulatory regions is recognized by the long non-coding RNA Meg3 that tightly binds to it, disrupting promoter-enhancer interactions^29^. In MEFs, this β-actin-dependent switch, with A compartments losing their structural integrity and transitioning toward B compartments, results in a more condensed and repressive chromatin state that directly reflects a fundamental transcriptional reprogramming affecting cell fate decisions through changes in chromatin accessibility^30-32^. These changes in cell fate decisions might be a consequence of changes in TGF-β signaling and ECM^33-35^. Recently, we showed that in MEFs lacking nuclear actin, ECM components involved in the process of TGF-β activation, such as fibronectin (Fn1)^36^, Ltbp1^37^, and thrombospondin-1 (Thbs1)^38^, were expressed at a significantly higher level^39^. Compatible with this scenario, β-actin KO MEFs display myofibroblast features, including enhanced abilities to produce and contract ECM^39^, likely due to enhanced TGF-β signaling. However, while nuclear actin regulators have been shown to play a role in pluripotency^40^, how nuclear β-actin regulates the ECM in the context of pluripotency is not known.

Here, we explored the importance of nuclear β-actin in regulating chromatin in the context of stemness and pluripotency. To this end, using CRISPR/Cas9, we generated mouse embryonic stem cells (mESCs) lacking β-actin (hereby referred to as KO) and a rescue mESC line where an N-terminally NLS-tagged β-actin construct is expressed in the KO background (hereby referred to as NLS). Transcriptional profiling of KO mESCs by RT-qPCR and RNA-seq reveals downregulation of pluripotency marker expression, which is partially rescued by exogenous nuclear β-actin in NLS mESCs. Results from ATAC-seq show that nuclear β-actin regulates pluripotency markers by controlling the state of chromatin and its accessibility in gene dense regions. Among the gene programs dysregulated in KO mESCs are those involved in lineage specification, ECM and mechanosensing. Their activity can be rescued by reintroducing β-actin in the nucleus. This dysregulation leads to marked changes in the ECM stiffness and the biomechanics of cells, altering their differentiation potential *in vitro* and *in vivo*, in a teratoma assay. We propose that by regulating chromatin architecture, the nuclear actin pool plays a fundamental role in pluripotency maintenance.

## Results

### Loss of nuclear β-actin affects stem cell pluripotency

To find out if the nuclear β-actin pool has a direct role in stem cell maintenance and differentiation, we generated a β-actin KO mESC line using CRISPR/Cas9, referred to as KO (Figure 1A). In addition, we established a mESC line to serve as a rescue cell line where an exogenous N-terminally NLS-tagged β-actin was reintroduced in the KO background, referred to as NLS (Figure 1A). As expected, in the NLS mESCs β-actin appears to be primarily in the nucleus in contrast to the WT condition where β-actin is found both in the cytoplasm and in the nucleus (Figure 1B,C). Importantly, the viability and cell count numbers were not changing between the different conditions (Figure 1D,E). However, the KO condition exhibited increased colony size compared to WT and this phenotype was rescued in the NLS condition. (Figure 1F,G). Next, we analyzed the extent of proliferation by staining for Alkaline Phosphatase (AP), a biomarker used for identifying active pluripotent stem cells, and Ki-67, a nuclear protein present during all active phases of the cell cycle (G1, S, G2, and mitosis) but absent in resting or non-dividing cells (G0 phase). Interestingly, in the KO condition we found a loss of AP staining (Figure 1H,I), but an increase in Ki-67 compared to WT and NLS mESCs (Figure 1J,K). Altogether, these results indicate that KO mESCs remain proliferative while losing stemness and pluripotency, suggesting a transition from an undifferentiated state toward differentiation. To confirm a possible loss of stemness, we next tested for expression of Oct4 and Sox2 in the β-actin KO background. Results from immunostaining and confocal microscopy show that neither are highly expressed in KO mESCs in contrast to WT and NLS mESCs (Figure 1L-N).

**Figure 1.**
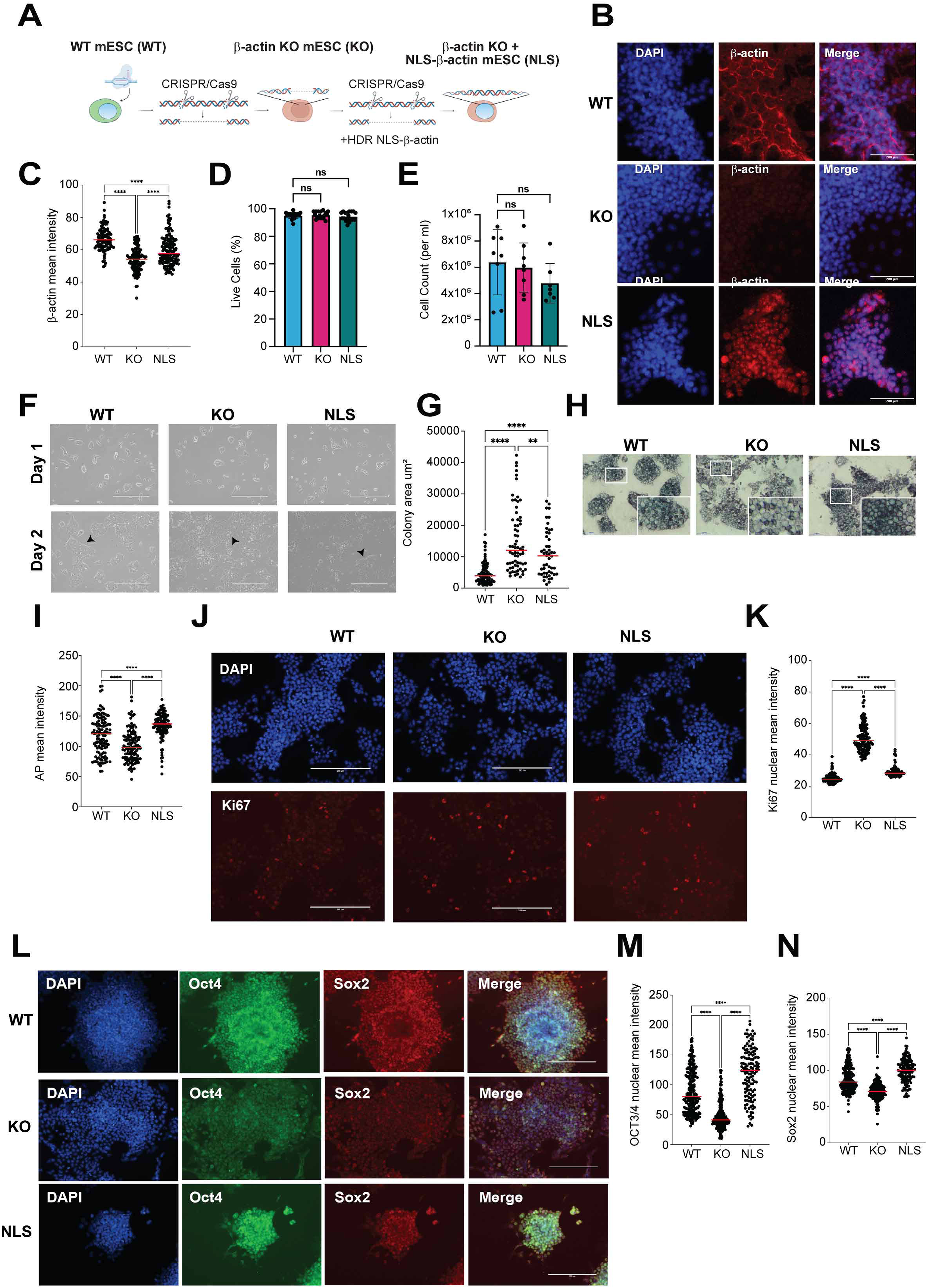
Nuclear β-actin depletion affects proliferation of mESC. **(A)** Schematic of WT, KO and NLS mESC constructs generated using CRISPR/Cas9. **(B)** Immunofluorescence images of undifferentiated WT, KO, and NLS mESC stained with an anti-β-actin monoclonal antibody (red) and DAPI (blue). Scale bar = 200µm. **(C)** Quantification of the mean fluorescence intensity in WT, KO and NLS mESCs. Each dot is a cell, n= 104 cells in WT, n= 110 cells in KO, n= 137 cells in NLS. One-way Anova, ****: p<0.0001 **(D)** Percentage of live cells after each passage. Each dot represents WT, KO and NLS mESCs collected for passaging, then counted before seeding and measured by automated cell counter. **(E)** WT, KO and NLS mESCs count after each passage measured by automated cell counter. **(F)** Brightfield images of WT, KO and NLS mESCs under maintenance conditions. Images show cells at Day 1 and 2 after seeding. Arrows indicate colonies that are well formed (WT and NLS) and poorly formed (KO). Scale bar = 400µm. (G) KO cells exhibit less colony formations than WT condition. NLS mESCs present an intermediary phenotype between WT and KO conditions. Each dot is a cell, n= 83 cells in WT, n= 69 cells in KO, n= 55 cells in NLS. One-way Anova, ****: p<0.0001, **: p=0.0049 **(H)** Alkaline phosphatase (AP) assay to monitor the state of cellular pluripotency. Decreased AP signal in KO condition compared to WT and NLS indicates decreased pluripotency under maintenance conditions. Scale bar = 100µm. **(I)** Quantification of AP signal. Each dot is a cell, n= 232 cells in WT, n= 270 cells in KO, n= 271 cells in NLS. One-way Anova, ****: p<0.0001, ns: P= 0.1873 **(J)** Cellular proliferation is not impaired upon β-actin depletion. Immunofluorescence images of WT, KO and NLS mESCs stained with Ki67 (red), and DAPI (blue), indicate no noticeable proliferative difference among the three genotypes. Scale bar = 200µm. **(K)** Quantification of Ki-67 signal obtained in (I). n= 145 cells in WT, n= 120 cells in KO, n= 110 cells in NLS. One-way Anova, ****: p<0.0001. **(L)** Immunofluorescence images of WT, KO and NLS mESCs stained with pluripotency factors Oct4 (green) and Sox2 (red). Scale bar = 200µm. **(M)** Quantification of Sox2 signal. n= 274 cells in WT, n= 197 cells in KO, n= 134 cells in NLS. One-way Anova, ****: p<0.0001. Scale bar: 100 um. **(N)** Quantification of Oct4 signal. n= 224 cells in WT, n= 237 cells in KO, n= 133 cells in NLS. One-way Anova, ****: p<0.0001. Scale bar: 100 µm.

Because expression of both Oct4 and Sox2 is rescued in the NLS condition, these data strongly suggest that the expression of these pluripotency factors depends on the nuclear pool of β-actin.

### Transcriptional profiling of mESCs reveals a direct role of nuclear β-actin in the maintenance of stemness

To assess nuclear β-actin’s role in stemness, we next isolated RNA from WT, KO and NLS mESCs for deep sequencing. Results from principal component analyses (PCA) performed on the normalized datasets show clear segregation among replicates of WT, KO and NLS mESCs as revealed by first (PC1) and second principal components (PC2), explaining 47% and 45% of the variation among the samples, respectively (Figure 2A). Clustering of the transcriptomic correlation matrix based on similarity confirmed that replicates of each condition cluster together and β-actin deletion leads to changes in gene expression (Figure 2B) which accompanies distinct phenotypic changes observed during the maintenance of the cells, particularly regarding colony formation (Figure 1F). To identify gene expression alterations associated with β-actin deletion in mESCs, we performed gene-by-gene analysis of covariance across three pairwise comparisons: β-actin KO versus WT, KO versus NLS, and WT versus NLS (Supplementary Figure 1). When comparing KO and WT mESCs, we identified 3657 significantly differentially expressed genes (p value < 0.05) (Figure 2C), 52% of which are downregulated in the β-actin KO condition (Supplementary Table 1). Gene Ontology (GO) analysis shows that the resulting changes in gene expression significantly disrupt several gene pathways (Figure 2D,E). GO terms related to ECM were found to be dysregulated in the comparison between KO and NLS but were not affected in the WT vs NLS groups comparisons (Supplementary Figure 2). These results suggest that loss of nuclear β-actin affects expression of ECM genes and this is rescued in NLS mESCs, altogether indicating a direct effect of nuclear β-actin on ECM-related gene programs. In agreement with GO term analysis, heat map of ECM genes (Figure 2F) shows that KO cells significantly upregulate the expression of *Col1a1* and *Col1a2*, known to be involved in cell differentiation by influencing the ECM environment^41^, as well as tenascin-C (*Tnc*), an ECM protein involved in proliferation, migration, and cell differentiation^42-44^. Interestingly, *Fbn2* is also upregulated and encodes fibrillin-2, a component of the ECM that plays a significant role in guiding cells towards specific differentiation pathways^45^. FBN2 interacts with proteins like UCMA and TGF-β, influencing cell differentiation and matrix formation. It plays a role in regulating cell fate, particularly during development and in stem cell niches. The *F*n1 gene, which codes for fibronectin, is also upregulated in the KO condition. Fibronectin plays a crucial role in various cellular processes, including cell adhesion, migration, and differentiation where it influences cell shape and maturation^46,47^. Compatible with the upregulation of ECM genes in KO mESCs (Figure 2F), we also found that several genes and gene families involved in stemness maintenance (Figure 2G) and differentiation (Figure 2H) are directly affected upon β-actin depletion. For instance, *Id1*, *Id2* and *Id3*, are all upregulated in KO cells (Figure 2G). *Id1* promotes T regulatory cell differentiation^48^, *Id1* and *Id3* expression are upregulated during differentiation, enhancing erythropoiesis^49^, whereas in neurogenesis Id proteins inhibit the activity of *bHLH* transcription factors that promote neural differentiation, thus preventing premature commitment to a neural lineage^50^. In mESCs, Id protein expression is influenced by external signals, such as growth factors and cytokines, which can trigger downstream signaling pathways that regulate Id protein levels^51^. This allows for a dynamic interplay between external cues and internal regulatory mechanisms controlling mESCs differentiation. Among those genes involved in lineage specification, we found that both *Tbx3* and *Tbx6* are upregulated in the KO cells (Figure 2H). *Tbx3* and *Tbx6* are both transcription factors involved in cell differentiation, but they play distinct roles in different developmental contexts and lineages^52-54^. *Tbx3* maintains pluripotency and self-renewal in stem cells while also promoting differentiation into specific lineages like mesendoderm^52,53^. *Tbx3* can both repress and activate gene expression, contributing to cell fate decisions during differentiation. On the other hand, *Tbx6* is critical for mesoderm formation and somite development^54^. Remarkably, almost all these ECM, stemness, and lineage genes are dysregulated in the KO mESCs but in many cases their levels are rescued when β-actin is reintroduced in the cell nucleus. This suggests that the nuclear pool of β-actin has a direct role in the maintenance of stemness and that loss of stemness upon nuclear β-actin depletion may be a consequence of altered gene expression (Figure 2F-H). Interestingly, although WT and NLS conditions exhibit similarities they also show differences in their transcriptomes since they do not exactly cluster in the PCA plots. It is, therefore, likely that the cells adjust their gene levels by compensatory mechanisms. For example, *dnmt3l* might be compensated by the dysregulation of *Tet1*, *Tet2* and *Tet3* that act through 5mc oxidation, and prdm14, which represses *Dnmt3b* genes and also acts through TET-mediated active DNA to maintain pluripotency. Moreover, levels of *Uhrf1* and *Dnmt1* genes are also rescued (Supplementary table 1), which might also compensate for *Dnmt3* gene levels. *Dppa3* is another example of a gene that is down regulated in KO condition and not rescued in the NLS condition. *Dppa3* is known to regulate stem cells self-renewal through regulating DNA methylation and stabilizing uhrf1 and Nanog despite not being rescued^55^. Thus, we speculate that loss of some gene activity that is not rescued in the NLS condition has probably to do with potential compensatory mechanisms which remain unclear at this stage and require further investigations using other models.

**Figure 2.**
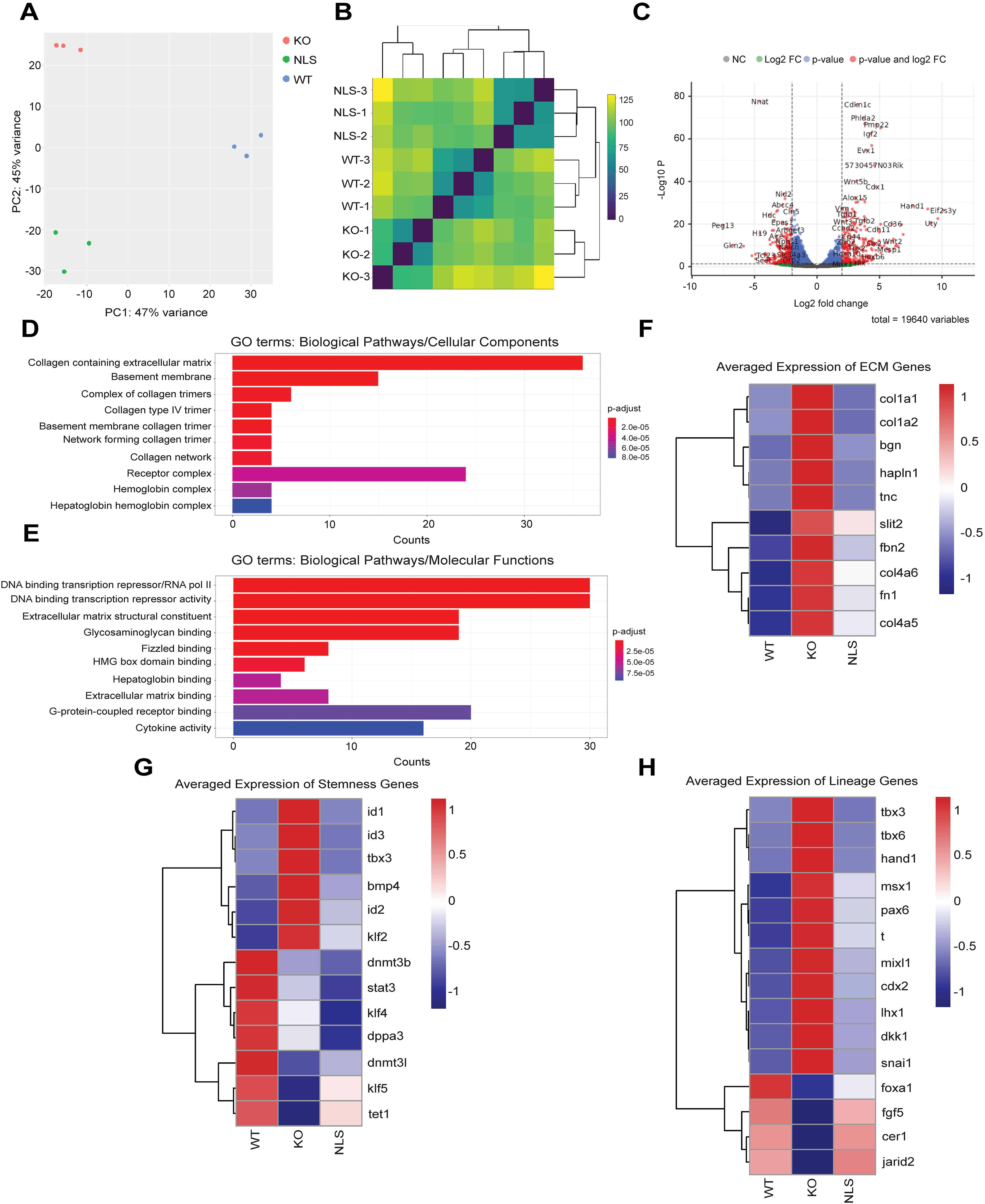
β-actin depletion in mESCs leads to transcriptional reprogramming. **(A)** PCA analysis shows that WT (blue), KO (red) and NLS (green) mESCs samples show distinctive clustered transcriptional profiles. **(B)** Hierarchical clustering analyses demonstrates that WT, KO and NLS mESCs samples exhibit distinct transcriptomes. **(C)** Volcano plot shows the differential expression of the normalized abundance of up and down regulated genes between KO and WT with both significantly up- and down-regulated genes. **(D,E)** Gene Ontology (GO) term enrichment analysis is shown as a bar plot of significantly overrepresented biological categories between KO and WT conditions. Bars represent enriched GO terms, and bar length indicates the degree of enrichment (or the number/proportion of genes associated with each term), highlighting the most prominent functional pathways in the gene set. Both cellular components and molecular functions are enriched for GO terms related to the extracellular matrix. **(F-H)** Nuclear β-actin depletion directly affects the expression of ECM genes as well as genes involved in stemness and lineages specification. Heat maps showing the differential expression patterns of genes associated with extracellular matrix (ECM) organization **(F)**, stemness **(G)**, and lineage specification **(H)** across WT, KO and NLS conditions. Each row represents a gene and each column represents a sample/group. Color intensity reflects average relative expression levels obtained from three independent biological replicates for WT, KO and NLS mESCs, with warmer colors indicating higher expression (red) and cooler colors (blue) indicating lower expression compared to the mean. Genes are grouped according to their functional categories to highlight coordinated transcriptional changes related to ECM remodeling, maintenance of stem cell properties, and commitment toward specific cellular lineages.

### Nuclear β-actin plays a direct role in regulating chromatin accessibility and expression of pluripotency genes

In β-actin KO MEFs, we reported that changes in heterochromatin revealed by elevated H3K9me3 and HP1α levels accompany significant transcriptional reprogramming ^22,23^. In KO mESCs we consistently observed increased H3K9me3 levels compared with WT and NLS mESCs (Figure 3A,B). Therefore, we performed ATAC-seq on WT, KO and NLS mESCs to find out if nuclear β-actin dependent changes in the transcriptome of KO mESCs depend on chromatin rearrangements. To obtain a global picture of changes in chromatin accessibility induced by nuclear β-actin loss, we first performed a differential peak calling analysis between WT and KO mESCs. When comparing KO and WT mESCs, we identified 684 significantly more accessible peaks and 180 significantly less accessible peaks, each showing a greater than twofold change in accessibility (Supplementary Table 2). Notably, in the KO condition we found decreased chromatin accessibility at TSSs compared to WT whereas NLS mESCs displayed a similar level of chromatin accessibility to WT (Figure 3C). This indicates that the reintroduction of β-actin solely in the cell nucleus rescues chromatin accessibility impacting downstream gene expression. GO term analysis of promoters with reduced accessibility revealed enrichment of several biological processes related to stemness maintenance, cell-fate commitment and regulation of differentiation (Supplementary Figure 3). Indeed, integrated genomic visualization (IGV) shows significant drops in the ATAC-seq signals of KO mESCs, particularly within the regulatory regions of *Oct4*, *Klf4*, *Sox2* and *Nanog*, compared to WT condition (Supplementary Figure 4), consistent with their decreased protein expression levels (Figure 1L-N). This loss of chromatin accessibility is only marginally detected in the NLS mESCs, indicating that by reintroducing β-actin in the cell nucleus, we potentially promote chromatin accessibility at the same loci. We, therefore, suggest that nuclear β-actin plays a direct role in regulating chromatin and thus expression of several key pluripotency genes.

**Figure 3.**
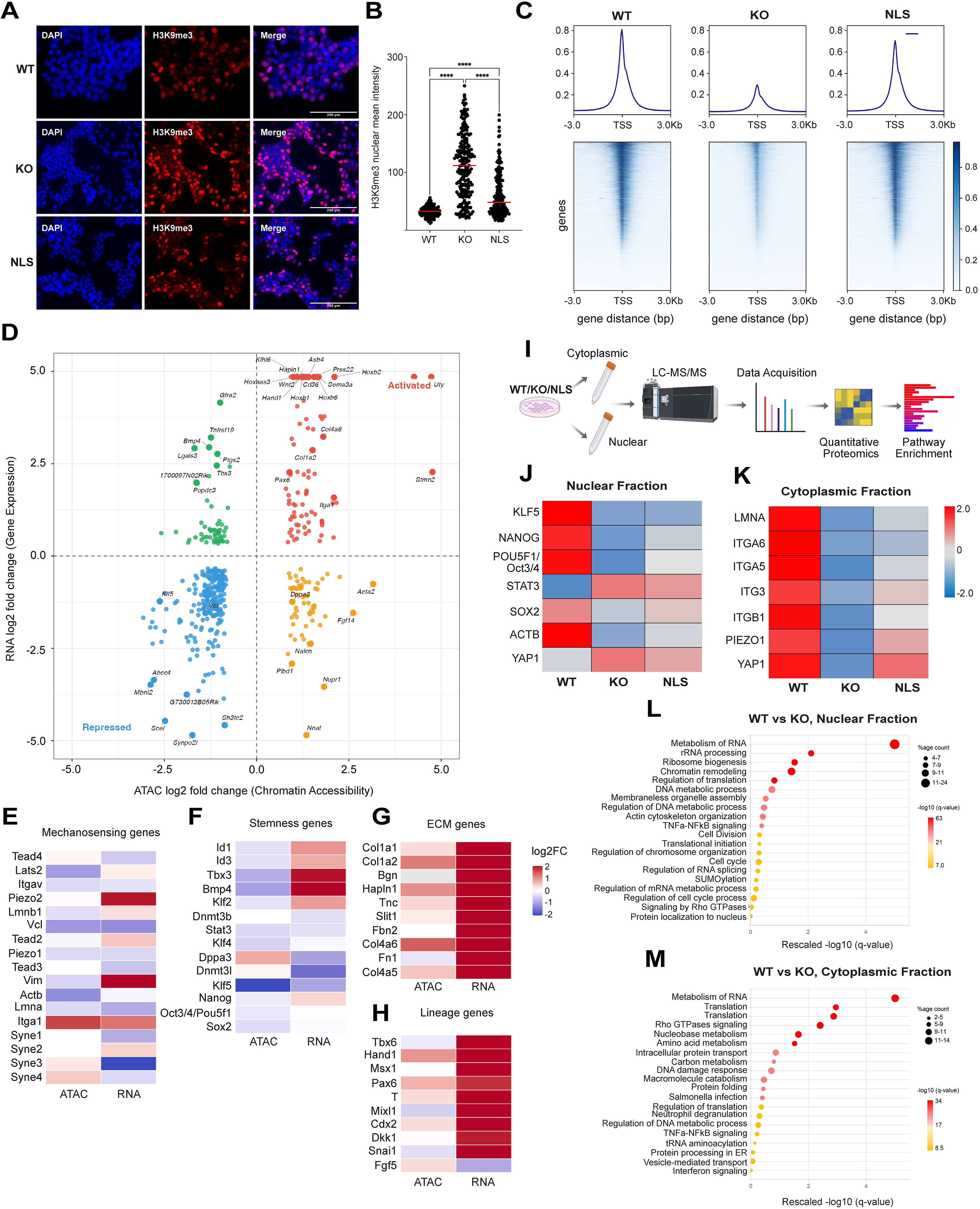
Chromatin architecture is directly affected by nuclear β-actin depletion, leading to increased heterochromatin levels and transcriptional reprogramming. **(A)** Immunofluorescence analysis with antibodies against H3K9me3 shows alterations in heterochromatin content in KO mESCs. **(B)** Quantification of H3K9me3 signal obtained in (A). n= 185 cells in WT, n= 228 cells in KO, n= 185 cells in NLS. One-way Anova, ****: p<0.0001 **(C)** ATAC-seq density plots and heatmaps of chromatin accessibility displaying scaled-read densities. Density plots show the distribution of normalized ATAC-seq signal intensity across genomic regions in WT, KO and NLS mESCs. Compared with WT, the KO condition displays loss of accessibility at TSS indicating altered chromatin openness at affected loci. In contrast, in the NLS condition, accessibility is restored back to WT levels, demonstrating recovery of chromatin structure. Heatmaps display ATAC-seq signal centered on TSS. Loss of accessibility in the KO relative to WT is clearly visible as reduced signal intensity. In the NLS condition, these accessibility patterns revert toward the WT state, indicating that the chromatin accessibility changes observed in the KO are largely rescued upon gene reintroduction of β-actin in the cell nucleus. The scale bar shows normalized RPKM. **(D)** Genome-wide integration of ATAC-seq and RNA-seq for KO versus WT mESCs. Each point is one gene that is significantly differential in both layers (RNA: adjusted p < 0.05 and |log FC| ≥ 0.25; ATAC: FDR < 0.05 and |log FC| ≥ 0.1; n = 1,624 genes). The x-axis is the ATAC promoter/nearest-TSS log fold-change and the y-axis is the RNA log fold-change. Quadrants define the integrated regulatory state: Activated (open + up, n = 280) and Repressed (closed + down, n = 431) are concordant (711 genes total); the off-diagonal quadrants (post-transcriptional, n = 211; Primed/open-not-expressed, n = 208) are discordant. Labelled genes are the curated ECM, stemness, lineage and mechanotransduction panel members detected in both layers, together with additional mechanosensor genes and the strongest genome-wide hits. The positive correlation between the two axes indicates that the majority of significant accessibility changes are matched by concordant expression changes. **(E-H)** KO-vs-WT chromatin accessibility and gene expression changes across the curated gene panel. Heatmap of KO-vs-WT log fold-changes for the 34 curated panel genes, grouped into four functional sets (ECM, Stemness, Lineage, Mechano; row-strip labels). The two columns show the change in chromatin accessibility (ATAC, left) and gene expression (RNA, right); for each gene the ATAC value is taken from its most significant peak (lowest DESeq2 adjusted p-value), the same peak used in the integration. Tile color encodes the log fold-change (red = increased in KO, blue = decreased in KO). These are the same fold-changes shown in the ATAC-vs-RNA integration scatter (panel D), allowing per-gene concordance to be read directly: genes red in both columns (e.g. ECM and lineage genes such as *Col1a2*, *Hand1* and *Cdx2*) gain accessibility and expression in KO, whereas genes with opposite signs (e.g. *Fn1*, *Tbx3*, *Bmp4*) change in only one layer. **(I)** Schematic overview of the experimental workflow. WT, KO, and NLS cells were subjected to subcellular fractionation to isolate nuclear and cytoplasmic protein fractions, followed by LC–MS/MS analysis. Quantitative proteomic data were analyzed to identify differentially abundant proteins and enriched biological pathways **(J-K)** Heatmaps showing the relative abundance (row-wise z-scores) of selected proteins in the nuclear (left) and cytoplasmic (right) fractions of WT, KO, and NLS cells. Red indicates higher relative abundance and blue indicates relative lower abundance. NLS cells show partial restoration of protein abundance compared with KO cells. The color scale represents normalized z-scores ranging from approximately −2 (blue) to +2 (red) **(L)** Bubble plot showing the top 20 significantly enriched biological pathways in the nuclear proteome of WT versus KO cells. Bubble size represents the number of proteins associated with each pathway, while color indicates enrichment significance (−log10 adjusted q-value), with darker red indicating greater significance. The x-axis shows the rescaled −log10 adjusted q-values, normalized to a 0-5 range for visualization. **(M)** Bubble plot showing the top 20 significantly enriched biological pathways in the cytoplasmic proteome of WT versus NLS cells. Bubble size represents the number of proteins associated with each pathway, while color indicates enrichment significance (−log10 adjusted *q*-value), with darker red indicating greater significance. The x-axis shows the rescaled −log10 adjusted *q*-values, normalized to a 0-5 range for visualization.

To examine whether the loss of β-actin leads to coordinated changes in chromatin accessibility and transcription, we integrated ATAC-seq and RNA-seq data, retaining genes independently significant in both assays (RNA adjusted p < 0.05, |log FC| ≥ 0.25; ATAC FDR < 0.05, |log FC| ≥ 0.1) (Supplementary table 3). When comparing KO versus WT (418 significant-in-both genes) conditions, accessibility and expression changes were predominantly aligned, with 304 genes (73%) falling in the concordant quadrants (79 Activated and 225 Repressed), giving a positive ATAC–RNA correlation (r=0.477) (Figure 3D). The concordant set of genes was enriched for stemness, ECM, mechanotransduction and early-lineage gene programs (Figure 3E-H). Among the ECM genes we identified *Col1a1, Col1a2, Tnc, Hapln1, Slit l*, as well as *Col4a6* and *Col4a5*, which form a known paired collagen-IV locus on chromosome X^56^. Since they share a bidirectional promoter, *Col4a6* and *Col4a5* are a cis-coregulated pair, in contrast to the other ECM genes which are not located in the same chromosomes and their likely relationship is trans-coregulation. Among the stemness genes we identified *Stat3* and *Klf5*. In mouse embryonic stem cells, STAT3 is known to be a major effector of the LIF–JAK–STAT3 pathway, which helps maintain naïve pluripotency and self-renewal. After activation by LIF, STAT3 is transported into the nucleus and it regulates transcriptional targets that preserve ESC identity and inhibit differentiation, especially mesoderm and endoderm programs^57^. Notably, Klf5 has been described among downstream targets associated with STAT3/LIF-mediated mESCs self-renewal, although Klf5 alone does not fully substitute for the complete LIF/STAT3 effect^58^. KLF5 is one of the Krüppel-like transcription factors associated with the pluripotency network, being expressed in mESCs, blastocysts and primordial germ cells. Compatible with evidence that Klf5 helps maintain the undifferentiated state partly by regulating Nanog and Oct3/4/Pou5f1 transcription, knocking down Klf5 perturbs mESC molecular identity and differentiation competence^59^. Within the concordant set of genes, we also identified *Hand1*, *Pax6*, *T/Brachyury/TBXT* and *Cdx2* which are all lineage-specifying transcription factors. Although they are not part of a single linear module, they sit in a branching lineage-decision network downstream of signaling pathways such as BMP, WNT, NODAL/Activin, FGF, Hippo/TEAD and chromatin state. Indeed, CDX2 has strong mechanistic links with the YAP/TEAD axis whereas T/Brachyury is linked to mechanics through WNT/β-catenin signaling. PAX6 and HAND1 are more likely downstream lineage outcomes shaped by the mechanical environment. Compatible with this, among the concordant genes exhibiting coordinated changes in chromatin accessibility and transcription, we also uncovered a set of genes involved in mechanosensing including *Itgav*, *Vcl*, *Piezo1*, *Lmna* and *Itga1*. We propose that these genes form a mechanosensing-to-transcription axis, functionally connected to KLF5, STAT3 and collagen/ECM genes^60^. As an example, ITGA1 encodes integrin α1, also called CD49a/VLA-1. It pairs mainly with ITGB1 to form α1β1 integrin, a cell-surface receptor for collagen and laminin with roles in adhesion, inflammation and fibrosis and through its direct regulation of focal adhesion signaling (FAK and RhoA) it acts on the YAP/TAZ pathway that in turn influences STAT3 and, thus, KLF5. By contrast, in NLS versus WT (425 genes) only 112 genes (26%) were concordant, and the two layers were weakly negatively correlated, indicating that in the NLS rescue line accessibility changes are largely uncoupled from matched expression changes (Supplementary Figure 5A-C). The KO versus NLS contrast had no significant-in-both genes and was excluded.

We next applied quantitative mass spectrometry to study if the changes in chromatin accessibility and gene expression upon β-actin depletion reflect the protein landscape in WT, KO and NLS mESCs (Figure 3I). Proteins were analyzed from both cytoplasmic and nuclear fractions, allowing us to assess how β-actin loss alters protein abundance and distribution across cellular compartments. After sample preparation, protein digestion, peptide labeling and mass spectrometry acquisition, the resulting data were processed and analyzed using statistical and pathway-based approaches to identify proteins and biological processes that changed across conditions (Supplementary Figure 6A-D). This analysis revealed that β-actin loss causes broad remodeling of the cellular proteome, affecting pathways linked to stem cell identity, nuclear organization, cytoskeletal regulation, mechanosensing and cell fate control. Importantly, many of these alterations were reduced or reversed when actin was restored in the nucleus, supporting the idea that nuclear β-actin plays a central role in maintaining the molecular state of pluripotent cells. The mass spectrometry results showed that, in the KO condition, key pluripotency factors including NANOG, POU5F1/OCT3/4 and SOX2 were downregulated at the protein level. KLF5 and STAT3 were also reduced, although to a lesser extent (Figure 3J; Supplementary table 4). Reintroduction of nuclear actin in NLS mESCs rescued, at least in part, the abundance of these factors (Figure 3J; Supplementary table 4,5). In addition, we detected reduced levels of mechanosensory and structural proteins, including Lamin A/C, integrins and PIEZO1, suggesting that β-actin loss also disrupts the ability of cells to sense and respond to mechanical cues (Figure 3K). Surprisingly, KO mESCs also showed a shift in YAP1 behavior from a predominantly cytoplasmic state toward a more nuclear function, indicating altered mechanotransduction signaling (Figure 3J,K). Pathway analysis revealed that these defects are likely to be due to impaired regulation of gene expression due to nuclear β-actin (Figure 3L,M; Supplementary Figure 6E,F). Overall, these findings support a model in which nuclear β-actin helps preserve pluripotency by coordinating transcriptional regulators, nuclear structure and mechanosensitive pathways.

Next, we integrated proximal promoter motif grammar, distal regulatory context and genome-wide ATAC-seq motif validation to explain how loss of the WT state and partial restoration by the NLS condition, affect chromatin accessibility at genes linked to ECM/niche, pluripotency-lineage control and mechanosensing/adhesion. Results from this analysis identified a potential 18-gene program comprising extracellular matrix and niche components, pluripotency and lineage regulators and mechanosensing and adhesion genes sharing a broad proximal-promoter motif grammar. Motifs associated with STAT, KLF/SP and HAND families were detected across all promoters, whereas TEAD/YAP, T-box, PAX, RUNX, SMAD, AP-1 and MEF2 motifs showed variable but generally extensive coverage; CDX motifs were comparatively less frequent, whereas SRF motifs were not detected in this promoter scan (Figure 4A). Integration of RNA expression with promoter accessibility and aggregate ATAC-seq signal across linked regions within ±20 kb showed the strongest correspondence in β-actin KO mESCs relative to WT. In the KO VS WT comparison, RNA and ATAC changes were highly correlated within the selected gene set, with Spearman coefficients of 0.82 for promoter accessibility and 0.87 for the aggregate regulatory interval (Figure 4B). Increased expression of several matrix-associated genes, including *Col1a1*, *Col1a2*, *Tnc*, *Slit1* and *Col4a5*, was accompanied by increased accessibility at their associated regulatory regions. Similar concordance was observed for lineage regulators such as *Hand1*, *Pax6*, T and *Cdx2*, whereas *Klf5* showed coordinated reductions in RNA expression and accessibility (Figure 4B). Re-expression of nuclear β-actin in NLS mESCs produced strong inverse RNA changes for many of these genes, consistent with partial transcriptional rescue (Figure 4B). However, the associated ATAC changes were generally smaller and RNA-ATAC correlations were substantially reduced in both the NLS-KO and NLS-WT comparisons. Residual differences between NLS and WT cells further indicated that restoration was gene selective rather than complete. Analysis of global ATAC-seq states showed that differential accessibility was associated with distinct combinations of transcription-factor motifs (Figure 4C). KO-associated opening was dominated by enrichment of SRF motifs, whereas regions that closed in KO cells were enriched for TEAD/YAP, STAT, KLF/SP and SMAD motifs. Regions that remained more accessible in NLS than WT cells were enriched for MEF2, TEAD/YAP, STAT, HAND and SRF motifs, while NLS-associated closed regions preferentially contained TEAD/YAP, STAT and KLF/SP motifs (Figure 4C). Given the general increase in H3K9me3 levels in the KO condition compared to WT and NLS, we next performed chromatin immunoprecipitation (ChIP) with an anti EZH2 antibody followed by deep sequencing (ChIP-seq). Parallel comparison of RNA and EZH2 signals identified five consistent inverse changes during nuclear β-actin rescue: *Slit1, Hand1, Pax6, T and Cdx2* were transcriptionally reduced in NLS relative to KO cells while EZH2 signal increased at their linked promoter and regulatory intervals (Figure 4D). These associations were retained across both promoter-restricted and aggregate interval analyses (Figure 4D), although they do not formally establish that EZH2 directly mediates transcriptional rescue. Finally, motif analysis of EZH2-defined regions showed that loci with higher EZH2 signal in KO cells had a significantly lower MEF2 motif burden, whereas no similarly robust associations were detected for the other candidate motif families (Figure 4E).

**Figure 4.**
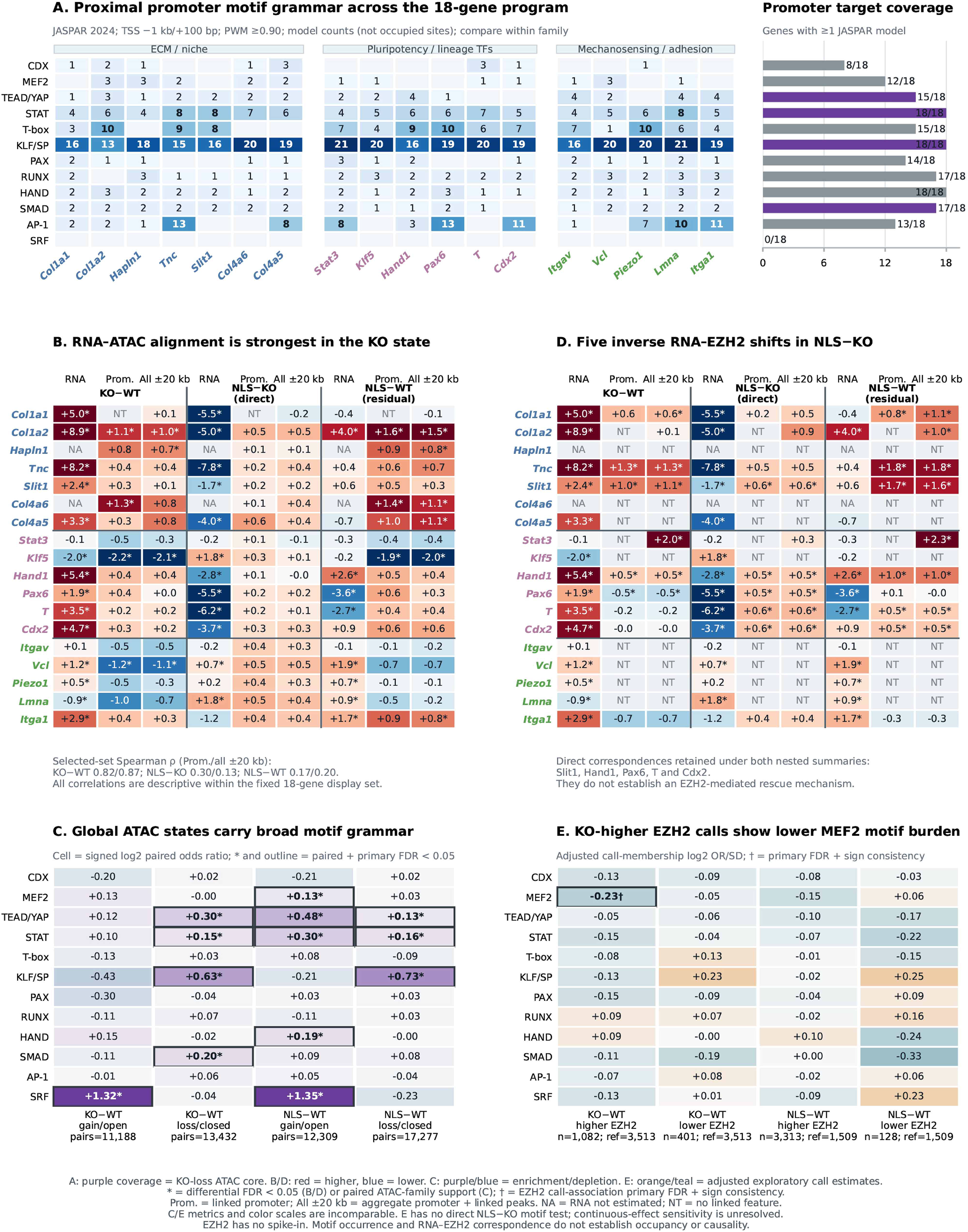
β-actin WT, KO, and NLS conditions were compared. Positive log fold changes indicate higher signal in the first-named condition. **(A)** Predicted promoter motif grammar across 18 genes grouped as ECM/niche, pluripotency/lineage transcription factors, or mechanosensing/adhesion. Proximal promoters (TSS −1 kb/+100 bp) were scanned using JASPAR 2024 position-weight matrices at a relative score threshold of 0.90. Each heatmap value is the number of family-member PWM models detected at least once in that promoter. Bars show the number of promoters containing at least one model from each family. Purple bars identify families supported among KO-associated ATAC-loss peaks in panel C. **(B)** RNA and linked ATAC effects for the three contrasts. “Prom.” represents the called promoter-peak model; “All ±20 kb” represents the aggregate promoter and linked proximal-peak model within ±20 kb of the TSS. Cells report log fold changes; red and blue indicate positive and negative effects, respectively, and asterisks denote assay-specific FDR < 0.05. Spearman correlations between RNA and ATAC effects were calculated across genes with complete measurements: KO−WT, ρ = 0.82 for promoters (n = 15) and ρ = 0.87 for ±20-kb aggregates (n = 16); NLS−KO, ρ = 0.30 and 0.13; and NLS−WT, ρ = 0.17 and 0.20, respectively. **(C)** Genome-wide motif-family enrichment across ATAC peaks gained or lost in KO−WT and NLS−WT comparisons. Differential peaks were uniquely paired with stable peaks matched for sequence length, GC content, baseline accessibility and distance to the nearest transcription start site; the number of matched pairs is shown below each column. Cells contain Haldane-corrected paired log odds ratios for motif-family occurrence, with purple and blue indicating enrichment and depletion. Asterisks and outlines denote enrichment supported at BH FDR < 0.05 by both the paired exact test and the primary matched-background enrichment analysis. **(D)** RNA and linked EZH2 effects displayed using the same gene order, contrasts and regional definitions as panel B. Cells report log fold changes, with asterisks indicating FDR < 0.05. In the direct NLS−KO comparison, Slit1, Hand1, Pax6, T and Cdx2 showed significant RNA and EZH2 changes in opposing directions under both promoter and ±20-kb aggregate models. **(E)** Motif-family burden in differential EZH2 interval sets. For each KO−WT or NLS−WT direction, FDR-significant EZH2 intervals were compared with reference intervals having adjusted P ≥ 0.10. Motif burden was defined as the fraction of family-member PWM models detected in each EZH2 interval. Cells show the adjusted log odds ratio of differential-set membership per standard-deviation increase in motif burden; models adjusted for EZH2 base mean, interval width, GC content, CpG observed/expected ratio and promoter overlap. Orange and teal indicate positive and negative associations. Daggers denote BH FDR < 0.05 with consistent effect direction across prespecified normalization and reference-definition sensitivities. Target and reference interval counts are shown beneath each column; NLS−KO was not tested in this panel. NA, RNA estimate unavailable; NT, no linked assay feature. Color intensity is scaled separately for RNA, ATAC and EZH2 and between panels C and E.

In summary, β-actin KO causes a broad collapse or redistribution of accessibility at regulatory elements controlling ECM, stemness and lineage genes, whereas the nuclear β-actin restoration partially rescues the regulatory network but leaves residual rewiring.

### Nuclear β-actin-dependent alteration of stem cell pluripotency is associated with changes in the ECM and nuclear morphology

We next studied if the observed β-actin-dependent chromatin changes led to alterations in the biochemical and biomechanical properties of the cellular microenvironment potentially associated with changes in mESCs pluripotency. Indeed, key ECM genes appear to be dysregulated in the β-actin KO condition. Analysis of the RNAseq data in WT, KO and NLS mESCs shows that ECM components such as tenascin (*Tnc*), fibronectin 1 (*Fn1*), *Col4a5*, and *Col4a6* are upregulated in the KO condition whereas in NLS mESCs their expression is rescued to levels similar to the WT condition (Supplementary Figure 5B; Supplementary Figure 7A-D). *Tnc* is an anti-adhesive glycoprotein often associated with tissue remodeling, inflammation, and development^61^, while *Fn1* plays a crucial role in cell adhesion, growth, migration, and differentiation by serving as a scaffold for other ECM molecules and by binding integrins on cell surfaces^62^. *Col4a5* and *Col4a6* encode alpha chains of type IV collagen, an essential structural component of basement membranes that contributes to ECM integrity and barrier functions^63^. Notably, loss of gene expression was accompanied by down regulation at the proteins level (Figure 3I-M; Supplementary Tables 4-5).

Immunofluorescence staining with antibodies to Fn1 showed increased protein levels in the ECM of KO mESCs (Figure 5A). Quantification of fluorescence intensity confirmed a significant increase in Fn1 signal in the KO condition relative to both WT and NLS cells, whereas no significant difference was detected between WT and NLS (Figure 5B). Similar increase in expression of ECM genes such as collagen and fibronectin was observed in MEFs upon nuclear β-actin depletion^39^ along with altered biomechanical properties such as, increased cell-surface stiffness, contractility, and enhanced activation of latent TGF-β1. In β-actin KO MEFs this drove myofibroblast differentiation, elevated α-SMA expression, ROS production and a profibrotic phenotype. Inhibiting TGF-β signaling partially reversed these stiffness-associated myofibroblast features. Latent TGF-β is mechanically activated by increased ECM stiffness and cellular contractility, creating a feed-forward loop that enhances mechanosensing and differentiation. So, in the KO mESCs we next assessed whether this extracellular matrix alteration is associated with changes in nuclear morphology by measuring nuclear circularity, a quantitative parameter that reflects how closely the nucleus resembles a perfect circle. To accurately define the nuclear boundary, cells were stained with an antibody against LAP2, a nuclear envelope marker. The LAP2 signal was then used to segment individual nuclei by applying a consistent intensity threshold, thereby generating nuclear masks from which circularity was calculated based on nuclear area and perimeter. Values close to 1 indicate a round nucleus, whereas lower values reflect a more elongated or irregular nuclear shape. This analysis showed that β-actin KO mESCs display a significant loss of nuclear circularity, with more elongated nuclei compared with both WT and NLS conditions (Figure 5C,D). The reduction in nuclear circularity is consistent with altered nuclear mechanical properties, as nuclear shape is determined by the balance between external forces transmitted by the cytoskeleton and the resistance provided by the nuclear lamina, chromatin organization, and the nuclear envelope. Thus, a less circular and more irregular nucleus may reflect changes in nuclear stiffness, lamina architecture, chromatin compaction or nucleo-cytoskeletal coupling. In line with this interpretation, Lamin A/C levels were reduced in β-actin KO cells compared with WT and NLS cells (Figure 5E-F), while YAP1 showed increased nuclear localization in the KO condition (Figure 5G-H), an observation that is compatible with our MS data (Figure 3). Together, these findings suggest that loss of nuclear β-actin leads to fibronectin accumulation, altered nuclear morphology and reduced Lamin A/C, creating a mechanobiological state compatible with increased extracellular matrix stiffness and activation of mechanotransduction pathways. In summary, nuclear β-actin is required not only for maintaining normal nuclear architecture, but also for preserving the mechanical balance between the extracellular matrix, the nucleus and mechanosensitive signaling in pluripotent stem cells.

**Figure 5.**
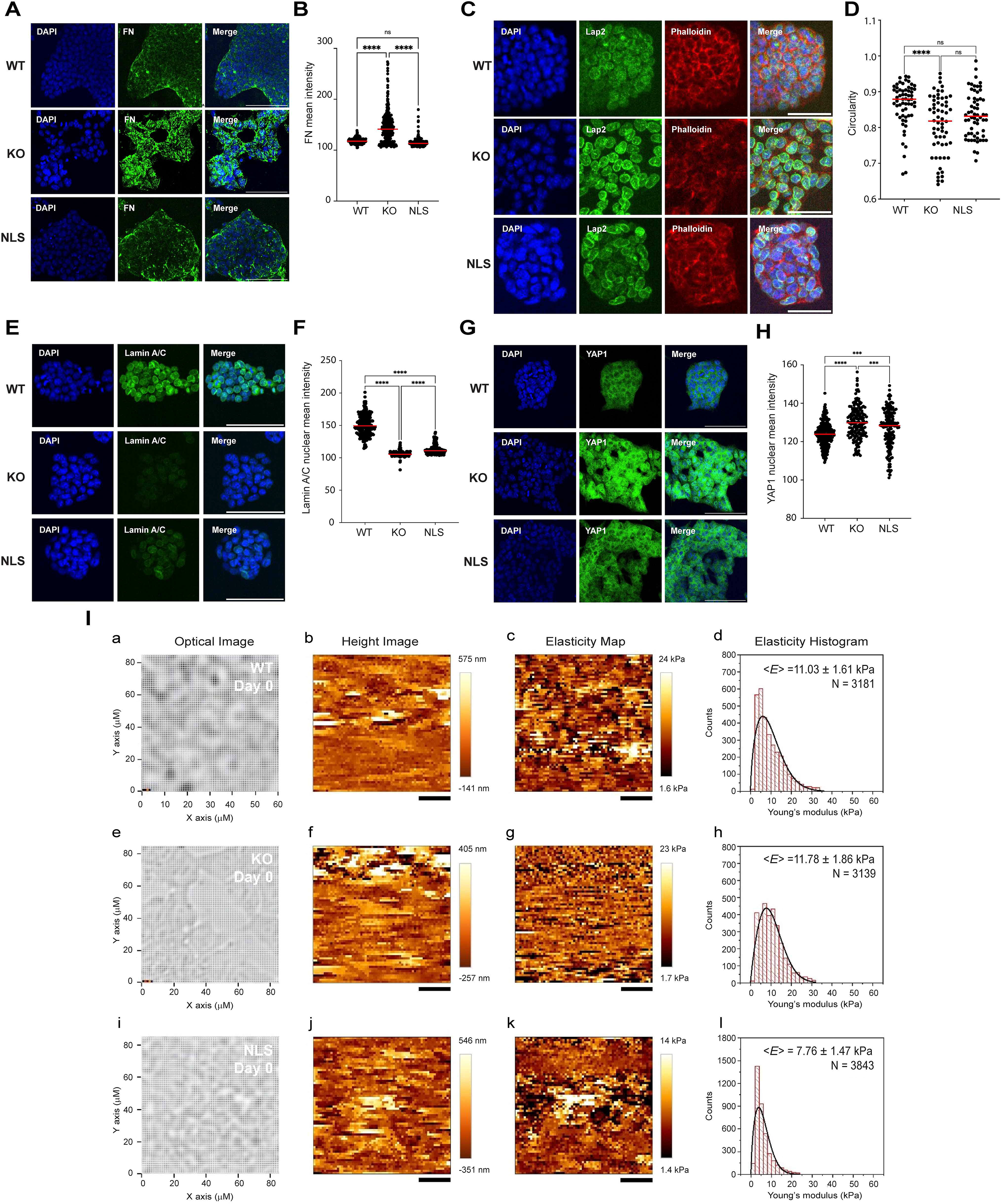
Nuclear β-actin depletion in mESCs leads to a significantly stiffer ECM. **(A)** Immunofluorescence staining for Fn (in green) and DAPI (in blue) shows increased protein levels upon β-actin depletion. Scale bar = 400µm. **(B)** Quantification of Fn signal: n= 263 cells in WT, n=276 cells in KO, n=295 cells in NLS. One-way Anova, **** : p<0.0001, ns: p=0.0989 **(C)** Immunofluorescence images of Lap2 (in green), Phalloidin (in red), and DAPI (in blue) shows increased nuclear circularity upon β-actin depletion. Scale bar: 50 um **(D)** Quantification of nuclear circularity : n= 61 cells in WT, n= 60 cells in KO, n=60 cells in NLS. One-way Anova, **** : p<0.0001, *** : p= 0.0008, ns: p= 0.5476. **(E)** Immunofluorescence staining for Lamin A/C (in green) and DAPI (in blue). Scale bar: 100 um **(F)** Quantification of Lamin A/C signal obtained in (E): n= 211 cells in WT, n= 191 cells in KO, n= 181 cells in NLS. One-way Anova, **** : p<0.0001. **(G)** Immunofluorescence staining for YAP1(in green) and DAPI (in blue) shows Yap1 increased nuclear localization upon β-actin depletion. Scale bar: 100 um. **(H)** Quantification of YAP1 nuclear signal obtained in (G): n= 247 cells in WT, n= 165 cells in KO, n=171 cells in NLS. One-way Anova, **** : p<0.0001, *** : p= 0.0008. **(I)** AFM-based ECM stiffness measurements at Day 0 for WT, KO, and NLS cells. (a, e, i) Optical images show the scanned areas for WT, KO, and NLS cells, respectively. (b, f, j) In the corresponding AFM height images, the ECMs show height < 1 µm. WT ECM exhibits a relatively uniform surface morphology, whereas KO and NLS ECMs display greater topographical variability. (c, g, k) Elasticity maps illustrate the spatial distribution of ECM stiffness within the scanned regions. (d, h, l) Histograms of Young’s modulus values derived from force measurements, with mean elasticity ⟨E⟩ ± SE obtained from Weibull probability density function fitting (solid black line). KO ECM exhibits a significantly higher mean elasticity than WT ECM and displays a broader stiffness distribution, indicative of increased mechanical heterogeneity. In contrast, NLS ECM exhibits the lowest mean elasticity and a narrower stiffness distribution, consistent with a softer and more mechanically uniform ECM. Elasticity maps are displayed using independently optimized color scales to visualize local stiffness variations and should not be used for direct quantitative comparison between conditions. AFM measurements were performed using three independent biological replicates per condition (n = 3). Scale bars = 20 µm.

Increased levels of ECM proteins are known to form compliant, developmentally plastic matrices instead of rigid load-bearing networks which would be compatible with a softer ECM, potentially with profound implications for mechanotransduction, including an effect on availability and presentation of growth factors such as TGF-β, Wnt, and FGF, further modifying stem cell behavior^64,65^. Atomic Force Microscopy (AFM) spectroscopy demonstrated that β-actin loss reduces the membrane stiffness in KO MEFs compared to WT^66^ and isolated KO nuclei have significantly lower stiffness than WT nuclei^67^. Reintroducing β-actin into KO nuclei restored both cellular and nuclear stiffness, demonstrating an essential role of the nuclear β-actin pool in maintaining mechanical strength of cells. Building on these observations, we next quantified ECM stiffness variations among WT, KO, and NLS mESCs using AFM. Our results show that in β-actin KO mESCs there are significant alterations in ECM stiffness heterogeneity. After defining the scan areas using optical images, AFM measurements were acquired in force-volume mode on cultured WT, KO, and NLS mESCs (Supplementary Figure 7). In force-volume mode, both topographical features and force data were simultaneously collected. During the process, AFM tip approached ECM surface, indented it until a constant loading force was reached, retracted from the surface, and repeated these steps across the scan area^68^. The relative deflection of the tip was recorded at each surface contact point, enabling the construction of a height images (Figure 5I, panels b, f, j). From the force curves, elasticity was calculated from the surface indentation at each pixel, and elasticity map was generated, displaying spatial stiffness variations (Figure 5I, panels c, g, k) and providing information on their relative distributions (Figure 5I, panels d, h, l). Because elasticity maps are displayed using independently optimized color scales to visualize local stiffness heterogeneity within each sample, direct visual comparison of color intensity between experimental conditions may not accurately reflect absolute stiffness differences. Therefore, quantitative comparisons were based on the Young’s modulus distributions obtained from the AFM measurements, while the elasticity maps were used as representative visualizations of local stiffness variations

AFM scans of WT, KO, and NLS mESCs showed significant stiffness variations (Table 1). The KO condition (11.78 ± 1.86 kPa) was ∼6% stiffer than WT ECM (11.03 ± 1.61 kPa), while NLS ECM had the lowest stiffness (6.69 ± 1.64 kPa). Elasticity maps (Figure 5I, panels c, g, k) demonstrated lower spatial variations in ECM stiffness, with WT showing the widest stiffness range at 32.37 kPa (1.62 kPa – 33.98 kPa), followed by KO ECM at 30.03 kPa (1.71 kPa – 31.74 kPa) and NLS ECM at 22.31 kPa (1.43 kPa – 23.74 kPa). Analysis of the Young’s modulus distributions revealed differences in ECM stiffness heterogeneity, with WT showing the widest stiffness range at 32.37 kPa (1.62-33.98 kPa), followed by KO ECM at 30.03 kPa (1.71-31.74 kPa) and NLS ECM at 22.31 kPa (1.43-23.74 kPa) (Figure 5I, panels d, h, l). Representative elasticity maps are shown in (Figure 5I panels c, g, k). Notably, the KO condition showed ∼15% and ∼27% greater variability in stiffness distribution compared to WT and NLS, respectively. These results suggest that loss of nuclear β-actin leads to increased ECM heterogeneity and stiffness.

**Table 1.**
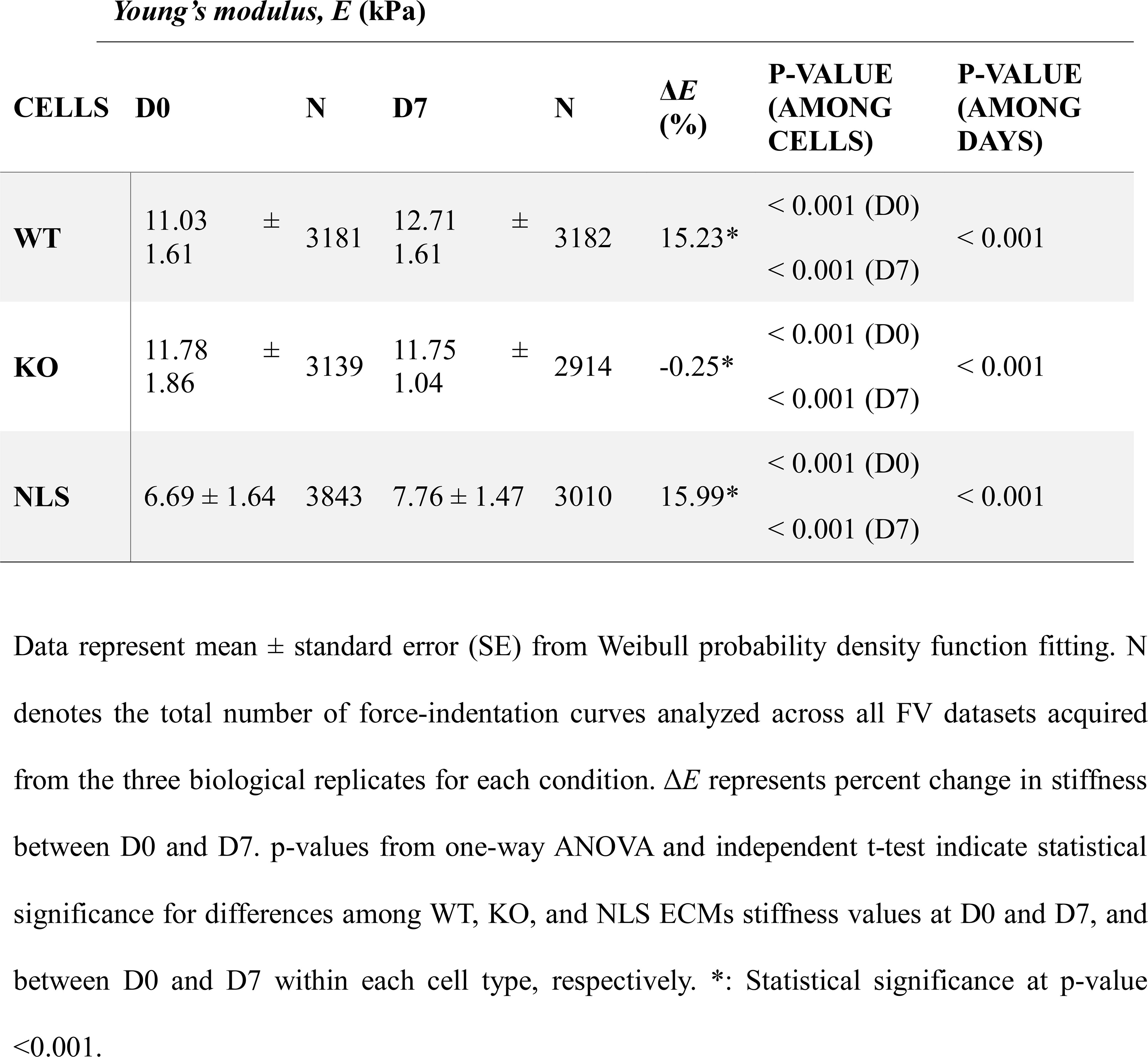
Summary of measured Young’s modulus of elasticity (*E*) among WT, KO, and NLS cells at Day 0 (D0) and Day 7 (D7).

Taken altogether these observations suggest that loss of nuclear β-actin leads to an impairment of force transmission through Lamin A/C as a plausible route by which changes in ECM stiffness could influence nuclear organization with a potential effect on differentiation.

### Nuclear β-actin depletion alters cell lineage commitment by affecting the ECM stiffness

Next, we explored whether loss of nuclear β-actin directly affects the differentiation potential of mESCs. The differentiation potential, also referred to as lineage restriction, is a fundamental process during embryonic development and cell fate specification as it consists in the gradual narrowing of a cell’s ability to become various other cell types. In pluripotent stem cells, such as mESCs, there is a high degree of plasticity - the ability to differentiate into any cell type from the three germ layers, ectoderm, mesoderm and endoderm. However, as these cells begin to receive cues from their microenvironment or intrinsic genetic programs, they start expressing lineage-specific transcription factors that signal the beginning of commitment to particular developmental pathways. From this perspective, embryoid bodies (EBs) are valuable tools for studying cell differentiation because, in vitro, they naturally form three-dimensional structures that mimic early embryonic development, allowing for the study of cell fate decisions and the formation of various cell types. Their ability to self-organize and differentiate into cells from all three germ layers makes them a relevant model for understanding how cells commit to specific lineages. To this end, we generated EBs by culturing WT, KO and NLS mESCs on gelatin-coated plates until they reached 70–80% confluency. Cells were detached by gentle dissociation to maintain cell viability and minimize clumping and then collected and resuspended in medium without LIF to achieve a seeding density of about 2–5×10 cells/ml. After forming hanging droplets on low-attachment culture dishes, WT, KO and NLS mESCs were allowed to aggregate into EBs (Figure 6A). The diameter of EBs can be a useful, though not definitive, measure to monitor the extent of differentiation. So, we followed their growth by wide field microscopy and after 2 and 3 days of growth, we measured the diameter to estimate the extent of differentiation (Figure 6B,C). Already after 2 days KO EBs were significantly larger than WT and the KO phenotype became even more striking after 3 days of growth with an almost 2-fold increase in diameter compared to WT condition (Figure 6B,C). No differences in growth were revealed between WT and NLS EBs (Figure 6B,C).

**Figure 6.**
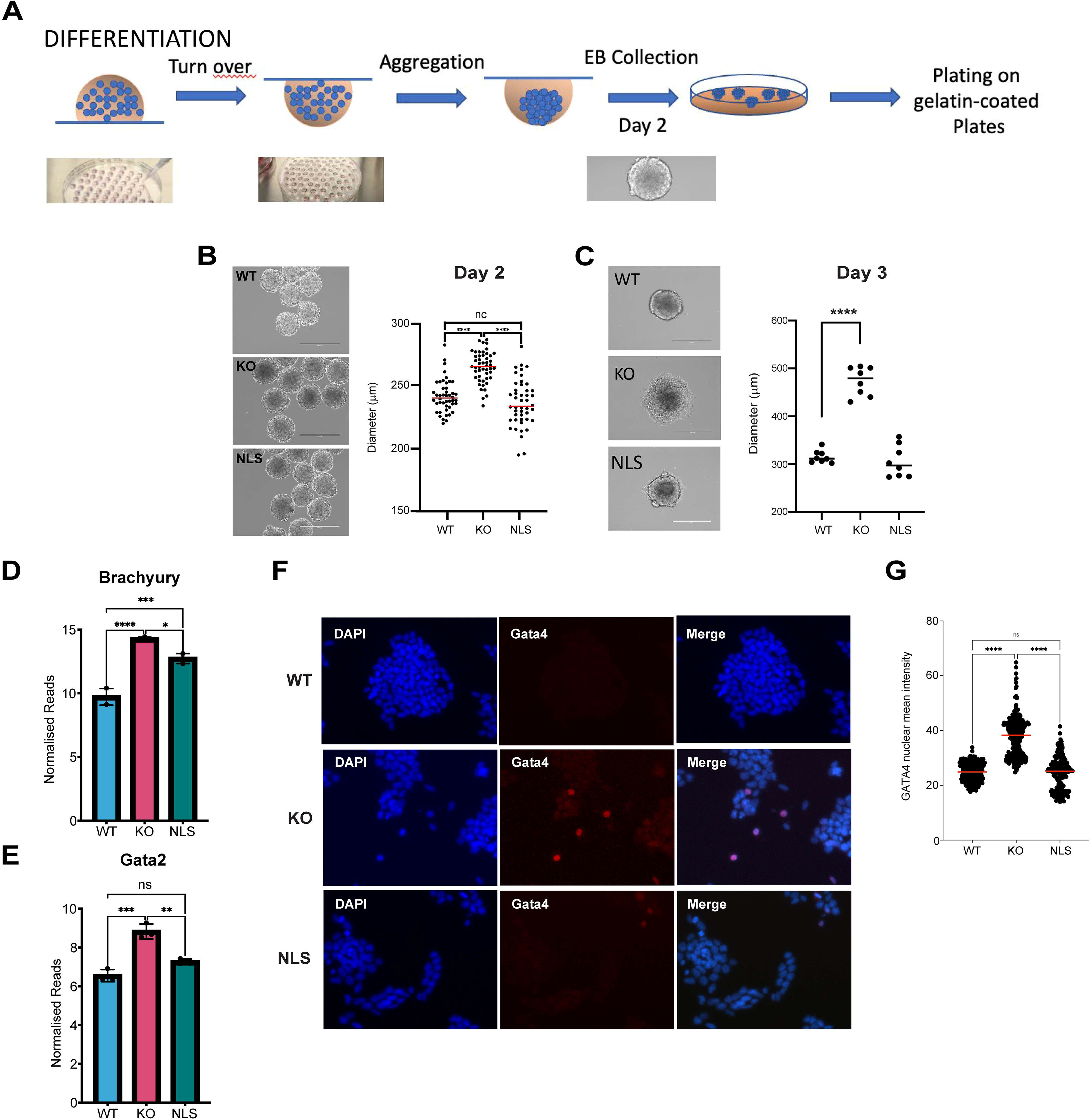
Differentiation Potential of mESCs to neuronal cell lineages. **(A)** Schematic of neuronal differentiation process using hanging droplets to form embryoid bodies (EB). **(B, C)** Brightfield images of EBs at Day 2 (hanging droplets) and 3 (plated on gelatin coated plates). Scale bar = 400 µm. Corresponding measurement of diameter of EBs reveal increased size of KO EBs. Each dot corresponds to 1 EB. One-way ANOVA indicates statistical significance in size difference. ns = non-significant (P>0.05); *P≤0.05; **P≤0.01; P***≤0.001; ****P≤0.0001. **(D-E)** Normalized reads from RNAseq of *Brachyury* and *Gata2* show increased expression of both factors upon β-actin depletion. **(F)** Immunofluorescence images of Gata4 indicate higher basal expression of this mesodermal factor in KO cells. Scale bars = 400µm. **(G)** Quantification of Gata4 signal obtained in (G): n= 330 cells in WT, n= 283 cells in KO, n= 183 cells in NLS. One-way Anova, **** : p<0.0001, ns: P= 0.7522.

To gain further insights, we studied the expression of Brachyury (also known as TBXT), GATA2 and GATA4 in undifferentiated mESCs (Figure 6D-F). Their increased expression is a key indicator of the transition from a proliferative state of self-renewal to a differentiating state^69^. Brachyury is a T-box transcription factor essential for mesoderm formation and differentiation^70^. Its upregulation indicates that a previously uncommitted cell is now biased toward a mesodermal fate, which includes tissues like muscle, bone and blood. Similarly, GATA2 is a zinc finger transcription factor involved in hematopoietic (blood) development and marks early blood and endothelial lineage commitment^71^. GATA4, on the other hand, plays a critical role in cardiac development and endodermal differentiation^72^. Increased expression of these factors is not merely a marker of lineage choice; it also enforces that choice. These transcription factors activate gene programs necessary for specific lineage differentiation while repressing genes associated with pluripotency and alternative fates. Therefore, as levels of Brachyury, GATA2 and GATA4 rise, the cell’s fate becomes more narrowly defined, and its ability to adopt other identities diminishes. This loss of differentiation potential means that the cell has passed a threshold beyond which it can no longer revert to a pluripotent state or easily transdifferentiate into other lineages. Compatible with an alteration in their differentiation potential, we found that in the KO condition the levels of *Brachyury* and *Gata2* mRNAs were significantly upregulated compared to WT condition and these levels were rescued in the NLS mESCs to a level comparable to WT (Figure 6D,E). Using immunofluorescence microscopy, we confirmed that GATA4 protein levels are upregulated in the KO condition compared to WT and NLS mESCs (Figure 6F). Quantification of fluorescence intensity confirmed a significant increase in GATA4 signal in the KO condition relative to both WT and NLS cells, whereas no significant difference was detected between WT and NLS (Figure 6G). Altogether these results suggest nuclear β-actin dependent alteration in the mechanisms underlying cell lineage decisions.

Next, we set out to differentiate WT, KO and NLS mESC to neurons to study if the nuclear β-actin dependent transcriptional reprograming negatively regulates key differentiation pathways. Differentiation into neuronal lineages was performed over a 14-day time course and cells were analyzed cells were analyzed at the undifferentiated stage at day 0 and when terminally differentiated at day 14 by immunostaining with antibodies against PAN neuronal markers βIII-tubulin (Tuj1) and MAP2 followed by confocal microscopy. After 14 days of directed differentiation toward a neuronal lineage, clear differences were observed between WT and KO conditions. While as expected WT and NLS cultures displayed strong expression for both βIII-tubulin (Tuj1) and MAP2, in KO cultures both proteins could barely be detected (Figure 7A-C). In WT and NLS conditions, immunostaining revealed extensive neurite outgrowth and a complex neuronal network morphology, consistent with increased cells’ maturation into neurons. The presence of both early (βIII-tubulin) and more mature (MAP2) neuronal markers indicates efficient progression through the differentiation program in contrast to the KO condition (Figure 7A, B). Furthermore, KO mESCs retained a non-neuronal morphology, suggesting that nuclear β-actin is essential for the acquisition of neuronal identity. Altogether these findings indicate that the observed differentiation defect in the KO cells is specifically attributable to loss of nuclear β-actin and can be functionally rescued upon NLS-tagged β-actin expression.

**Figure 7.**
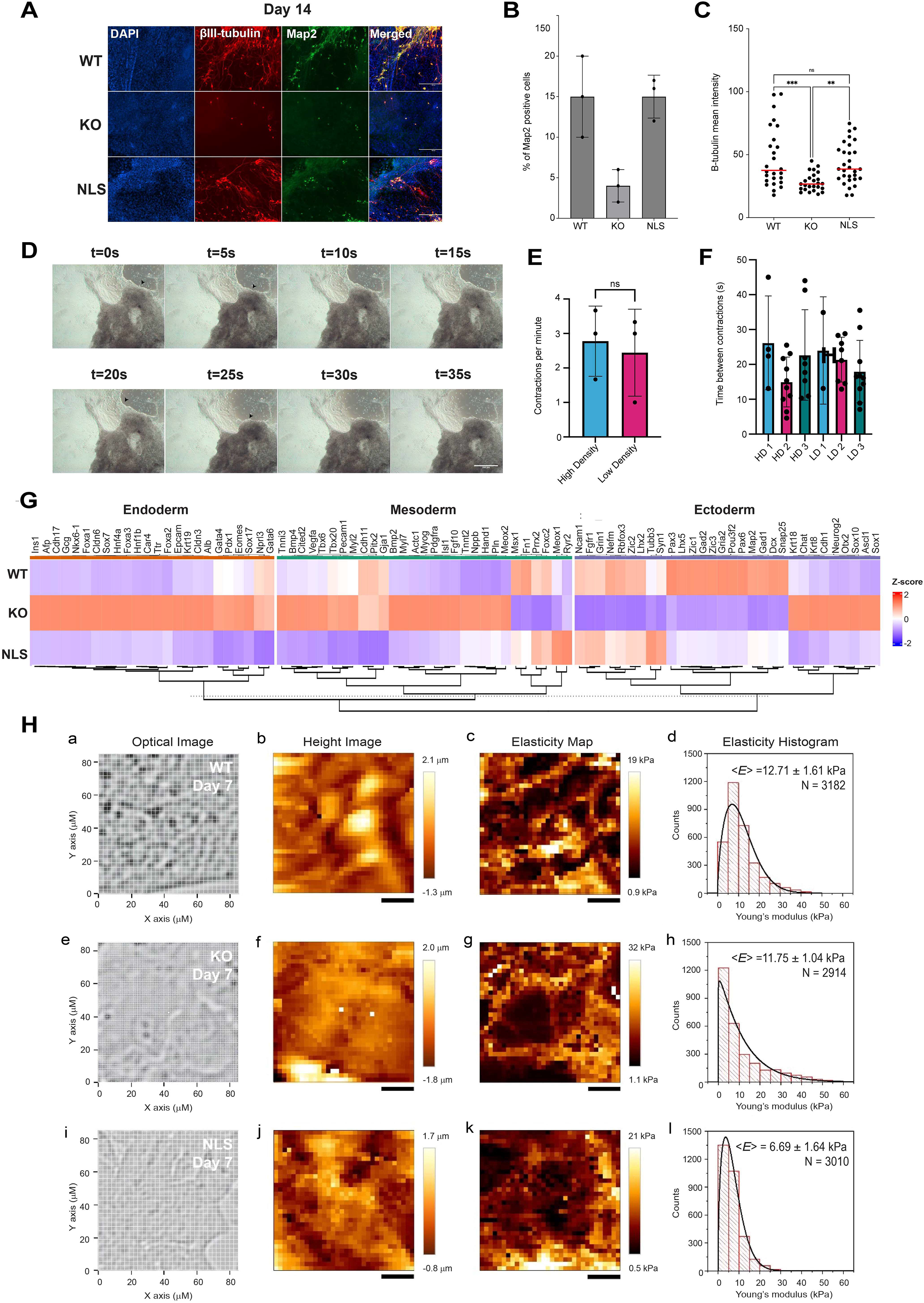
β-actin KO mESCs fail to undergo neuronal differentiation and preferentially give rise to cardiomyocyte-like cells, suggesting a general impairment in cell lineage definition. **(A)** Immunofluorescence staining with antibodies against the neuronal markers βIII tubulin and Map2 on WT, KO and NLS mESCs neuronal differentiation at D14. While at D0 none of the conditions - WT, KO and NLS mESCs - express βIII tubulin and Map2. **(B)** Percentage of Map2 positive cells quantified using the Map2 signal obtained in (A): n= 19 cells in WT, n= 23 cells in KO, n= 21 cells in NLS. One-way Anova, **** : p<0.0001, ns: p= 0.2766. (C) Quantification of βIII-tubulin signal: n= 26 cells in WT, n= 25 cells in KO, n= 30 cells in NLS. One-way Anova, *** : p= 0.0003, **: p= 0.069, ns: p= 0.5306. **(D)** Snapshots of contracting KO cells on day 14 after neuronal differentiation. Snapshots are taken 5 seconds apart. Contraction is highlighted by the black arrows, which highlight regions going from light to dark as the cells contract and become denser. This is apparent in the first two frames (t=0s and t=5s) and frames five and six (t=20s and t=25s). Additionally, in the sixth frame (t=25s), we see a portion of the attached cells detach from the plate, indicating the strength of the contraction. Scale bar = 200μm. **(E,F)** 3-minute recordings of various contracting KO cells from multiple wells of HD (n=3) and LD (n=3) seeded EBs were manually analyzed for the number of contractions per minute as well as the time between contractions. This was determined with a stopwatch that had a lap feature, so the time between contractions could be calculated. **(G)** Heatmap of selected differentially expressed Endoderm, Mesoderm and Ectoderm genes among WT, KO and NLS mESCs shows that reintroduction of NLS-tagged β-actin rescues compromised expression levels in the KO condition. Rows represent marker genes and columns represent the three experimental groups Expression values are presented as row Z-scores, with red indicating relatively higher expression and blue indicating relatively lower expression. **(H)** AFM-based ECM stiffness measurements at Day 7 for WT, KO, and NLS cells. (a, e, i) Optical images show the scanned areas for WT, KO, and NLS ECMs, respectively. (b, f, j) AFM height images reveal increased (> 1 µm) topographical variations compared to Day 0, with WT and KO ECMs displaying greater relative height variations (> 2 µm). (c, g, k) Elasticity maps illustrate the spatial distribution of ECM stiffness within the scanned regions. (d, h, l) Histograms of Young’s modulus values derived from force measurements, with mean elasticity ⟨E⟩ ± SE obtained from Weibull probability density function fitting (solid black line). WT and KO ECMs exhibit significantly different mean elasticity values, with KO ECM displaying a broader stiffness distribution indicative of increased mechanical heterogeneity. In contrast, NLS ECM exhibits the lowest mean elasticity and a narrower stiffness distribution, consistent with a softer and more mechanically uniform ECM. Elasticity maps are displayed using independently optimized color scales to visualize local stiffness variations and should not be used for direct quantitative comparison between conditions. AFM measurements were performed using three independent biological replicates per condition (n = 3). Scale bars = 20 µm.

Next, we subjected WT, KO and NLS mESCs to an identical directed neuronal differentiation protocol but this time cultures were monitored longitudinally by live-cell imaging to track morphology and lineage progression. WT and NLS mESCs followed the expected neuronal trajectory, forming neurite-bearing networks like in Figure 7A (images not shown). In contrast, KO cultures reproducibly diverged, generating compact, contracting aggregates and sheet-like regions with spontaneous rhythmic beating, consistent with an ectopic mesodermal/cardiomyocyte-like fate. Beating behavior was recorded by live microscopy (Supplementary Video 1), and contraction frequency was quantified from video tracks (beats per minute) across multiple fields and biological replicates. Contraction frequencies were summarized and plotted to compare differentiated cells from High- or Low-Density KO EBs, 2x10^5^ or 1x10^6^ cells per mL when seeding the EBs, respectively (Figure 7D-F). Taken together, these results suggest that by regulating expression of gene programs at the chromatin level, the nuclear β-actin pool is required for cell lineage specification. Indeed, RNA-seq analysis on total RNA isolated from WT, KO and NLS mESCs at day 14 of neuronal differentiation shows extensive differential expression of genes associated with germ layer specification - endoderm, mesoderm, and ectoderm - between WT and KO conditions (Figure 7G; Supplementary Figures 8 and 9). In WT mESCs, the transcriptional profile shows strong upregulation of ectodermal and neurogenic genes, consistent with proper neuronal differentiation, alongside low expression of mesodermal and endodermal markers. In contrast, KO mESCs exhibit a clear global transcriptional reprogramming, with reduced expression of ectodermal genes and inappropriate activation or persistence of mesodermal and endodermal signatures, indicating a disruption in lineage commitment. Among the ectodermal genes we found genes involved in early ectoderm induction such as *Ncam*, *Zic1/2/3*, *Pax6, and Fgfr1* which primarily contribute to ectoderm formation by specifying neuroectoderm against surface ectoderm^73-76^. We also found *Pax3*, which together with *Zic1* and *Zic3* is important for plate border by specifying neural crest and pre-placodal ectoderm^74,76^. Compatible with the observed loss of differentiation potential, we also found that *Tubb3*, *Map2*, *Rbfox3*, *Syn1*, and *Dcx,* known to directly contribute to neural maturation and neurogenesis, are downregulated^77,78^. Notably, this aberrant transcriptional state is significantly rescued in the NLS condition, where gene expression patterns are restored to levels comparable to wild type, with re-established ectodermal gene expression and suppression of non-neural lineage markers, demonstrating recovery of proper differentiation.

The formation of mesodermal or cardiomyocyte-like tissue is compatible with increased ECM stiffness, a well-established regulator of cell fate through mechanotransduction pathways^79,80^. Increased stiffness enhances cytoskeletal tension and activates signaling molecules such as YAP/TAZ, RhoA/ROCK, and integrin-mediated pathways, which can promote mesodermal differentiation from pluripotent stem cells^81,82^. Cardiac lineage specification, in particular, is favored by intermediate-to-stiff substrates that *in vivo* approximate the mechanical properties of developing myocardium^82^. While excessively high stiffness may impair maturation or function, moderately increased ECM stiffness can support mesodermal commitment and cardiomyocyte-like differentiation, especially when combined with appropriate biochemical cues. Given the moderately increased ECM stiffness and differential expression of ECM genes in the KO condition after 4 (Supplementary Figures 10 and 11, Supplementary table 7) and 7 days of differentiation (Supplementary Figures 12 and 13), we next applied AFM to WT, KO and NLS mESCs at day 7 (D7) into the differentiation protocol. By D7, WT and NLS ECMs displayed significant stiffness increases to 12.71 ± 1.61 kPa (+15.23%) and 7.76 ± 1.47 kPa (+15.99%), respectively, compared to undifferentiated cells (day 0, D0). In contrast, the KO condition showed only a minimal change in stiffness (-0.25%) relative to D0 (Table 1).

Analysis of the Young’s modulus distributions indicated substantial ECM remodeling during differentiation. Elasticity histograms showed that WT (0.28-49.35 kPa) and KO (0.48-68.44 kPa) ECMs exhibited broader stiffness distributions than NLS ECM (0.57-43.14 kPa) (Figure 7H). Consistent with the histogram analysis, representative elasticity maps illustrating the spatial distribution of ECM stiffness are shown in (Figure 7H panels c, g, k). As observed at D0, the KO condition exhibited the greatest stiffness heterogeneity, with a stiffness range of 67.96 kPa, corresponding to ∼39% and 60% greater variability than WT ECM (49.07 kPa) and NLS ECM (42.57 kPa), respectively.

Taken altogether these findings suggest that by regulating gene expression through a chromatin-based mechanism, nuclear β-actin controls downstream genes implicated in ECM stiffness and directly affects mESCs differentiation.

### Nuclear β-actin depletion negatively regulates the proliferative state of mESCs *in vivo*

We next investigated whether depletion of nuclear β-actin inhibits mESCs proliferation *in vivo*. To this end, we assayed the ability of WT, KO and NLS mESCs to form teratomas. These tumors are known to contain all three germ layers (ectoderm, mesoderm, endoderm) when transplanted into immunodeficient mice and, therefore, represent an ideal model to study mESCs pluripotency and self-renewal^83^.

We injected WT, KO and NLS mESCs subcutaneously into female BALB/c nude mice and monitored teratoma formation for 18 days (Figure 8A). Animals injected with WT mESCs exhibited prominent abdominal enlargement consistent with substantial tumor burden, whereas mice injected with NLS mESCs displayed moderately sized masses. In contrast, mice injected with KO mESCs developed markedly smaller tumors with minimal abdominal distension (Figure 8B, C). Comparison of the tumors among the WT, KO, and NLS groups over the 18-day duration of the experiment revealed substantial growth in the WT and NLS groups relative to the KO group (Figure 8D). Specifically, tumor measurements by digital caliper showed rapid expansion in the WT group beginning around days 8–10, reaching a final volume of 1,717 ± 41 mm³ by day 18 (Figure 8E). In contrast, tumors in the NLS group exhibited slower growth kinetics, attaining a volume of 1,169 ± 156 mm³ at endpoint, corresponding to an estimated 37.98 % reduction (0.7-fold) compared with WT (Figure 8E). Notably, the KO group showed markedly impaired tumor growth, with a final tumor volume of 174 ± 123 mm³ (Figure 8E). This represents approximate reductions of approximately 10- and 8-fold compared to the WT and NLS groups, respectively with no significant differences in the mice body weight between all the groups (Figure 8F). Collectively, these findings indicate that depletion of nuclear β-actin profoundly restricts teratoma growth compared to the WT condition, and this phenotype is partly rescued when β-actin is reintroduced in the nucleus of KO cells.

**Figure 8.**
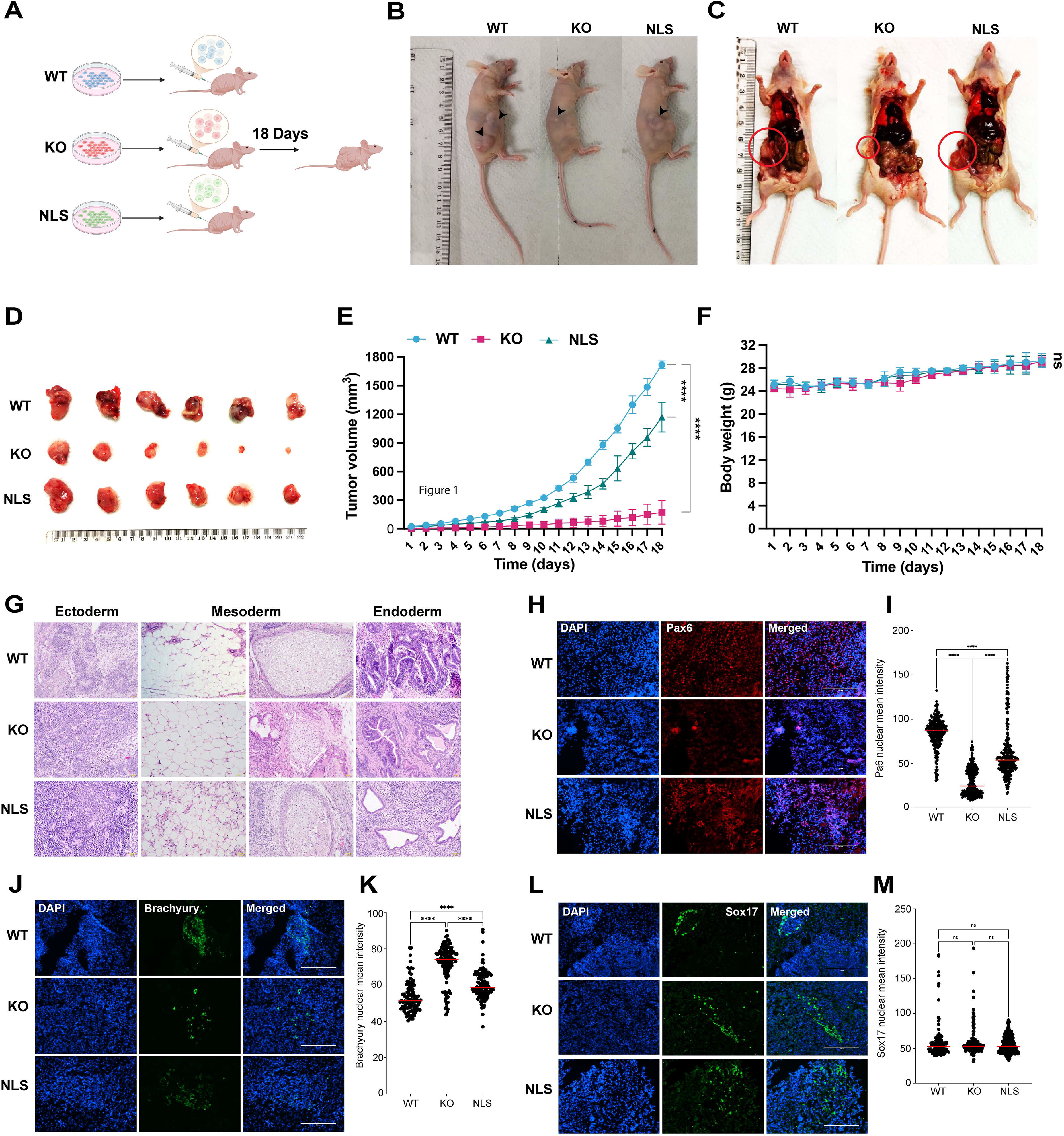
*In vivo* teratoma studies show that loss of β-actin directly affects the differentiation potential of mESCs. **(A)** Schematic of teratoma assay. Nude mice were randomly allocated to three experimental groups (WT, NLS, KO; n = 6 per group) and implanted subcutaneously in the right flank with 4 × 10 mouse embryonic stem cells. After 18 days, the mice were sacrificed and tumors were dissected for further analysis. **(B)** Nude mice after 18 days post-injection. Black arrowheads indicate visible subcutaneous tumors. **(C)** Red circles indicate subcutaneous teratomas in dissected nude mice. **(D)** Excised tumors from the WT, KO, and NLS groups illustrating differences in tumor size. **(E)** Longitudinal tumor volume measurements over the 18-day duration of the experiment show distinct growth profiles across the WT, KO, and NLS groups. Statistical comparisons were performed using one-way ANOVA on day 18 sample values. ns = non-significant; *P* > 0.05; \*\*\*\**P* < 0.001. **(F)** Daily body weight measurements for the duration of the experiment. Body weight of the study mice served as an indicator of systemic health. **(G)** Hematoxylin and Eosin (H&E) staining reveal the presence of all 3 germ layers, Ectoderm, Mesoderm, and Endoderm, indicating that all cell lines are pluripotent. Scale bar = 50 µm**. (H–M)** Teratoma sections derived from WT, KO, and NLS mESCs were analyzed by immunofluorescence staining for germ layer markers. Sections were stained with antibodies against the ectoderm marker PAX6 (**H,I**), the mesoderm marker Brachyury (TBXT) (**J,K**), and the endoderm marker SOX17 (**L,M**). (**I**) Quantification of Pax6 signal obtained in (H): n= 284 cells in WT, n= 282 cells in KO, n= 277 cells in NLS. One-way Anova, **** : p<0.0001. (**K**) Quantification of Brachyury (TBXT) signal obtained in (**J**): n= 107 cells in WT, n= 122 cells in KO, n= 132 cells in NLS. One-way Anova, **** : p<0.0001. (**M**) Quantification of Sox17 signal obtained in (H): n= 146 cells in WT, n= 167 cells in KO, n= 241 cells in NLS. One-way Anova, ns: p=0.9928 (A-B), p= 0.3612 (A-C), p= 0.3994 (B-C).

Next, we performed histological evaluation on paraffin-embedded tumor sections stained with hematoxylin and eosin (H&E). All three mESC cell lines (WT, KO, NLS) formed teratomas containing derivatives of the three germ layers. In all three cases, histological analysis revealed the formation of neural rosettes (ectoderm), cartilage and adipose tissue (mesoderm) and primitive gut-like epithelium (endoderm) (Figure 8G). To further characterize lineage specification along with the extent of ectoderm, mesoderm and endoderm tissue formation, we performed immunohistochemistry (IHC) using lineage-specific markers. WT and NLS teratomas stained positively for ectodermal (Pax6) (Figure 8H,I), mesodermal (Brachyury) (Figure 8J,K), and endodermal (Sox17) markers (Figure 8L,M), corroborating the H&E observations. Notably, in the KO condition, rare small tissue structures were formed that showed staining restricted to mesodermal and endodermal markers, with limited expression of the ectodermal marker Pax6 (Figure 8H-M).

Thus, while WT and NLS mESCs retained full tri-lineage differentiation capacity, KO mESCs displayed impaired or skewed lineage specification, indicating that nuclear β-actin plays an essential role in cell fate decision by controlling gene expression.

## Discussion

In this study, we identify nuclear β-actin as a central regulator of mESCs pluripotency that couples chromatin state to extracellular matrix (ECM)-dependent mechanics. β-actin ablation disrupted the pluripotency network, chromatin accessibility, the transcriptome and proteome, ECM composition and mechanical heterogeneity, ultimately altering lineage decisions. Re-expression of NLS-tagged β-actin rescued many of these defects, demonstrating that the nuclear actin pool is sufficient to restore key features of stem-cell identity and developmental competence.

Loss of β-actin in mESCs compromised pluripotency as evidenced by reduced alkaline phosphatase staining and OCT4, SOX2, and NANOG expression despite preserved cell numbers. Moreover, KO cells showed increased Ki-67 expression and proteomic and transcriptomic profiles that markedly diverged from those of WT cells, distinguished by the depletion of core pluripotency factors and reduced KLF5 and STAT3 levels, together with changes in structural and mechanosensing proteins. Reintroduction of NLS-tagged β-actin rescued many of the phenotypic features of WT cells, although their transcriptomic profile did not fully return to that of WT cells. This partial rescue suggests that nuclear re-expression restores essential regulatory functions while compensatory states established after β-actin loss persist. Although further analysis such as acute β-actin depletion using knockdown or inducible degron systems in mESCs will shed further light on its direct effects, our results support the conclusion that nuclear β-actin is required to maintain the molecular state underlying pluripotency.

Our integrated genomic analyses place this effect at the level of chromatin regulation. KO mESCs showed reduced accessibility at regulatory regions of Oct4, Sox2, Nanog and Klf4. Genome-wide RNA-ATAC integration showed 73% concordance among genes significant in both assays, with enrichment for stemness, ECM, mechanotransduction and early-lineage programs. Within the 18-gene ECM–pluripotency–mechanosensing program, RNA changes closely tracked promoter and proximal regulatory accessibility (Spearman r = 0.82 and 0.87). Promoter analysis revealed a shared regulatory grammar involving STAT and KLF/SP factors together with TEAD/YAP, T-box, PAX, RUNX, SMAD, AP-1 and MEF2 families. Globally, regions opening in KO cells were enriched for SRF motifs, whereas regions closing in KO cells were enriched for TEAD/YAP, STAT, KLF/SP and SMAD motifs. These patterns provide a mechanistic framework for the simultaneous loss of pluripotency transcription and activation of alternative lineage/ECM programs. In the NLS rescue, transcription recovered more strongly than accessibility and only 26% of genes significant in both layers were concordant relative to WT, indicating a gene-selective recovery with residual chromatin rewiring. EZH2 gain accompanied repression of Slit1, Hand1, Pax6, T and Cdx2 during rescue, but this selective association does not support a simple global PRC2-dependent mechanism of repression. Nuclear β-actin therefore appears to maintain a permissive regulatory architecture whose disruption redistributes transcription-factor access and selectively engages repressive pathways.

A major consequence of this chromatin rewiring is altered communication between the nucleus and the extracellular environment. KO mESCs upregulated fibronectin, collagens, tenascin-C and other ECM-related genes, accumulated fibronectin protein and a greater ECM stiffness heterogeneity. These ECM alterations were also accompanied by changes in nuclear morphology, reduced Lamin A/C and mechanosensing proteins and increased nuclear YAP1 suggesting a disrupted nucleus-ECM mechanosensing axis. The integrated RNA-ATAC data connect ECM components with *Itgav, Vcl, Piezo1, Lmna* and *Iiga1*, suggesting a mechanosensing-to-transcription axis linked to KLF5/STAT3 and lineage regulators. This is consistent with YAP/TEAD recruitment of BAF and evidence that β-actin participates in a YAP/TEAD-containing complex^84,85^. Because YAP/TEAD also regulates ECM-associated transcription^86^, nuclear β-actin may coordinate chromatin remodeling with mechanotransduction. Our AFM data further suggest that changes in ECM mechanics extend beyond mean stiffness to include increased spatial heterogeneity potentially creating distinct mechanical environments that alter the pluripotency state of KO cells.

These molecular and mechanical defects have clear functional consequences for lineage specification. KO embryoid bodies showed altered growth and increased mesoderm-associated markers, while directed neuronal differentiation failed to generate βIII-tubulin- and MAP2-positive networks. Instead, KO cultures formed contracting, cardiomyocyte-like structures and retained inappropriate mesodermal/endodermal signatures, whereas NLS cells largely recovered the neuroectodermal program. This switch is consistent with repression of neural regulatory loci, altered mechanosensing and defective ECM remodeling; dysregulated YAP-dependent signaling may further reinforce it^87^. *In vivo*, KO cells formed markedly smaller teratomas with strongly reduced ectodermal representation, whereas nuclear β-actin re-expression partially restored tumor growth and tri-lineage differentiation. Nuclear β-actin is therefore required not only for pluripotency-factor expression but for the developmental competence to execute appropriate lineage choices.

Collectively, our findings position nuclear β-actin as a key player integrating chromatin accessibility, transcription-factor networks and ECM-dependent mechanotransduction. Previous work established roles for nuclear β-actin in higher-order genome organization and ECM regulation^23,24,39^; the present study connects these functions directly to pluripotency and lineage fidelity. Although the partial NLS-tagged β-actin rescue and selective EZH2 associations indicate that additional chromatin-remodeling and compensatory mechanisms remain to be defined, the convergence of genomic, proteomic, biophysical analysis, differentiation and *in vivo* evidence supports a unified model in which nuclear β-actin preserves the transcriptional and mechanical plasticity required for cell-fate decisions.

## Materials and Methods

### Preparation of β-actin KO and NLS mESCs

Mouse ES cells were cultured on mouse embryonic fibroblasts feeder cells (MEFs) from GIBCO on Gelatin coated plates. The day before nucleofection feeder MEF cells (C57BL/6) were seeded in a 24well format (1.2x10e5cells/well). Cells were nucleofected using the Amaxa 4D-Nucleofector system with program CM-150. For the RNP complex 30pmol of spCas9 nuclease (V3 from IDT) was combined with 60pmol of sgRNA. This reaction was incubated in Cas9 nuclease reaction buffer (IDT) in DEPC- treated water to a total of 3µl, for 20min on RT. HDR templates were added shortly before incubation time was over. For mActinFRT KnockIn, 1µg dsDNA HDR was used, for mActinNLS as well as Flippase Knockin, 2µg HDR template were added (concentrations of plasmids were adjusted to 1 µg/µl or 2µg/µl respectively). In the meantime, cells were detached with TrypLE Express, counted and spun down at 200g for 5min. 5e4 mESC per condition were resuspended in 20µl P3 Primary 4D-Nucleofector Master mix (16.4µl P3 solution and 3.6µl supplement). 18µl of cell suspension were added to 4µl of RNP/HDR template mixture and 20µl were transferred to one well of a Nucleocuvette™ Strip. Immediately after nucleofection, cells were transferred in pre-warmed 24 well plates containing MEF feeders, ESC medium and HDR Enhancer (IDT) in a concentration of 0.72µl/ml). Medium was changed the next day and cells were grown until 60% confluency. Cell population was then expanded until aliquots could be frozen. For this, cells were detached, spun down at 200g for 5min and re-suspended in Bambanker hRM (GC Lymphotec Inc.) with a concentration of 10e6 cells/ml. Cells were frozen 1°C per min and stored in liquid Nitrogen.

For Genotyping, edited pools were cultured feeder free for 3 passages before cells were pelleted and lysed for gDNA extractin according to manufacturer’s protocol (Machery Nagel NucleoSpin Tissue Kit). After confirmation of gene editing by PCR, cells were seeded as single cell colonies. For this, p100 plates were Gelatin coated and 3x10e6 MEFs were seeded 24h prior to seeding 500 edited cells. The medium was changed every other day and cells were grown until colonies formed. These colonies were then mechanically picked by scraping them off with a P200 tip under the microscope and transferred onto 96 well plates coated with MEFs.

These 96 well plates were then replica split onto two 96 well plates. While one was kept in culture, the other plate was used to genotype the clones and to confirm successful gene editing. Cells were lysed with QuickExtract DNA Extraction Solution (Biosearch Technologies) and Cell lysate was used in a genotyping PCR. Edited clones were further expanded and stocks were frozen.

Around 10e6 cells were lysed and genomic DNA was extracted according to manufacturer’s protocol (Machery Nagel NucleoSpin Tissue Kit). Cell Clones were checked for off-target effects of the CRISPR editing by Sanger sequencing. For this the 500bp around the 10 most likely off-targets predicted by the CRISPOR algorithm per used sgRNA were amplified by PCR and sequenced^88^.

### Immunofluorescence staining and quantification

WT, KO and NLS mESCs were seeded onto 0.1% gelatin coated coverslips and at the proper confluency fixed with 4% Paraformaldehyde for 20min at RT. Cells were washed 3 times with PBS prior to incubation with PBS-Tween (0.5%) for 1h at RT. Coverslips were again washed 3 times with PBS. Primary antibodies were diluted in PBS-BSA (1%) and coverslips were incubated with primary antibody solution for 2h at RT or at 4°C overnight. Cells were washed with PBS 3 times and then incubated with secondary antibodies diluted in PBS-BSA (1%) 1h at RT in the dark. Cells were again washed with PBS 3 times. DAPI, diluted 1:500 in PBS was applied for 10min at RT in the dark prior to 3 washes with PBS. Coverslips were then mounted on glass slides using a mounting medium. Various staining were used: Oct4 (ab18976), Sox2(ab97959), Ki67, Fn(ab2413), GATA4 (sc-25310), BIIITubulin (ab78078), Map2 (ab32454), H3K9me3 (ab8898), β-actin (Sigma-Aldrich A2228), Lamin A/C (SAB4200236), Lap2(ab223768), Yap1(sc101199), Phalloidin(A22287). Images were acquired as z stacks with a step size of 1 µm for each position. Using ImageJ software, the chosen z-plane of the nucleus was used to make a mask (DAPI channel) and the phalloidin staining was used to generate the cell/cytosol mask. The mean intensity of each staining was quantified using the DAPI mask for nuclear measurements or the Phalloidin mask for cell measurements and was plotted using Prism 11. For the Map2, beta tubulin intensity: the total intensity was calculated and divided by the total number of DAPI positive nuclei and this intensity was plotted. The percentage of Map2 positive and beta tubulin positive cells was plotted separately. Images were taken using dragonfly Endor spinning desk microscope, with objectives of either 40x or 60x.

### Maintenance of mouse embryonic stem cells

WT, KO, and NLS E14 mESCs were cultured in P60 culture dishes that were coated with 3 mL of 0.1% gelatin solution and incubated at 37 °C for 15 min. mESC medium was pre-warmed to 37 °C. The gelatin solution was then replaced with 5 mL of mESC medium. A total of 5.0 × 10□ cells were seeded for each condition. Cultures were maintained at 37 °C and 5% CO□ for 2 days after seeding. Once cultures reached 70–90% confluency, cells were processed for downstream experiments or passaged. The mESC medium was composed of 410mL DMEM, 75mL of FBS, 5mL of 100x Sodium Pyruvate, 5mL of 100x NEAA, 0.5mL of 1000x β-mercaptoethanol, and 20ng/mL of LIF. Medium was filtered with 0.22µm filter before use.

### Alkaline Phosphatase assay

Alkaline phosphatase (AP) staining was performed using the Purple-Color AP Staining Kit (System Biosciences, Palo Alto, CA, USA; cat. no. AP100P-1) according to the manufacturer’s instructions. A total of 10,000 cells were seeded per well in 96-well plates and cultured for 24 h at 37 °C with 5% CO before staining. Cells were washed once with 1× PBS, fixed with Fix Solution for 5 minutes at room temperature, washed again with 1× PBS, incubated with AP Stain in the dark for 20 minutes, and washed with 1× PBS to stop the reaction, after which cells were imaged by bright-field microscopy.

### qRT-PCR

Total RNA was isolated using RNAzol (Sigma-Aldrich, Saint Louis, MO, USA; cat. no. R4533) following the manufacturer’s protocol. For cDNA synthesis, 500 ng total RNA per reaction was reverse transcribed using the RevertAid First Strand cDNA Synthesis Kit (Thermo Fisher Scientific; cat. no. K1622). The resulting cDNA was diluted and used as template for quantitative real-time PCR (qPCR) with SYBR Green qPCR Mix (Thermo Fisher Scientific; cat. no. 4309155) on a StepOnePlus Real-Time PCR System (Thermo Fisher Scientific; cat. no. 4376600). Expression levels of target genes were normalized to the housekeeping gene GAPDH. Primer sequences used for qPCR are listed in Supplementary Table 9.

### ATAC-seq

ATAC-seq was performed using the Zymo-Seq™ ATAC Library Kit (Zymo Research, Cat. #D5458), following the manufacturer’s protocol. Mouse embryonic stem cells (mESCs) representing wild-type (WT), β-actin knockout (KO), and nuclear localization signal β-actin rescue (NLS) were cultured under standard conditions. For each genotype, three biological replicates were processed, with 50,000 viable cells used per replicate, as determined by trypan blue exclusion and automated cell counting.

Cells were harvested at the undifferentiated state, washed with PBS, and resuspended in ATAC-S Buffer. After adding cold ATAC Lysis Buffer, samples were incubated on ice to lyse cells and isolate nuclei. The lysed samples were washed with ATAC Wash Buffer and centrifuged at 1,000 × g for 10 minutes to pellet nuclei. The nuclei were then incubated with Pre-Tagmentation Buffer and Tn5 transposase at 37°C for 30 minutes in a thermoshaker at 1,000 rpm.

Following transposition, DNA was purified using Zymo-Spin IC Columns and eluted in DNA Elution Buffer. Libraries were then amplified using the kit-supplied ATAC Library PCR Mix and unique dual index (UDI) primers (one per replicate). PCR was carried out for 10 cycles, optimized for cell line samples. Libraries were purified again using spin columns and quantified using Qubit and TapeStation (Agilent) to verify size distribution and absence of adapter dimers. The final ATAC-seq libraries were sequenced using the Illumina HiSeq 2500 platform (performed at the NYU Abu Dhabi Sequencing Center).

### ATAC-seq preprocessing

Raw sequencing reads were processed to remove low-quality bases and adapter contamination using Trimmomatic v0.64, with quality control performed via **FastQC** (http://www.bioinformatics.babraham.ac.uk/projects/fastqc). Trimming parameters included: trimmomatic_adapter. fa:2:30:10, TRAILING:3, LEADING:3, SLIDINGWINDOW:4:15, and MINLEN:36. High-quality paired-end reads were subsequently aligned to the mouse reference genome (GRCm38) using BWA-MEM v0.65. Resulting alignments were cleaned, sorted, and deduplicated (removal of PCR and optical duplicates) using Picard Tools (http://broadinstitute.github.io/picard). To visualize chromatin accessibility, BigWig coverage tracks were generated from the cleaned BAM files using deepTools (bamCoverage) with the following parameters: --binSize 10, --extendReads, --ignoreDuplicates, and --normalizeUsing RPKM. ENCODE blacklisted regions were excluded. Replicate BigWig files were averaged using the bigwigCompare tool with the --operation mean option. Aggregated signal over genomic regions of interest was calculated and visualized using the computeMatrix function in deepTools.

### RNA-seq Analysis

NASQAR, a web-based platform for High-throughput sequencing data analysis and visualization was used to do differential expression transcriptional analysis. Through NASQAR, DEseq2 was used to perform pairwise differential expression comparisons between WT and KO mESCs and generate a Volcano plot, Distance heatmap and a PCA plot. GO analysis was also done on NASQAR to generate GO plots for Cellular Components and Molecular Functions. Heatmaps were generated in R (v4.3.0) using the pheatmap package to visualize the expression patterns of selected gene sets across wild-type (WT), β-actin knockout (KO), and nuclear-localized β-actin (NLS) mESCs at Day 0. Raw gene expression matrices from bulk RNA-seq were first imported using the readr package. Gene sets related to specific biological functions (e.g., stemness, lineage commitment, extracellular matrix) were defined manually, and the corresponding expression values were extracted from the full dataset using the dplyr package. Replicates for each condition (WT_1–3, KO_1–3, NLS_1–3) were retained as individual columns or averaged using rowMeans to create representative condition-specific profiles when required. Expression values were transformed into a numeric matrix. The matrix was then scaled row-wise (z-score) to allow for comparison of relative expression across samples. Heatmaps were plotted using pheatmap with hierarchical clustering applied to genes (rows), while preserving the experimental condition order for samples (columns). A custom diverging color palette (navy–white–firebrick3) was used to depict low to high expression.

### ATAC-Seq – RNA-Seq integration

Accessibility (ATAC-seq) and expression (RNA-seq) were analysed separately with DESeq2. ATAC peaks were assigned to genes with HOMER (mm10), and each gene was represented by its most significant peak (lowest adjusted p-value). A gene was called significant in a layer at adjusted p < 0.05 with |log FC| ≥ 0.25 (RNA) or ≥ 0.1 (ATAC). Genes significant in both layers were classified by fold-change sign as concordant (Activated, open + up; Repressed, closed + down), discordant (Primed or Post-transcriptional), or Stable; the KO-vs-NLS contrast had none and was excluded.

### Preparation of protein digests, TMT labeling, and high pH reverse-phase fractionation

Advanced proteomic analysis of cytoplasmic and nuclear fractions was performed on mouse celllysate proteins from wild type (n = 3), knockout (n = 3), and NLS (n = 3) groups. Briefly, 100 µg aliquots of each sample were reduced (20 mM DTT, 70 °C, 30 min) and S-alkylated (50 mM IAA, 1 h in the dark). Excess IAA was quenched with 20 mM DTT. Proteins were precipitated with 15% trichloroacetic acid (TCA) overnight at 4 °C, pelleted by centrifugation at 16,000 × g for 15 min at 4 °C, washed with ice-cold acetone, air-dried, and resuspended in 50 mM ammonium bicarbonate. Protein digestion was performed using an MS-grade trypsin/Lys-C mix at an enzyme-to-protein ratio of 1:50 (w/w) for 24 h at 37 °C, and the reaction was quenched with 1 µL formic acid. The resulting peptides were labeled with TMTpro 16-plex reagents according to the manufacturer’s instructions (Thermo Fisher Scientific), and the labeled samples were pooled. The pooled sample was fractionated offline using a high-pH reversed-phase peptide fractionation kit according to the manufacturer’s protocol (Pierce). A total of 11 fractions were collected, dried, and reconstituted in 1% formic acid prior to nano-LC-MS/MS analysis on an Orbitrap Fusion Lumos system (Thermo Scientific).

### Liquid chromatography tandem mass spectrometry

LC-MS/MS was performed on a fully automated Ultimate 3000 nano-LC system coupled online to an Orbitrap Fusion Lumos mass spectrometer. Mobile phases consisted of 0.1% formic acid in water (A) and 0.1% formic acid in 95% acetonitrile (B). Peptides reconstituted in solvent A were loaded onto a C18 nano-trap column (Acclaim PepMap, 250 mm × 75 µm, Thermo Scientific), washed with 0.5% acetonitrile and 0.1% formic acid, and separated using a 150 min linear gradient. The gradient was as follows: 1% B for 5 min, 1–5% B for 1 min, 5–18% B for 59 min, 18–32% B for 30 min, 32–45% B for 10 min, and 45–90% B for 1 min. A 12 min wash at 90% B was used to minimize carryover, followed by 20 min equilibration at 1% B. The flow rate was maintained at 350 nL/min. Peptides were ionized by nano-electrospray at 2.0 kV, and the capillary temperature was set to 320 °C.

Spectra were acquired on an Orbitrap Fusion Lumos system operated in data-dependent acquisition mode using an SPS-MS3 workflow. MS1 spectra were collected in the Orbitrap at a resolution of 120,000, with an AGC target of 200,000 and a maximum injection time of 50 ms. Precursors with charge states of 2–7 and intensity greater than 5,000 were selected, and monoisotopic precursor selection was set to peptide. Previously analyzed precursors were excluded using dynamic exclusion of 60 s with a ±10 ppm window. MS2 precursors were isolated with a quadrupole isolation window of 0.7 m/z, and MS2 spectra were acquired in the ion trap with an AGC target of 10,000, a maximum injection time of 70 ms, and CID collision energy of 35%. For MS3 analysis, the Orbitrap was operated at a resolution of 50,000, with an AGC target of 50,000 and a maximum injection time of 105 ms. MS3 fragmentation was performed by HCD at a normalized collision energy of 60% to maximize TMT reporter ion yield, and synchronous precursor selection was enabled to include up to 10 MS2 fragment ions in the MS3 scan.

### MS Data processing

Raw MS data were searched against the mouse UniProt database, and protein groupings were assigned using PD 3.2. These groupings were then refined with a custom script to select the best-annotated master proteins. Peptide matches mapping to multiple proteins were resolved by reference to the master protein list generated during the total protein analysis.

### MS Data analysis and protein selection

Multivariate analyses, including principal component analysis (PCA) and partial least squares-discriminant analysis (PLS-DA), were performed using SIMCA 18 (Sartorius, Umeå, Sweden). The data were mean-centered and Pareto scaled prior to analysis. Model robustness was assessed by cross-validation using the leave-one-out approach, and model performance was evaluated based on the fit parameters R2 X and R2 Y as well as the predictive ability Q2. A pairwise comparative PLS-DA was also performed and used for significance testing across protein features in the dataset. Significantly altered proteins in KO and NLS extracts relative to wild type were selected based on variable importance in projection (VIP) scores from the PLS-DA model and adjusted p-values from Student’s t-test with false discovery rate correction using the Benjamini–Hochberg method. Proteins with VIP > 1.0 were considered significantly altered.

### Protein enrichment and pathway analysis

Proteins with VIP > 1.0 in the WT/KO, WT/NLS, and KO/NLS comparisons were analyzed using Metascape (https://metascape.org/gp) for pathway enrichment analysis. Gene Ontology (GO) and pathway enrichment analyses were performed to identify biologically relevant pathways associated with significantly up-regulated and down-regulated proteins. The enrichment analysis included WikiPathways, KEGG Pathway, GO Biological Processes, and Reactome Gene Sets. Terms were selected based on a p-value < 0.01, a minimum count of 22, and an enrichment factor > 3.69. The top 20 enrichment terms were visualized as a bubble plot colored by rescaled -log10(q value).

### EZH2 ChIP-Seq

ChIP-Seq for Ezh2 were performed as described in Mahmood et al (2021). Mouse Embryonic fibroblasts (three biological replicates per condition and controls) were crosslinked with 1% formaldehyde (Sigma-Aldrich, F8775) for 10 min, and the reaction was quenched with 0.125 M glycine for 5 min. Cells were initially lysed in LB1 buffer containing 50 mM HEPES- KOH (pH 7.5), 10 mM NaCl, 1 mM EDTA, 10% glycerol, 0.5% NP-40, and 0.25% Triton X- 100. Nuclei were collected by centrifugation and washed with LB2 buffer containing 10 mM Tris-HCl (pH 8.0), 200 mM NaCl, 1 mM EDTA, and 0.5 mM EGTA. Nuclear lysis was then performed using LB3 buffer containing 10 mM Tris-HCl (pH 8.0), 100 mM NaCl, 1 mM EDTA, 0.5 mM EGTA, 0.1% sodium deoxycholate, and 0.5% N-lauroylsarcosine. Chromatin was fragmented using a Qsonica sonicator for four 3-min cycles at 80% amplitude. Fragmentation was assessed by electrophoresis on a 0.8% agarose gel. For each immunoprecipitation, 100 µg of fragmented chromatin was incubated with 5 µg of the relevant antibody: EZH2 (D2C9) XP Rabbit monoclonal antibody (Cell Signaling Technology). Antibody–chromatin complexes were captured using Pierce Protein A/G Magnetic Beads (Thermo Fisher Scientific). The bead-bound immunocomplexes were washed twice with low-salt wash buffer containing 0.1% SDS, 2 mM EDTA, 1% Triton X-100, 20 mM Tris-HCl (pH 8.0), and 150 mM NaCl, followed by two washes with high-salt buffer containing the same components with 500 mM NaCl. Chromatin was eluted in buffer containing 50 mM Tris-HCl (pH 8.0), 10 mM EDTA, and 1% SDS. Reverse crosslinking was achieved overnight at 65°C. Samples were subsequently treated RNase A for 30 min at 37°C. This was followed by the addition of 4 µL of 0.5 M EDTA, 8 µL of 1 M Tris-HCl, and 1 µL of proteinase K at 20 mg/mL, corresponding to a final concentration of approximately 0.2 mg/mL. Samples were incubated for 2 h at 42°C. DNA was purified using phenol, chloroform, and isoamyl alcohol and used for ChIP-seq libraries were prepared using the TruSeq NanoDNA Library Preparation Kit (Illumina, San Diego, CA, USA) and sequenced on a NextSeq 550 platform at the NYU Abu Dhabi Sequencing Center.

### Bioinformatics methods

Day-0 wild-type (WT), β-actin-knockout (KO) and nuclear β-actin re-expression (NLS) samples were analysed. The RNA-seq analysis comprised four biological replicates per condition (12 libraries; GEO GSE327304), the ATAC-seq analysis comprised four WT, three KO and two NLS biological replicates (nine libraries; GEO GSE327117), and the EZH2 ChIP-seq (GEO GSE346145) analysis comprised three replicate libraries per condition together with one condition-matched input library for each condition. Unless otherwise stated, genomic analyses used the mouse GRCm38/mm10 assembly, GENCODE mouse release M25 gene annotation and chromosomes 1-19, X and Y. Ensembl gene identifiers were reduced to their stable identifier by removing version suffixes before cross-assay joins. Three contrasts were evaluated throughout the gene-level analysis: KO minus WT, NLS minus KO and NLS minus WT. Positive log2 fold changes therefore indicate higher abundance, accessibility or EZH2 signal in the first-named condition. KO minus WT quantified the knockout state, NLS minus KO quantified the direct response to nuclear β-actin re-expression, and NLS minus WT quantified the residual difference from WT. See below points 1-12 for detailed bioinformatics procedures.

#### 1. RNA-seq differential-expression analysis

Gene-level raw counts for the day-0 samples were reconstructed from the GSE327304 raw-count archive. Genes were retained when they had at least 10 counts in at least four libraries. Differential expression was analysed in DESeq2 using the design ∼ replicate + condition, where replicate was the R1-R4 blocking factor shared across conditions. DESeq2 Wald contrasts were calculated for KO minus WT, NLS minus KO and NLS minus WT. Unshrunken log2 fold changes were used for integration and visualization, and P values were adjusted within each contrast by the Benjamini-Hochberg method.

#### 2. ATAC-seq peak reconstruction and differential accessibility

Replicate-level ATAC-seq BAM files were recovered from the source merged BAMs using the preserved read-group tags. Properly paired alignments with mapping quality at least 30 were retained, while reads flagged as unmapped, mate-unmapped, secondary, failed quality control or duplicate were removed (samtools view -f 2 -F 1804 -q 30). Per-replicate peaks were called with MACS3 in paired-end mode (callpeak -f BAMPE -g mm -q 0.01 --keep-dup all). Peak calls from all nine libraries were restricted to chromosomes 1-19, X and Y, sorted, merged and assigned unique identifiers, producing a consensus universe of 162,388 intervals. One properly paired fragment per molecule was counted over each interval. Differential accessibility was analysed with edgeR. Initial trimmed-mean-of-M-values (TMM) factors were calculated, filterByExpr was applied with min.count=10, and TMM normalization was recalculated after filtering, retaining 161,141 intervals. A no-intercept condition design was fitted with robust dispersion estimation and robust quasi-likelihood generalized linear models. Quasi-likelihood F tests were performed for KO minus WT, NLS minus KO and NLS minus WT, and P values were adjusted by the Benjamini-Hochberg method. All nine libraries were used for the primary ATAC analysis.

#### 3. EZH2 ChIP-seq processing, peak definition and quantification

Paired-end reads were processed with fastp v0.24.0 and aligned to mm10 with Bowtie2 v2.5.4 using the very-sensitive preset, a maximum fragment length of 2,000 bp, and the options --no-mixed, --no-discordant and --dovetail. Proper pairs with mapping quality at least 30 were retained (samtools view -f 2 -F 2828 -q 30). Mate information was repaired with samtools fixmate, PCR duplicates were removed with samtools markdup, and alignments on mitochondrial, non-primary or ENCODE mm10 v2 blacklist regions were excluded. EZH2 broad peaks were called separately for each replicate library with MACS3 v3.0.3 in BAMPE mode using the corresponding condition-matched input (-q 0.01 --broad --broad-cutoff 0.1 --keep-dup all). For each condition, regions present in at least two of the three replicate peak sets were merged and intersected with the peak set obtained from pooled condition-level ChIP libraries. The WT, KO and NLS consensus sets were then merged into a non-overlapping master universe of 5,339 EZH2 intervals. Paired fragments were counted over the master intervals with featureCounts/Subread v2.0.8 using -p --countReadPairs -B -C; 5,338 intervals with a summed count of at least 10 across the nine EZH2 libraries entered differential analysis. ChIP-seq quality control included fastp and alignment summaries, duplicate fractions and estimated library sizes from samtools markdup, deepTools fingerprints, Pearson and Spearman correlations and principal-component analysis from 10-kb genomic bins, replicate- and consensus-peak fractions of reads in peaks, and pairwise Jaccard similarity of replicate peak sets. Strand cross-correlation was calculated with phantompeakqualtools. The primary EZH2 normalization used relative-log-expression (RLE) size factors estimated from blacklist-filtered 10-kb genomic-bin counts. Bin counts were generated with deepTools v3.5.6 for the nine EZH2 ChIP libraries. Bins with zero total count across all nine libraries were removed, leaving a 247,633-bin matrix from which the RLE factors were estimated. These externally estimated size factors were assigned to a DESeq2 model with design ∼ condition. Wald contrasts were calculated for KO minus WT, NLS minus KO and NLS minus WT, with Benjamini-Hochberg adjustment within each normalization-specific contrast. Peak-derived RLE, peak-derived TMM, background-derived TMM and full-library normalization were evaluated as normalization comparisons. Full-library size factors were the final paired-fragment library sizes, calculated as final filtered aligned reads divided by two and geometrically centred across the nine libraries. Because no exogenous spike-in was included, EZH2 fold changes represent relative changes on the stated normalization scale.

#### 4. Gene-centered RNA, ATAC and EZH2 integration

Protein-coding genes were obtained from GENCODE M25. Promoters were defined in a strand-aware manner from 1,000 bp upstream to 100 bp downstream of the gene-record transcription start site (TSS). Two nested peak-to-gene models were used. The promoter model included consensus intervals that overlapped the promoter, whereas the proximal model included every consensus interval overlapping the TSS-centred plus or minus 20-kb window. The latter therefore contained promoter and non-promoter proximal intervals. ATAC and EZH2 intervals were assigned to genes independently, without requiring an ATAC-EZH2 overlap. An interval was counted once per gene and model but could contribute to more than one gene when gene-centred windows overlapped. Raw ATAC or EZH2 interval counts were summed by gene and sample within each regional model. ATAC gene aggregates were analysed with edgeR using the original library sizes and TMM factors estimated from the complete consensus-peak universe. filterByExpr was applied with min.count=10, retaining the complete-universe library sizes, followed by robust dispersion estimation and quasi-likelihood testing. EZH2 gene aggregates with at least 10 total fragments were analysed with DESeq2 using the background-RLE size factors described above. Assay-specific P values were adjusted by the Benjamini-Hochberg method within each regional model and contrast. RNA estimates were joined to ATAC and EZH2 estimates by stable Ensembl gene identifier and contrast. The promoter ATAC and EZH2 values displayed for the selected genes in Figure 4 were obtained from this called-peak aggregation model (promoter_called); the direct promoter-window counts described below were used for the separate genome-wide RNA-EZH2 promoter correspondence analysis.

Figure 4 used a prespecified 18-gene display set divided into extracellular-matrix or niche genes (Col1a1, Col1a2, Hapln1, Tnc, Slit1, Col4a6 and Col4a5), pluripotency or lineage regulators (Stat3, Klf5, Hand1, Pax6, T and Cdx2), and mechanosensing or adhesion genes (Itgav, Vcl, Piezo1, Lmna and Itga1). For each gene and contrast, one RNA effect estimate was displayed alongside regional ATAC or EZH2 estimates from the promoter and aggregate plus or minus 20-kb called-peak models. Asterisks denoted the assay-specific Benjamini-Hochberg FDR threshold of 0.05. Within-set RNA-ATAC and RNA-EZH2 associations were summarized descriptively by complete-case Spearman correlations within the fixed 18-gene set for each contrast and regional model; correlation P values were adjusted across the 12 assay-pair, contrast and regional-model combinations and were not used to define figure support. Direct inverse RNA-EZH2 correspondences were defined as NLS-minus-KO effects with FDR below 0.05 in both assays and opposite RNA and EZH2 fold-change signs; correspondences highlighted in the figure were required in both regional models.

#### 5. Genome-wide cross-assay correspondence analyses

For the direct all-gene promoter analysis, paired EZH2 fragments were also counted over the strand-aware TSS-minus-1-kb to TSS-plus-100-bp intervals with featureCounts using -p --countReadPairs -B -C -O. The -O option permitted a fragment overlapping more than one promoter to contribute to each applicable promoter. Promoters with at least 10 total counts were analyzed with DESeq2 using the background-RLE factors and three contrasts defined above. An RNA-EZH2 promoter rescue correspondence required both assays to pass FDR below 0.05 in KO minus WT and NLS minus KO, the NLS-minus-KO effect to oppose the KO-minus-WT effect within each assay, and the absolute NLS-minus-WT residual to be smaller than the absolute KO-minus-WT effect. RNA and EZH2 were additionally required to change in opposite directions in both the knockout and direct re-expression steps. FDR was controlled within each assay; the cross-assay intersection was applied as a deterministic correspondence rule. KO-state EZH2-ATAC-RNA triads were evaluated across four gene-assignment models: the called-peak promoter model, the aggregate TSS plus or minus 20-kb model, a strand-aware mouse Inferelator window extending 50 kb upstream and 2 kb downstream of the TSS while including the complete gene body and constraining boundaries at adjacent gene bodies, and assignment of each peak to its nearest protein-coding TSS without a distance cap. Within a model, a triad required FDR below 0.05 in all three assays for KO minus WT, an EZH2 effect opposite to both the ATAC and RNA effects, and ATAC and RNA effects in the same direction. The primary cross-model definition required this rule in at least three of the four models. Three-assay directional movement was evaluated by requiring the direct NLS-minus-KO effect to oppose the KO-minus-WT effect and the absolute NLS-minus-WT residual to be smaller than the absolute KO-minus-WT effect separately for each assay.

#### 6. Aggregate ATAC movement toward WT

Aggregate ATAC movement was assessed separately for the promoter and plus or minus 20-kb gene models. Genes were first selected at FDR below 0.05 in the KO-minus-WT contrast. A selected gene was classified as moving toward WT when the NLS-minus-KO effect opposed the KO-minus-WT effect and the absolute NLS-minus-WT residual was smaller than the absolute KO-minus-WT effect. The observed directional fraction was tested by an exact label-randomization procedure that held WT labels fixed, reassigned the three KO and two NLS labels among the five non-WT libraries, and repeated model fitting, KO-minus-WT selection and directional scoring over the observed filterByExpr-retained gene universe for all ten possible assignments. A one-sided P value was calculated as the proportion of assignments with a directional fraction at least as large as the observed fraction. A 95% interval for the observed fraction was obtained from 5,000 chromosome-block bootstrap samples.

A model-based bootstrap was also performed under a no-rescue null in which NLS expected counts were constrained to the fitted KO expectation while retaining sample-specific library offsets and tagwise negative-binomial dispersions. For each of 2,000 simulations per regional model (seed 20260729), counts were generated from the fitted negative-binomial distribution and filterByExpr, dispersion estimation, the three-condition quasi-likelihood fit, KO-minus-WT FDR selection and directional scoring were repeated. The bootstrap P value used the plus-one estimator, (b+1)/(B+1), where b was the number of simulated directional fractions at least as large as the observed value and B was the number of valid simulations.

#### 7. Promoter motif occurrence in the 18-gene set

Sequences for the strand-aware TSS-minus-1-kb to TSS-plus-100-bp promoters were extracted from mm10. JASPAR 2024 CORE vertebrate non-redundant position-frequency matrices were restricted to 12 prespecified families: AP-1, CDX, HAND, KLF/SP, MEF2, PAX, RUNX, SMAD, SRF, STAT, T-box and TEAD/YAP. A pseudocount of 1e-4 was added to each motif matrix. Nucleotide-background frequencies were estimated from the target promoter sequences after adding a one-count pseudocount to each base, and motif probabilities were converted to log2-odds position-weight matrices. Both strands were scanned. A motif model was considered present when its maximum relative PWM score was at least 0.90. Each heat-map cell reports the number of distinct motif models in the family with at least one qualifying site in that promoter; the accompanying coverage bar reports the number of the 18 promoters with at least one qualifying model. Families supported in the KO-loss ATAC analysis were highlighted using the same colour annotation across panels.

#### 8. Genome-wide ATAC motif-family analysis

The 161,141 ATAC intervals retained by the primary all-nine-library edgeR analysis were scanned for 141 HOCOMOCO v14 CORE motifs assigned to the same 12 TF families. Motifs were converted to log2-odds PWMs using a pseudocount of 1e-4 and nucleotide frequencies estimated over the complete ATAC peak-sequence universe. Both strands were scanned, and motif presence was defined by a maximum relative PWM score of at least 0.90. Family presence in a peak was defined as the presence of at least one constituent motif model. For KO minus WT and NLS minus WT, differential peaks were defined by FDR below 0.05 and separated by fold-change sign into gain and loss sets. Reference peaks from the corresponding contrast were required to have FDR of at least 0.50 and an absolute log2 fold change no greater than 0.25. Differential peaks were matched without replacement to reference peaks using standardized log2(sequence length + 1), GC fraction, edgeR logCPM and log10(nearest-protein-coding-TSS distance + 1). Greedy nearest-neighbour matching used a Euclidean-distance caliper of 2.5 in this four-dimensional z-standardized covariate space and prioritized differential peaks by absolute log2 fold change. For each family, motif presence was compared within matched peak pairs. A one-sided exact binomial test on discordant pairs, equivalent to an exact one-sided McNemar test, assessed enrichment in the differential set. Effect sizes were reported as log2 matched odds ratios with a 0.5 Haldane correction, and P values were adjusted across the 12 families within each peak set. The one-sided test evaluated enrichment only; negative effect estimates were displayed as directional estimates and were not interpreted as tests of motif depletion. In a second enrichment specification, non-set intervals were selected from five-quantile strata of log2(sequence length + 1), GC fraction, edgeR logCPM and log10(nearest-protein-coding-TSS distance + 1). Progressively reduced matching keys were used when no candidate was available in the complete stratum, and at most 20 controls were retained per occupied stratum. Family presence was compared by a one-sided Fisher exact test with Benjamini-Hochberg adjustment across the 12 families within each peak set. Main-figure support required positive enrichment at FDR below 0.05 in both the uniquely paired stable-peak and covariate-binned analyses.

#### 9. Genome-wide ATAC motif accessibility

Motif-associated accessibility was quantified independently of EZH2 with chromVAR v1.28.0 using all nine ATAC libraries, all 161,141 edgeR-retained intervals and the 141-motif HOCOMOCO panel. GC fraction was supplied as the sequence-bias covariate, and 50 chromVAR background iterations were used to match peaks jointly on GC content and average accessibility (set.seed(260714)). Motif deviation z-scores were analysed with limma v3.62.1 using a no-intercept condition design, contrasts for KO minus WT, NLS minus KO and NLS minus WT, and robust empirical-Bayes moderation. P values were adjusted across all 141 motif models within each contrast.

#### 10. ATAC footprint analysis

TF footprinting was performed as a replicate-aware analysis using differential peak definitions from the all-nine-library edgeR model and the uniquely matched reference peaks described above. Footprint tracks were generated independently from fastp-trimmed, BWA-MEM-aligned ATAC-seq FASTQs. Properly paired MAPQ-30 alignments were retained (samtools view -f 2 -F 2828 -q 30), and duplicates were removed with Picard. Seven higher-depth libraries were used for this analysis (WT1, WT2, WT4, KO3, KO4, NLS1 and NLS2); WT3 and KO2 remained in the all-nine differential peak definitions but were not used to generate the seven-library tracks. The seven BAM files were randomly subsampled toward 57,875,123 alignments per library, with a maximum observed deviation of 0.011% from the target depth. TOBIAS ATACorrect was applied using the mm10 genome and the ATAC consensus peak set, followed by ScoreBigwig over the union of differential and matched-reference regions. Exact HOCOMOCO motif instances were defined at a relative PWM threshold of 0.95. Omitted positions within the validated scoring mask were restored as zero, and scores were averaged over the complete motif width. Motif-site means were averaged within motif and peak, after which motif means were averaged within TF family and peak. Reciprocal target-reference peak pairs were retained for a feature only when both members contained that motif or family, and a motif or family context was estimable only when at least 20 complete reciprocal pairs were available. Biological libraries, rather than motif sites or peaks, were used as the inferential unit. KO-minus-WT, NLS-minus-KO and NLS-minus-WT effects were tested using exhaustive two-sided label permutations and Welch tests; Benjamini-Hochberg adjustment was applied separately within feature type, differential peak set, contrast and score definition. For peak sets selected in KO minus WT, directional rescue required the NLS-minus-KO effect to oppose the KO-minus-WT effect and the absolute NLS-minus-WT residual to be smaller than the absolute KO-minus-WT effect. As a lower-depth sensitivity analysis, all nine ATAC libraries were deterministically equalized by keyed read-name selection to exactly 7,350,953 complete pairs (14,701,906 alignments) per library. The same differential peak sets, motif sites, reciprocal peak pairs, sparse-zero score reconstruction and replicate-level tests were then applied. For the seven libraries common to both analyses, motif- and family-level footprint estimates were also compared between the higher- and lower-depth tracks.

#### 11. EZH2 interval motif-burden analysis

Sequences for all 5,338 tested EZH2 intervals were extracted from the GRCm38 top-level reference FASTA. The 141 HOCOMOCO v14 motifs and 12-family assignment used for the ATAC analysis were scanned directly over the EZH2 interval universe with the same pseudocount, bidirectional scanning and relative PWM threshold of 0.90. The nucleotide background was estimated independently from the complete EZH2 interval-sequence universe. For each interval and family, motif burden was defined as the fraction of the family’s motif models present in that interval. GC fraction and CpG observed-to-expected ratio were calculated from the interval sequence. The promoter covariate indicated overlap with the pipeline-defined, strand-aware region extending 2 kb upstream and 500 bp downstream of the TSS of any GENCODE vM25 annotated gene; this covariate was separate from the protein-coding TSS-minus-1-kb to TSS-plus-100-bp promoter used for gene linking. Motif-burden models were fitted separately for KO minus WT and NLS minus WT, and separately for intervals with relatively higher or lower EZH2 signal. Direction-specific target intervals had DESeq2 FDR below 0.05 and a fold-change sign matching the tested direction. The primary reference group comprised intervals with DESeq2 FDR of at least 0.10; opposite-direction significant intervals and intervals between the target and reference thresholds were not included in that model. A minimum of 20 target and 20 reference intervals was required. For each family, a binomial generalized linear model related target-group membership to family motif burden, log1p(baseMean), log1p(interval width), GC fraction, CpG observed-to-expected ratio and promoter status. All continuous predictors were z-standardized; promoter status was binary. HC3 heteroscedasticity-consistent standard errors and 95% confidence intervals were used, and effects were reported as log2 odds ratios per standard-deviation increase in motif burden. Benjamini-Hochberg adjustment used the 12 planned families as a fixed denominator within each normalization, contrast, direction and reference definition. The primary analysis used background-RLE normalization and the FDR-at-least-0.10 reference. The figure support symbol additionally required preservation of the coefficient sign with a stricter background-RLE reference of FDR at least 0.50, peak-RLE normalization and full-library normalization. As a continuous analysis, EZH2 log2 fold change was modelled across all tested intervals using the same predictor and covariates. Ordinary least-squares models with HC3 standard errors and inverse-variance weighted least-squares models with weights 1/lfcSE^2 and HC3 standard errors were fitted separately. P values were adjusted across the 12 families within each normalization, contrast and regression method.

#### 12. Statistical analysis and figure generation

Unless otherwise stated, tests were two-sided and multiple-testing correction used the Benjamini-Hochberg procedure with an FDR threshold of 0.05. The direction-specific motif-enrichment tests were one-sided as specified above. Figure 4 comprised the 18-gene promoter motif grammar (panel A), selected-gene RNA-ATAC effects (panel B), genome-wide ATAC motif-family enrichment (panel C), selected-gene RNA-EZH2 effects (panel D) and genome-wide EZH2 interval motif-burden models (panel E). The figure was generated with Python v3.13.12, pandas v2.3.3, NumPy v2.4.4 and Matplotlib v3.10.8 as a 180 × 250-mm native-vector PDF and a 300-dpi PNG; the EZH2 motif-burden models used SciPy v1.17.1 and statsmodels v0.14.6.

### Atomic Force Microscopy

AFM measurements were conducted using JPK NanoWizard V BioScience system (Bruker) with an inverted optical microscope, and colloidal silicon oxide spherical tips (∼6.6 μm diameter) attached to cantilevers with ∼0.08 N m-1 nominal spring constant (NanoAndMore).

Cells were cultured in 35 × 10 mm Petri dishes and then transferred to the stage of AFM. The thermal tune method was used to determine the cantilever spring constant. Additionally, the photodiode sensitivity was calibrated against cell-free areas. Measurements were conducted at 37.5°C (± 0.1°C) using a temperature-controlled heating holder (Bruker).

Force measurements were performed in force-volume mode over 85 × 85 μm² scan areas. The experimental setup and an example of an AFM force curve is shown in Supplementary Figure 7. AFM measurements were performed on three independent biological replicates for each condition (WT, KO, and NLS). For each biological replicate, at least one independent field of view was analyzed in FV mode. Each FV dataset generated a grid of force-indentation curves (64 × 64 pixels for D0 with a pixel size of ∼1.33 μm and 32 × 32 pixels for D7 with a pixel size of ∼2.66 μm), which were subsequently used for stiffness analysis. Cantilever approach and retraction velocities were set to 4 μm s-1, and constant loading force of 27.198 nN was applied for the indentation of cell surfaces.

Force curves free from noise and artifacts were analyzed using an in-house MATLAB software. Young’s modulus of elasticity (E), representing ECM stiffness, was determined using the Hertz model:

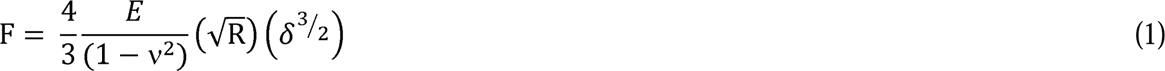

where F is the loading force, R is the AFM tip radius, δ is the surface indentation depth, and ν is the Poisson ratio (set to 0.5 assuming incompressibility). For each force curve, the Hertzian fit range was set to 40% of the maximum indentation to ensure elasticity. Data points with fitting goodness < 0.8 were excluded from analysis.

The JPK DirectOverlay module (Bruker) was used to integrate optical images into the AFM workspace, enabling the direct selection of predefined AFM scan areas and measurement pixels within the optical images. Representative height images and corresponding FV maps in D0 and D7 sets were generated using JPK DP software (Bruker). In the height images, plane fitting, line leveling, and median filtering were applied to remove background variations. In the FV maps, invalid pixels were replaced with the minimum *E* values, and the map data range was recorded as the colorbar scale range.

The *E* values were presented as mean ± standard error (SE) from Weibull probability density function fitting using OriginPro software (OriginLab). Statistical comparisons were made using one-way ANOVA for ECM stiffness differences among cell types at D0 and D7, and independent t-test to compare values between D0 and D7 using SPSS Statistics software (IBM). Significance was set at p < 0.001.

### Neuronal Differentiation of mESCs

Neuronal differentiation of mESCs was carried out essentially as described, with minor modifications^89^. At day 0, mESCs were aggregated into embryoid bodies (EBs) in mESC medium lacking LIF and retinoic acid (RA). On day 2, EBs were transferred to gelatin-coated 24-well plates, and the medium was changed to mESC medium without LIF supplemented with 500 nM RA. The medium was subsequently replaced every 2 days with fresh mESC medium without LIF containing 500 nM RA until day 7 or day 14, at which time downstream analyses were performed. RA concentration, EB seeding density, and the timing of EB plating were empirically optimized in preliminary experiments; unless otherwise indicated, all experiments were performed using 500 nM RA, the recommended EB seeding density, and EB plating on day 2.

### Tumor/Teratoma Induction Studies

All procedures involving animals were conducted in strict accordance with institutional and federal guidelines and were approved by the New York University Abu Dhabi Institutional Animal Care and Use Committee. Female BALB/c nude mice (athymic; 5–6 weeks of age; body weight 25–30 g) were used for teratoma formation studies. Upon arrival, mice were acclimatized for one week prior to the experiments. Animals were housed in a specific pathogen-free ventilated cage system with autoclaved woodchip bedding, and irradiated rodent chow and sterile water were provided *ad libitum*. Temperature was maintained at 21–24 °C with 40–60% relative humidity under a 12/12 h light/dark cycle.

A total of 18 female BALB/c nude mice were randomly assigned into three experimental groups (n = 6 per group). Teratomas were established via subcutaneous injection of mouse embryonic stem cell (mESCs) lines: wild-type (WT), knockout (KO, lacking β-actin), and nuclear localization signal-tagged β-actin construct into the KO background (hereafter referred to as NLS) cells. Prior to injection, cells were harvested during the logarithmic growth phase, counted using an automated cell counter, and assessed for viability (> 90%) using trypan blue exclusion. Each mouse received a single subcutaneous injection into the right flank consisting of 4 × 10^5^ cells suspended in 0.1 mL sterile 10 mM PBS (pH 7.4). Following inoculation, animals were weighed daily and monitored for general health status, mobility, grooming behavior, and signs of pain or distress. Tumor growth was assessed periodically using digital calipers. Tumor volume (V) was calculated using the standard formula^90^:

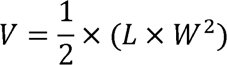

where L represents the longest tumor diameter and W the perpendicular diameter. Animals were monitored daily for signs of distress or weight loss. Humane endpoints were defined in accordance with IACUC guidelines as per protocol number 21-0005. After 18 days post-injection, mice were euthanized by CO inhalation followed by cervical dislocation. Tumors were excised promptly and processed for histological analysis.

### Teratomas histological analysis

Excised tissues were fixed in 10% neutral buffered formalin for 48 h at RT, processed through graded ethanol and xylene, embedded in paraffin, and sectioned at 4–5 µm thickness. Sections were stained with hematoxylin and eosin (H&E) for histopathological evaluation using light microscopy^91^.

### Teratoma cryopreservation, sectioning and immunofluorescence

Teratomas were fixed overnight in 4% paraformaldehyde (PFA), cryoprotected in 30% (w/v) sucrose in PBS at 4 °C until the tissue sank, embedded in optimal cutting temperature (OCT) medium, and frozen in precooled isopropanol on dry ice. Blocks were stored at -80 °C until sectioning. Cryostar NX70 was used to obtain 10 μm thick cryosections that were collected on Superfrost Plus slides, air-dried for 1 h at RT, dried further with cold air for 15 min, and stored at -80 °C until use. For immunofluorescence staining, sections were brought to room temperature, rehydrated in PBS (3 × 5 min), permeabilized in PBS–Triton X for 30 min, and blocked for 1 h at RT in a humidified chamber. Primary antibodies diluted (1:200) in blocking solution were applied overnight at 4 °C, followed by PBS rinse and washes in PBS–Tween 20 (3 × 10 min). Secondary antibodies (1:400) were then applied for 1 h at room temperature in the dark, followed by PBS rinse and PBS–Tween 20 washes (3 × 10 min). Slides were mounted with mounting medium, cured for 2–3 h RT, and stored at 4 °C protected from light until imaging. Finally, images were acquired on a fluorescence microscope (NIKON LV100 upright microscope, and the images were processed with the ECLIPSE LV software).^92^.

## Data Availability

All experiments are done from three biological replicates unless stated otherwise. The RNA-Seq data set of WT, KO and NLS mESCs used in this study (Figures 2) is publicly available in the Gene Expression Omnibus (GEO) database under accession number GSE327304. The RNA-Seq data of WT, KO and NLS mESCs at day 4, day 7 and day 14 (Figures 4, Figure 6, Supplementary figures 6-9) are also publicly available in the Gene Expression Omnibus (GEO) under the same accession number GSE327304. ATAC-Seq data of WT, KO and NLS mESCs is publicly available in the Gene Expression Omnibus (GEO) database under accession number GSE327117. EZH2 ChIP-Seq data of WT, KO and NLS mESCs is publicly available in the Gene Expression Omnibus (GEO) database under accession number GSE346145. Mass spectrometry data sets are available via ProteomeXchange with identifier PXD080676. Source data are provided with this paper.

## Code Availability

Codes are available at https://github.com/NYUAD-Core-Bioinformatics/beta_actin_stemcell_pluripotency.

## Acknowledgments

This work is supported by grants from NYU Abu Dhabi, the Sheikh Hamdan Bin Rashid Al Maktoum Award for Medical Sciences and Tamkeen under the NYU Abu Dhabi Research Institute Award to the NYU Abu Dhabi Center for Genomics and Systems Biology (ADHPG-CGSB) to PP. We thank the NYU Abu Dhabi Center for Genomics and Systems Biology, in particular Marc Arnoux and Mehar Sultana for RNA sequencing, as well as the NYU Abu Dhabi Core Technology Platform Resources, including the NYU Abu Dhabi imaging center, in particular Dr. Rachid Rezgui, for help with the microscopy. We appreciate the computational platform provided by the Center for Genomics and Systems Biology and the NYU Abu Dhabi HPC team and are especially thankful to Dr. Nizar Drou for technical help.

## Author Contributions Statement

PP conceived the research and wrote the manuscript. CC, AAN and ZA performed imaging, biochemical experiments, and data analysis and prepared the figures with the help of SCD. NHE performed all functional genomics analyses including RNA-Seq and ATAC-Seq; ES, GAS performed the bioinformatics analysis; LA and MSA performed mass spectrometry; MM and PL planned and performed the teratoma assay in nude mice and, together with CC, characterized the teratomas by immunohistochemistry; MD and MAQ performed all AFM measurements; PP, CB, SCD and ML were involved in the design and production of β-actin KO mESCs and NLS β-actin mESCs. PP supervised all the research. All authors read and approved the manuscript.

## Competing Interests Statement

The authors declare no competing interests.

## Notes

https://www.ncbi.nlm.nih.gov/geo/

### Competing Interest Statement

The authors have declared no competing interest.

### Summary of Updates

In the revised manuscript, we have strengthened the study by including new bioinformatics analysis integrating RNA-Seq and ATAC-Seq in all conditions (WT, KO and NLS mESCs) with motif grammar obtained from a gene regulatory network and new ChIP-Seq data obtained in the three genotypes with an antibody against EZH2, the catalytic subunit of the repressive PRC2 chromatin remodeling complex. We have also included new quantitative mass spectrometry data (in nuclear and cytoplasmic extracts from WT, KO and NLS mESCs) supporting that alterations of gene activity due to changes in chromatin accessibility upon β-actin depletion are supported at the protein level. Together these results identify a cell identity gene program regulated at chromatin level by nuclear β-actin. Pluripotency gene loci are downregulated, there is aberrant expression of ECM genes, changes in ECM stiffness and this activates mechanotransduction by facilitating nuclear uptake of the transcription factor YAP1.

## References

1 Bonev, B. & Cavalli, G. Organization and function of the 3D genome. Nature Reviews Genetics 17, 661–678 (2016).

2 Rowley, M. J. & Corces, V. G. Organizational principles of 3D genome architecture. Nature Reviews Genetics 19, 789–800 (2018).

3 Schlesinger, S. & Meshorer, E. Open chromatin, epigenetic plasticity, and nuclear organization in pluripotency. Developmental Cell 48, 135–150 (2019).

4 Pękowska, A. et al. Gain of CTCF-anchored chromatin loops marks the exit from naive pluripotency. Cell Systems 7, 482–495. e410 (2018).

5 Denholtz, M. & Plath, K. Pluripotency in 3D: genome organization in pluripotent cells. Current opinion in cell biology 24, 793–801 (2012).

6 Apostolou, E. et al. Genome-wide chromatin interactions of the Nanog locus in pluripotency, differentiation, and reprogramming. Cell stem cell 12, 699–712 (2013).

7 King, H. W. & Klose, R. J. The pioneer factor OCT4 requires the chromatin remodeller BRG1 to support gene regulatory element function in mouse embryonic stem cells. Elife 6, e22631 (2017).

8 Chandramohan, D. et al. Dual role of Oct4 and Sox2 in controlling the developmental capacity and timing of tissue morphogenesis in the embryonic lineage. Developmental Cell (2025).

9 Kim, Y. S. et al. Rap1 controls epiblast morphogenesis in sync with the pluripotency states transition. Developmental cell 57, 1937–1956. e1938 (2022).

10 Dixon, J. R. et al. Chromatin architecture reorganization during stem cell differentiation. Nature 518, 331–336 (2015).

11 Denholtz, M. et al. Long-range chromatin contacts in embryonic stem cells reveal a role for pluripotency factors and polycomb proteins in genome organization. Cell stem cell 13, 602–616 (2013).

12 Hnisz, D. et al. Super-enhancers in the control of cell identity and disease. Cell 155, 934–947 (2013).

13 Schmitt, A. D. et al. A compendium of chromatin contact maps reveals spatially active regions in the human genome. Cell reports 17, 2042–2059 (2016).

14 Bhat, P., Honson, D. & Guttman, M. Nuclear compartmentalization as a mechanism of quantitative control of gene expression. Nature Reviews Molecular Cell Biology 22, 653–670 (2021).

15 Fueyo, R. et al. Lineage specific transcription factors and epigenetic regulators mediate TGFβ-dependent enhancer activation. Nucleic acids research 46, 3351–3365 (2018).

16 Isbel, L., Grand, R. S. & Schübeler, D. Generating specificity in genome regulation through transcription factor sensitivity to chromatin. Nature Reviews Genetics 23, 728–740 (2022).

17 Nagy, G., Bojcsuk, D., Tzerpos, P., Cseh, T. & Nagy, L. Lineage-determining transcription factor-driven promoters regulate cell type-specific macrophage gene expression. Nucleic acids research 52, 4234–4256 (2024).

18 Gorkin, D. U., Leung, D. & Ren, B. The 3D genome in transcriptional regulation and pluripotency. Cell stem cell 14, 762–775 (2014).

19 Zheng, H. & Xie, W. The role of 3D genome organization in development and cell differentiation. Nature reviews Molecular cell biology 20, 535–550 (2019).

20 Kagey, M. H. et al. Mediator and cohesin connect gene expression and chromatin architecture. Nature 467, 430–435 (2010).

21 Ulferts, S., Lopes, M., Miyamoto, K. & Grosse, R. Nuclear actin dynamics and functions at a glance. Journal of cell science 137, jcs261630 (2024).

22 Xie, X. et al. β Actin dependent global chromatin organization and gene expression programs control cellular identity. The FASEB Journal 32, 1296–1314 (2018).

23 Mahmood, S. R. et al. β-actin dependent chromatin remodeling mediates compartment level changes in 3D genome architecture. Nature communications 12, 5240 (2021).

24 Mahmood, S. R., Said, N. H. E., Gunsalus, K. C. & Percipalle, P. β-actin mediated H3K27ac changes demonstrate the link between compartment switching and enhancer-dependent transcriptional regulation. Genome Biology 24, 18 (2023).

25 Visa, N. & Percipalle, P. Nuclear functions of actin. Cold Spring Harbor perspectives in biology 2, a000620 (2010).

26 Percipalle, P. et al. Actin bound to the heterogeneous nuclear ribonucleoprotein hrp36 is associated with Balbiani ring mRNA from the gene to polysomes. The Journal of cell biology 153, 229–236 (2001).

27 Percipalle, P. et al. An actin–ribonucleoprotein interaction is involved in transcription by RNA polymerase II. Proceedings of the National Academy of Sciences 100, 6475–6480 (2003).

28 Percipalle, P. et al. Nuclear actin is associated with a specific subset of hnRNP A/B-type proteins. Nucleic acids research 30, 1725–1734 (2002).

29 Hosny El Said, N., et al. Nuclear actin-dependent Meg3 expression suppresses metabolic genes by affecting the chromatin architecture at sites of elevated H3K27 acetylation levels. Nucleic acids research 53, gkaf280 (2025).

30 Xie, X., Jankauskas, R., Mazari, A. M., Drou, N. & Percipalle, P. β-actin regulates a heterochromatin landscape essential for optimal induction of neuronal programs during direct reprograming. PLoS Genetics 14, e1007846 (2018).

31 Gjorgjieva, T. et al. Loss of β actin leads to accelerated mineralization and dysregulation of osteoblast differentiation genes during osteogenic reprogramming. Advanced Science 7, 2002261 (2020).

32 Al-Sayegh, M. et al. β-actin contributes to open chromatin for activation of the adipogenic pioneer factor CEBPA during transcriptional reprograming. Molecular biology of the cell 31, 2511–2521 (2020).

33 Kitisin, K. et al. TGF-β signaling in development. Science’s STKE 2007, cm1–cm1 (2007).

34 Watt, F. M. & Huck, W. T. Role of the extracellular matrix in regulating stem cell fate. Nature reviews Molecular cell biology 14, 467–473 (2013).

35 Gaarenstroom, T. & Hill, C. S. TGF-β signaling to chromatin: how Smads regulate transcription during self-renewal and differentiation. Semin Cell Dev Biol 32, 107–118 (2014). 10.1016/j.semcdb.2014.01.009

36 Fontana, L. et al. Fibronectin is required for integrin o: vβ6 mediated activation of latent TGF β complexes containing LTBP 1. The FASEB journal 19, 1798–1808 (2005).

37 Buscemi, L. et al. The single-molecule mechanics of the latent TGF-β1 complex. Current biology 21, 2046–2054 (2011).

38 Murphy-Ullrich, J. E. & Poczatek, M. Activation of latent TGF-beta by thrombospondin-1: mechanisms and physiology. Cytokine Growth Factor Rev 11, 59–69 (2000). 10.1016/s1359-6101(99)00029-5

39 Xie, X. & Percipalle, P. Elevated transforming growth factor β signaling activation in β actin knockout mouse embryonic fibroblasts enhances myofibroblast features. Journal of Cellular Physiology 233, 8884–8895 (2018).

40 Miyamoto, K. et al. Nuclear Wave1 is required for reprogramming transcription in oocytes and for normal development. Science 341, 1002–1005 (2013).

41 Padhi, A. & Nain, A. S. ECM in differentiation: a review of matrix structure, composition and mechanical properties. Annals of biomedical engineering 48, 1071–1089 (2020).

42 May, M., Denecke, B., Schroeder, T., Götz, M. & Faissner, A. Cell tracking in vitro reveals that the extracellular matrix glycoprotein Tenascin-C modulates cell cycle length and differentiation in neural stem/progenitor cells of the developing mouse spinal cord. Biology open 7, bio027730 (2018).

43 Zhang, Z. et al. Fibroblast derived tenascin C promotes S chwann cell migration through β1 integrin dependent pathway during peripheral nerve regeneration. Glia 64, 374–385 (2016).

44 Schaberg, E., Götz, M. & Faissner, A. The extracellular matrix molecule tenascin-C modulates cell cycle progression and motility of adult neural stem/progenitor cells from the subependymal zone. Cellular and Molecular Life Sciences 79, 244 (2022).

45 Quondamatteo, F. et al. Fibrillin-1 and fibrillin-2 in human embryonic and early fetal development. Matrix biology 21, 637–646 (2002).

46 Humphries, M. J., Obara, M., Olden, K. & Yamada, K. M. Role of fibronectin in adhesion, migration, and metastasis. Cancer investigation 7, 373–393 (1989).

47 Garcıa, A. J., Vega, M. a. D. & Boettiger, D. Modulation of cell proliferation and differentiation through substrate-dependent changes in fibronectin conformation. Molecular biology of the cell 10, 785–798 (1999).

48 Liu, C. et al. Id1 expression promotes T regulatory cell differentiation by facilitating TCR costimulation. The Journal of Immunology 193, 663–672 (2014).

49 Zhao, Q. et al. Combined Id1 and Id3 deletion leads to severe erythropoietic disturbances. PloS one 11, e0154480 (2016).

50 Jung, S. et al. Id proteins facilitate self-renewal and proliferation of neural stem cells. Stem cells and development 19, 831–841 (2010).

51 Ying, Q.-L., Nichols, J., Chambers, I. & Smith, A. BMP induction of Id proteins suppresses differentiation and sustains embryonic stem cell self-renewal in collaboration with STAT3. Cell 115, 281–292 (2003).

52 Dong, L., Lyu, X., Faleti, O. D. & He, M. L. The special stemness functions of Tbx3 in stem cells and cancer development. Semin Cancer Biol 57, 105–110 (2019). 10.1016/j.semcancer.2018.09.010

53 Weidgang, C. E. et al. TBX3 directs cell-fate decision toward mesendoderm. Stem cell reports 1, 248–265 (2013).

54 Hofmann, M. et al. WNT signaling, in synergy with T/TBX6, controls Notch signaling by regulating Dll1 expression in the presomitic mesoderm of mouse embryos. Genes & development 18, 2712–2717 (2004).

55 Zhao, S. et al. Dppa3 facilitates self-renewal of embryonic stem cells by stabilization of pluripotent factors. Stem Cell Research & Therapy 13, 169 (2022).

56 Sund, M., Maeshima, Y. & Kalluri, R. Bifunctional promoter of type IV collagen COL4A5 and COL4A6 genes regulates the expression of α5 and α6 chains in a distinct cell-specific fashion. Biochemical Journal 387, 755–761 (2005).

57 Graf, U., Casanova, E. A. & Cinelli, P. The role of the leukemia inhibitory factor (LIF)—pathway in derivation and maintenance of murine pluripotent stem cells. Genes 2, 280–297 (2011).

58 Tai, C.-I. & Ying, Q.-L. Gbx2, a LIF/Stat3 target, promotes reprogramming to and retention of the pluripotent ground state. Journal of cell science 126, 1093–1098 (2013).

59 Parisi, S. et al. Klf5 is involved in self-renewal of mouse embryonic stem cells. Journal of cell science 121, 2629–2634 (2008).

60 Sun, Z., Guo, S. S. & Fässler, R. Integrin-mediated mechanotransduction. Journal of Cell Biology 215, 445–456 (2016).

61 Chiquet-Ehrismann, R. & Tucker, R. P. Tenascins and the importance of adhesion modulation. Cold Spring Harbor Perspectives in Biology 3, a004960 (2011).

62 Pankov, R. & Yamada, K. M. Fibronectin at a glance. Journal of cell science 115, 3861–3863 (2002).

63 Kühn, K. Basement membrane (type IV) collagen. Matrix Biology 14, 439–445 (1995).

64 Taipale, J. & Keski Oja, J. Growth factors in the extracellular matrix. The FASEB Journal 11, 51–59 (1997).

65 Saraswathibhatla, A., Indana, D. & Chaudhuri, O. Cell–extracellular matrix mechanotransduction in 3D. Nature reviews Molecular cell biology 24, 495–516 (2023).

66 Xie, X., Deliorman, M., Qasaimeh, M. A. & Percipalle, P. The relative composition of actin isoforms regulates cell surface biophysical features and cellular behaviors. Biochimica et Biophysica Acta (BBA)-General Subjects 1862, 1079–1090 (2018).

67 Deliorman, M., Gjorgjieva, T., Xie, X., Percipalle, P. & Qasaimeh, M. A. in 2023 International Conference on Manipulation, Automation and Robotics at Small Scales (MARSS). 1–6.

68 Viji Babu, P. K., Rianna, C., Mirastschijski, U. & Radmacher, M. Nano-mechanical mapping of interdependent cell and ECM mechanics by AFM force spectroscopy. Scientific reports 9, 12317 (2019).

69 Technau, U. & Scholz, C. B. Origin and evolution of endoderm and mesoderm. International Journal of Developmental Biology 47, 531–539 (2003).

70 Technau, U. Brachyury, the blastopore and the evolution of the mesoderm. Bioessays 23, 788–794 (2001).

71 De Val, S. & Black, B. L. Transcriptional control of endothelial cell development. Developmental cell 16, 180–195 (2009).

72 Pikkarainen, S., Tokola, H., Kerkelä, R. & Ruskoaho, H. GATA transcription factors in the developing and adult heart. Cardiovasc Res 63, 196–207 (2004). 10.1016/j.cardiores.2004.03.025

73 Rogers, C. D., Moody, S. A. & Casey, E. S. Neural induction and factors that stabilize a neural fate. Birth Defects Research Part C: Embryo Today: Reviews 87, 249–262 (2009).

74 Aruga, J. & Mikoshiba, K. Role of BMP, FGF, calcium signaling, and Zic proteins in vertebrate neuroectodermal differentiation. Neurochemical research 36, 1286–1292 (2011).

75 Zhang, X. et al. Pax6 is a human neuroectoderm cell fate determinant. Cell Stem Cell 7, 90–100 (2010). 10.1016/j.stem.2010.04.017

76 Stachowiak, M. K. & Stachowiak, E. K. Evidence based theory for integrated genome regulation of ontogeny—An unprecedented role of nuclear FGFR1 signaling. Journal of cellular physiology 231, 1199–1218 (2016).

77 Konstantinides, N. & Desplan, C. Neuronal differentiation strategies: insights from single-cell sequencing and machine learning. Development 147, dev193631 (2020).

78 Ayanlaja, A. A. et al. Distinct features of doublecortin as a marker of neuronal migration and its implications in cancer cell mobility. Frontiers in molecular neuroscience 10, 199 (2017).

79 Gaetani, R. et al. When stiffness matters: mechanosensing in heart development and disease. Frontiers in Cell and Developmental Biology 8, 334 (2020).

80 Majkut, S. et al. Heart-specific stiffening in early embryos parallels matrix and myosin expression to optimize beating. Current biology 23, 2434–2439 (2013).

81 Dupont, S. et al. Role of YAP/TAZ in mechanotransduction. Nature 474, 179–183 (2011).

82 Lv, H. et al. Mechanism of regulation of stem cell differentiation by matrix stiffness. Stem cell research & therapy 6, 103 (2015).

83 Bulic-Jakus, F., Katusic Bojanac, A., Juric-Lekic, G., Vlahovic, M. & Sincic, N. Teratoma: from spontaneous tumors to the pluripotency/malignancy assay. Wiley Interdiscip Rev Dev Biol 5, 186–209 (2016). 10.1002/wdev.219

84 Hillmer, R. E. & Link, B. A. The roles of Hippo signaling transducers Yap and Taz in chromatin remodeling. Cells 8, 502 (2019).

85 Wang, H. et al. Regulation of YAP activity by nuclear G-actin binding. Nucleic acids research 54, gkag248 (2026).

86 Piersma, B., Bank, R. A. & Boersema, M. Signaling in fibrosis: TGF-β, WNT, and YAP/TAZ converge. Frontiers in medicine 2, 59 (2015).

87 Song, I. & Dityatev, A. Crosstalk between glia, extracellular matrix and neurons. Brain research bulletin 136, 101–108 (2018).

88 Concordet, J.-P. & Haeussler, M. CRISPOR: intuitive guide selection for CRISPR/Cas9 genome editing experiments and screens. Nucleic acids research 46, W242–W245 (2018).

89 Witteveldt, J. & Macias, S. Differentiation of mouse embryonic stem cells to neuronal cells using hanging droplets and retinoic acid. Bio-protocol 9 (2019).

90 Tomayko, M. M. & Reynolds, C. P. Determination of subcutaneous tumor size in athymic (nude) mice. Cancer chemotherapy and pharmacology 24, 148–154 (1989).

91. Suvarna, K. S., Layton, C. & Bancroft, J. D. Bancroft’s theory and practice of histological techniques E-Book. (Elsevier health sciences, 2018).

92 Nelakanti, R. V., Kooreman, N. G. & Wu, J. C. Teratoma formation: a tool for monitoring pluripotency in stem cell research. Curr Protoc Stem Cell Biol 32, 4a.8.1–4a.8.17 (2015). 10.1002/9780470151808.sc04a08s32

