## Supplementary figure legends and figures for "Nuclear β-actin–dependent chromatin accessibility governs stem cell pluripotency and extracellular matrix gene programs to maintain cellular biomechanics for cell lineage decisions"

**Supplementary figures and tables legends**

**Supplementary figure 1.** Pairwise differential expression analysis among the three genotypes, WT, KO and NLS mESCs visualized by MA and volcano plots. (A–C) MA plots displaying the relationship between log fold change (y-axis) versus the mean of normalized counts (x-axis), for (A) KO vs WT, (B) KO vs NLS and (C) WT vs NLS. Each dot represents a single gene, only red marked genes are significantly differentially expressed, while non-significant genes are shown in grey. Horizontal reference lines indicate thresholds for fold change. (D, E) Volcano plots showing the distribution of genes based on $\log_2$ fold change (x-axis) and $-\log_{10}$ adjusted p-value (y-axis) for (D) KO vs NLS and (E) WT vs NLS. Significantly upregulated and downregulated genes are highlighted in Red, with vertical and horizontal dashed lines indicating fold change and significance cutoffs, respectively. Together, these plots summarize the magnitude and statistical significance of differential gene expression across all genotypes.

**Supplementary figure 2.** Gene Ontology (GO) enrichment analysis based on all differentially expressed genes identified by RNAseq analysis between experimental conditions in WT, KO and NLS mESCs. Bar plots show the top enriched GO terms identified from differentially expressed genes comparing KO vs NLS and WT vs NLS. The analyses show Cellular Component and Molecular Function associated gene ontology terms plotted in descending order based on their significance. Each bar represents an individual GO term. Bar length represents the number of genes associated with each enriched GO term (Count), and bar color indicates the adjusted *P* value (p adjust), with darker red corresponding to higher statistical significance. Together, these panels highlight functional differences and shared biological processes across WT, KO, and NLS conditions.

**Supplementary figure 3.** Gene Ontology (GO) enrichment analysis of ATAC-seq differential chromatin accessibility in KO versus WT conditions. (A–C) Gene Ontology enrichment analysis of regions with differential chromatin accessibility identified by ATAC-seq comparing knockout (KO) and wild type (WT) conditions. (A) Enriched GO terms associated with biological processes. (B) Enriched GO terms associated with cellular components. (C) Enriched GO terms associated with molecular functions. Each point (or bar) represents a significantly enriched GO term derived from genes linked to differentially accessible regions. Enrichment significance is indicated by adjusted p-values and/or $-\log_{10}$(p-value), while the number of associated genes is reflected by point size or bar length where applicable. Color gradients indicate the magnitude of enrichment.

These analyses reveal functional categories and regulatory features associated with chromatin accessibility changes between KO and WT conditions.

**Supplementary figure 4.** Integrated genome visualization (IGVs) shows that depletion of β-actin from the mESCs nuclei leads to increased heterochromatin levels at regulatory regions of pluripotency genes which are directly downregulated at the transcriptional level. RT-qPCR showing the *Oct4*, *Sox2*, *Klf4* and *Nanog* relative abundance vs *Gapdh* in WT, KO and in NLS. Data are shown as mean expression fold change ± SD ($n = 4$). P-values indicated are based on one-way ANOVA multiple comparisons in GraphPad Prism version 10. Green boxes identify the upstream gene regulatory regions.

**Supplementary figure 5**. (A) Genome-wide integration of ATAC-seq and RNA-seq for NLS versus WT mESCs. Each point is one gene that is significantly differential in both layers (RNA: adjusted $p < 0.05$ and $|\log_2FC| \geq 0.25$; ATAC: $FDR < 0.05$ and $|\log_2FC| \geq 0.1$; $n = 2{,}013$ genes).

The x-axis is the ATAC promoter/nearest-TSS $\log_2$ fold-change and the y-axis is the RNA $\log_2$ fold-change. Quadrants define the integrated regulatory state: Activated (open + up, n = 183) and Repressed (closed + down, n = 265) are concordant (448 genes total); the off-diagonal quadrants (Post-transcriptional, n = 508; Primed/open-not-expressed, n = 366) are discordant. Labelled genes are the curated ECM, stemness, lineage and mechanotransduction panel members detected in both layers, together with additional mechanosensor genes and the strongest genome-wide hits. (B-C) Directional accessibility and expression changes for panel genes in KO and NLS relative to WT. Side-by-side $\log_2$ fold-change heatmaps for curated panel genes that are significant-in-both in at least one contrast (n = 32 genes). Columns are the two integration contrasts, KO vs WT and NLS vs WT (the KO-vs-NLS integration is empty and is not shown). (Left) RNA $\log_2$FC ; (Right) ATAC $\log_2$FC; red = increased, blue = decreased. Asterisks denote significance in that layer/contrast (*** $p < 0.001$, ** $p < 0.01$, * $p < 0.05$). Row-side colors mark gene-set membership (ECM, Stemness, Lineage, Mechanosensing). These heatmaps show the direction and magnitude of change relative to WT for each contrast. (D) Chromatin accessibility and transcriptional changes are globally concordant in β-actin KO mESCs. Genome-wide integration of ATAC-seq and RNA-seq for KO versus WT mESCs. Each point is one gene that is significantly differential in both layers (RNA: adjusted $p < 0.05$ and $|\log_2\text{FC}| \geq 0.25$; ATAC: FDR < 0.05 and $|\log_2\text{FC}| \geq 0.1$; n = 1,624 genes). The x-axis is the ATAC promoter/nearest-TSS $\log_2$ fold-change and the y-axis is the RNA $\log_2$ fold-change. Quadrants define the integrated regulatory state: Activated (open + up, n = 280) and Repressed (closed + down, n = 431) are concordant (711 genes total); the off-diagonal quadrants (post-transcriptional, n = 211; Primed/open-not-expressed, n = 208) are discordant. Labelled genes are the curated ECM, stemness, lineage and mechanotransduction panel members detected in both layers, together with additional mechanosensory genes and the strongest genome-wide hits. The positive correlation between the two axes indicates that the majority of significant accessibility changes are matched by concordant expression changes.

**Supplementary figure 6.** Mass spectrometry analysis. (A-D) MS PCA plots (E) Bubble plot showing the top 20 significantly enriched biological pathways in the nuclear proteome of WT versus NLS cells. Bubble size represents the number of proteins associated with each pathway, while color indicates enrichment significance (−log10 adjusted q-value), with darker red indicating greater significance. The x-axis shows the rescaled −log10 adjusted q-values, normalized to a 0-5 range for visualization. (**F**) Bubble plot showing the top 20 significantly enriched biological pathways in the nuclear proteome of WT versus NLS cells. Bubble size represents the number of proteins associated with each pathway, while color indicates enrichment significance (−log10 adjusted q-value), with darker red indicating greater significance. The x-axis shows the rescaled −log10 adjusted q-values, normalized to a 0-5 range for visualization.

**Supplementary figure 7.** ECM composition and ECM stiffness measurements. (A-D) RNA levels for ECM factors obtained from the RNAseq data set in undifferentiated WT, KO and NLS mESCs. (**E**) A micrograph (scale bar: 40 µm) shows the AFM setup used for ECM stiffness measurements. ~6.6 µm-diameter spherical AFM tip was precisely aligned above cells of interest to enable force measurements in FV mode within 85 × 85 $\mu m^2$ scan area. In FV mode, tip approach-retraction process was repeated with a resolution of at least 32 × 32 pixels or 64 × 64 pixels. During measurements, cells were maintained at 37.5°C to ensure physiological conditions. (**F**) A typical force curve acquired on the cell surface is shown, where the AFM tip approaches (red arrow) and retracts (blue arrow) from the cell surface. The tip–cell contact point (black arrow) is identified, and the resulting surface indentation is defined by the constant applied loading force. For stiffness measurements, a Hertzian fit was used in the elastic region of the indentation (~40% of the total indentation, dashed purple line) to estimate the local Young's modulus of elasticity.

**Supplementary figure 8.** Differential expression analysis among the three genotypes, WT, KO and NLS mESCs at day 14 (D14) into neuronal differentiation. (A) Principal component analysis (PCA) analysis shows that WT (blue), KO (red) and NLS (green) mESCs samples exhibit distinctive clustered transcriptional profiles. (B) Hierarchical clustering heatmap demonstrates that WT, KO and NLS mESCs samples exhibit distinct transcriptomes. (C-E) MA plots displaying the relationship between mean of normalized counts ($\log_2$ average counts) and $\log_2$ fold change for (C) KO vs WT, (D) NLS vs WT, and (E) KO vs NLS. Genes with statistically significant differential expression are highlighted (red dots), while non-significant genes are shown in gray. Horizontal reference lines indicate thresholds for fold change where applicable. (F-H) Volcano plots showing differential gene expression based on $\log_2$ fold change (x-axis) and $-\log_{10}$ adjusted p-value (y-axis) for (F) KO vs WT and (G) NLS vs WT and (H) KO vs NLS. Significantly upregulated and downregulated genes are highlighted in Red, with vertical and horizontal dashed lines indicating fold change and significance cutoffs, respectively. Together, these plots summarize the magnitude and statistical significance of differential gene expression across all genotypes.

**Supplementary figure 9.** UpSet plots for RNAseq in WT and KO mESCs obtained at day 14 (D14). UpSet plots show intersections in a matrix, with the rows of the matrix corresponding to the sets, and the columns to the intersections between these sets. Genomic Regions represent different regions of the genome enriched in the different conditions. Each region corresponds to a specific genomic locus or set of loci. The connected dots represent the intersection of genomic

regions in WT and KO conditions. The bar shows the count of occurrences for each combination of sets. It provides a visual representation of the distribution of genomic regions and in WT and KO mESCs. BP, biological processes; MF, molecular functions; CC, cellular components.

**Supplementary figure 10.** Differential expression analysis among the three genotypes, WT, KO and NLS mESCs at day 4 (D4) into neuronal differentiation. (A) PCA analysis shows that WT (blue), KO (red) and NLS (green) mESCs samples show distinctive clustered transcriptional profiles. (B) Hierarchical clustering analyses demonstrate that WT, KO, and NLS mESCs samples exhibit distinct transcriptomes. (C-E) MA plots displaying the relationship between mean of normalized counts ($\log_2$ average counts) and $\log_2$ fold change for (C) KO vs WT, (D) NLS vs WT, and (E) KO vs NLS. Genes with statistically significant differential expression are highlighted (red dots), while non-significant genes are shown in gray. Horizontal reference lines indicate thresholds for fold change where applicable. (F-H) Volcano plots showing differential gene expression based on $\log_2$ fold change (x-axis) and $-\log_{10}$ adjusted p-value (y-axis) for (F) KO vs WT, (G) NLS vs WT, and (H) KO vs NLS. Significantly upregulated and downregulated genes are highlighted in distinct colors, with vertical and horizontal dashed lines indicating fold change and significance cutoffs, respectively. Together, these plots summarize the magnitude and statistical significance of differential gene expression across all genotypes.

**Supplementary figure 11.** UpSet plots for RNAseq in WT and KO mESCs obtained at day 4 (D4). UpSet plots show intersections in a matrix, with the rows of the matrix corresponding to the sets, and the columns to the intersections between these sets. Genomic Regions represent different regions of the genome enriched in different conditions. Each region corresponds to a specific genomic locus or set of loci. Intersection as connected dots represent the intersection of genomic

regions in WT and KO conditions. The bar shows the count of occurrences for each combination of sets. It provides a visual representation of the distribution of genomic regions and in WT and KO mESCs. BP, biological processes; MF, molecular functions; CC, cellular components.

**Supplementary figure 12.** Differential expression analysis among the three genotypes, WT, KO and NLS mESCs at day 7 (D7) into neuronal differentiation. (A) PCA analysis shows that WT (blue), KO (red) and NLS (green) mESCs samples show distinctive clustered transcriptional profiles. (B) Hierarchical clustering analyses demonstrate that WT, KO, and NLS mESCs samples exhibit distinct transcriptomes. (C-E) MA plots displaying the relationship between mean of normalized counts ($\log_2$ average counts) and $\log_2$ fold change for (C) KO vs WT, (D) NLS vs WT, and (E) KO vs NLS. Genes with statistically significant differential expression are highlighted (red dots), while non-significant genes are shown in gray. Horizontal reference lines indicate thresholds for fold change where applicable. (F-H) Volcano plots showing differential gene expression based on $\log_2$ fold change (x-axis) and $-\log_{10}$ adjusted p-value (y-axis) for (F) KO vs WT, (G) NLS vs WT, and (H) KO vs NLS. Significantly upregulated and downregulated genes are highlighted in distinct colors, with vertical and horizontal dashed lines indicating fold change and significance cutoffs, respectively. Together, these plots summarize the magnitude and statistical significance of differential gene expression across all genotypes.

**Supplementary figure 13.** UpSet plots for RNAseq in WT and KO mESCs obtained at day 7 (D7). UpSet plots show intersections in a matrix, with the rows of the matrix corresponding to the sets, and the columns to the intersections between these sets. Genomic Regions represent different regions of the genome enriched in different conditions. Each region corresponds to a specific genomic locus or set of loci. Intersection as connected dots represent the intersection of genomic

regions in WT and KO conditions. The bar shows the count of occurrences for each combination of sets. It provides a visual representation of the distribution of genomic regions and in WT and KO mESCs. BP, biological processes; MF, molecular functions; CC, cellular components.

**Supplementary video 1.** Cell fate lineage is compromised upon nuclear β-actin depletion. Live cell imaging shows that upon neuronal differentiation β-actin KO mESCs acquire a contracting phenotype compatible with altered expression of germ layers.

**Supplementary table 1.** Transcriptional profiling of WT, β-actin KO and NLS mESC by RNAseq at day 0 (D0).

**Supplementary table 2.** Analysis of chromatin accessibility by ATACseq performed on WT, βactin KO and NLS mESC by RNAseq.

**Supplementary table 3.** ATAC-seq and RNA-seq integration

**Supplementary table 4.** Normalized abundances of **4,068 nuclear proteins** identified and quantified across WT, KO, and NLS cells. Variable Importance in Projection (VIP) scores were calculated from the normalized protein abundances using multivariate analysis for the WT vs. KO and WT vs. NLS comparisons. Proteins with **VIP > 1** were considered significant and were used for subsequent pathway enrichment analysis

**Supplementary table 5**. Normalized abundances of **4,693 cytoplasmic proteins** identified and quantified across WT, KO, and NLS cells. Variable Importance in Projection (VIP) scores were calculated from the normalized protein abundances using multivariate analysis for the WT vs. KO and WT vs. NLS comparisons. Proteins with **VIP > 1** were considered significant and were used for subsequent pathway enrichment analysis

**Supplementary table 6.** Transcriptional profiling of WT, KO and NLS mESC by RNAseq at day 14 (D14) into the neuronal differentiation protocol

**Supplementary table 7.** Transcriptional profiling of WT, KO and NLS mESC by RNAseq at day 4 (D4) into the neuronal differentiation protocol

**Supplementary table 8.** Transcriptional profiling of WT, KO and NLS mESC by RNAseq at day 7 (D7) into the neuronal differentiation protocol

**Supplementary table 9.** Primers sequences used in qRT-PCR analyses

Supplementary figure 1

### RNA-Seq at Day 0

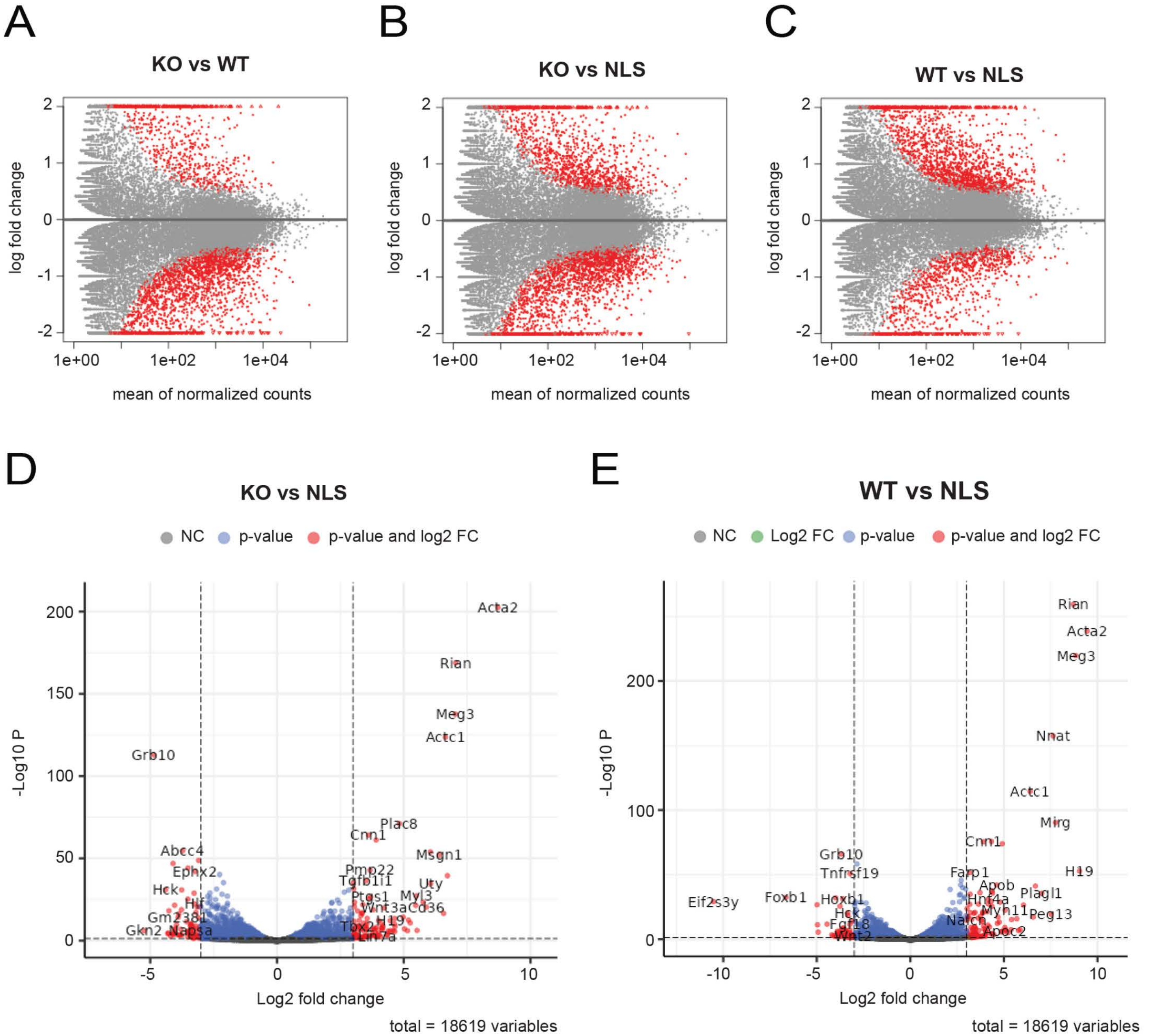

Supplementary figure 2

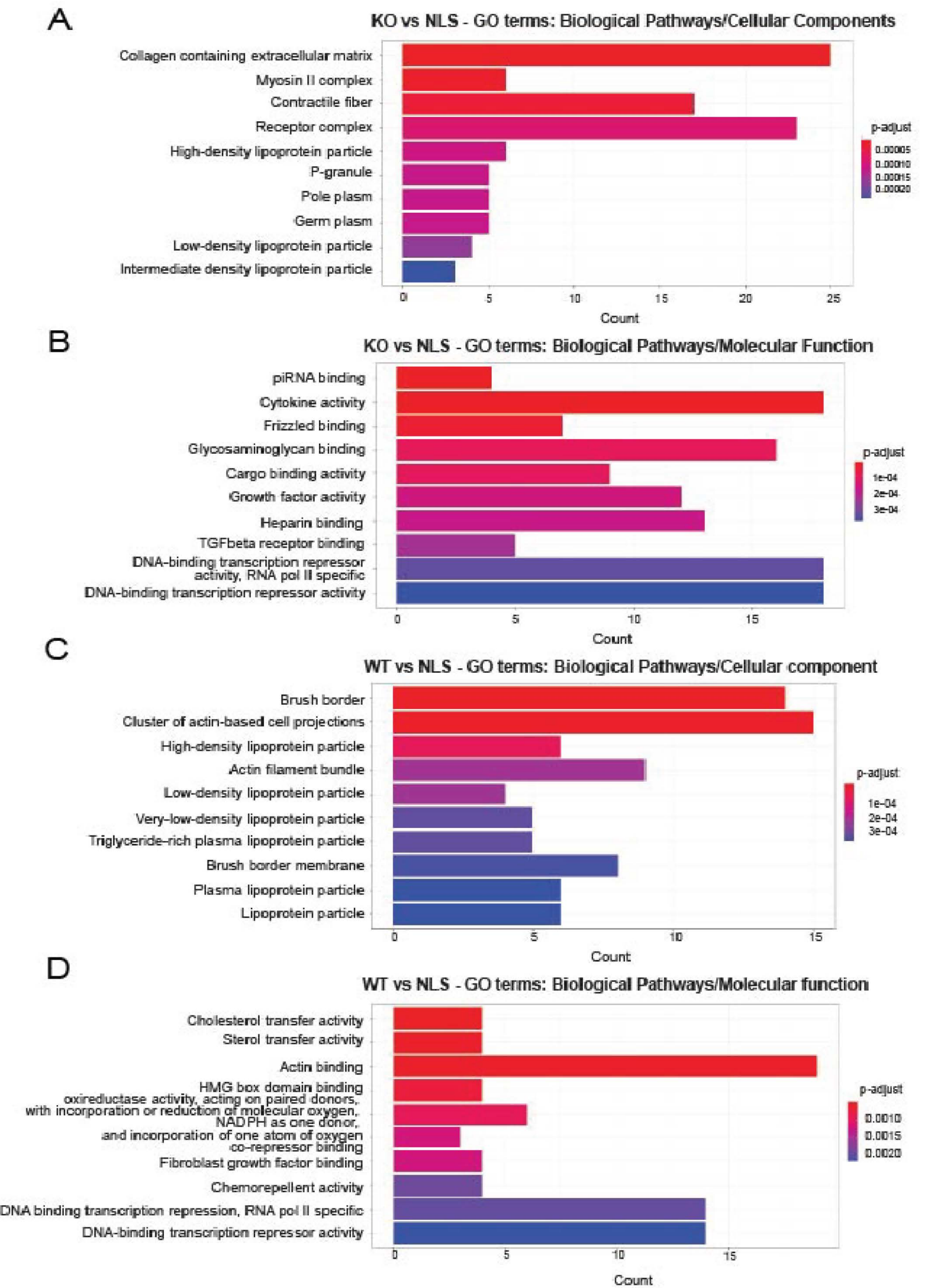

Supplementary figure 3

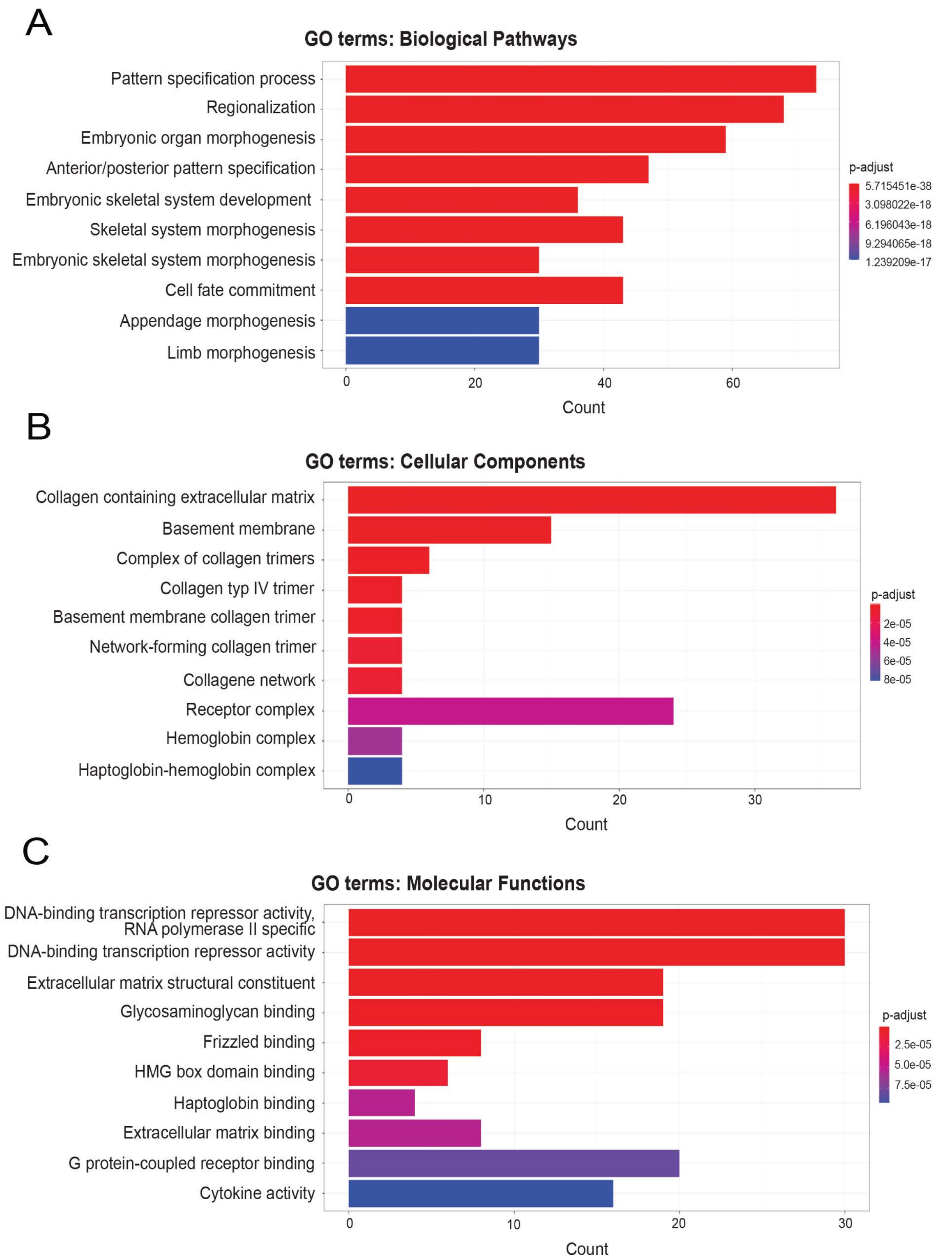

Supplementary figure 4

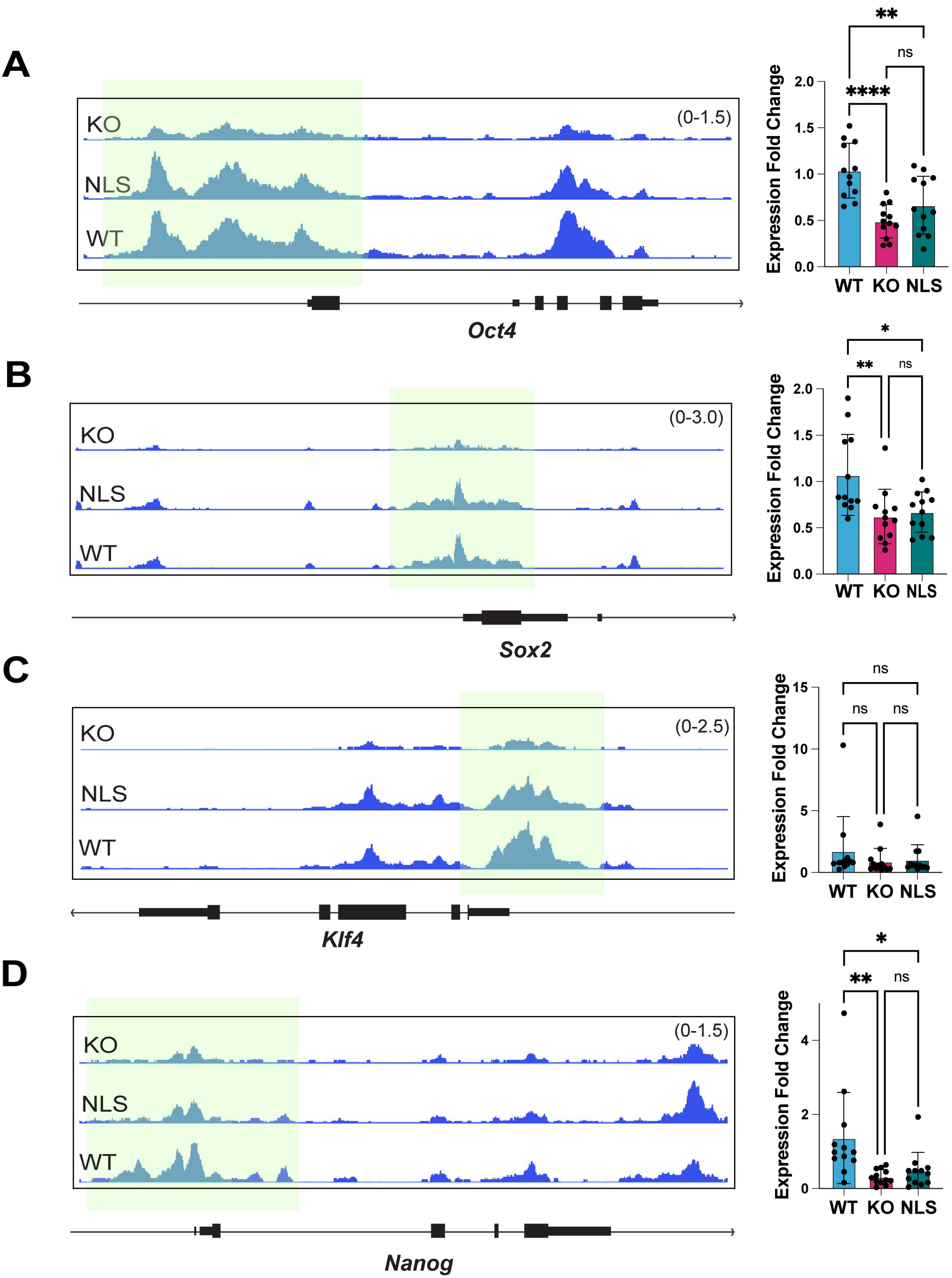

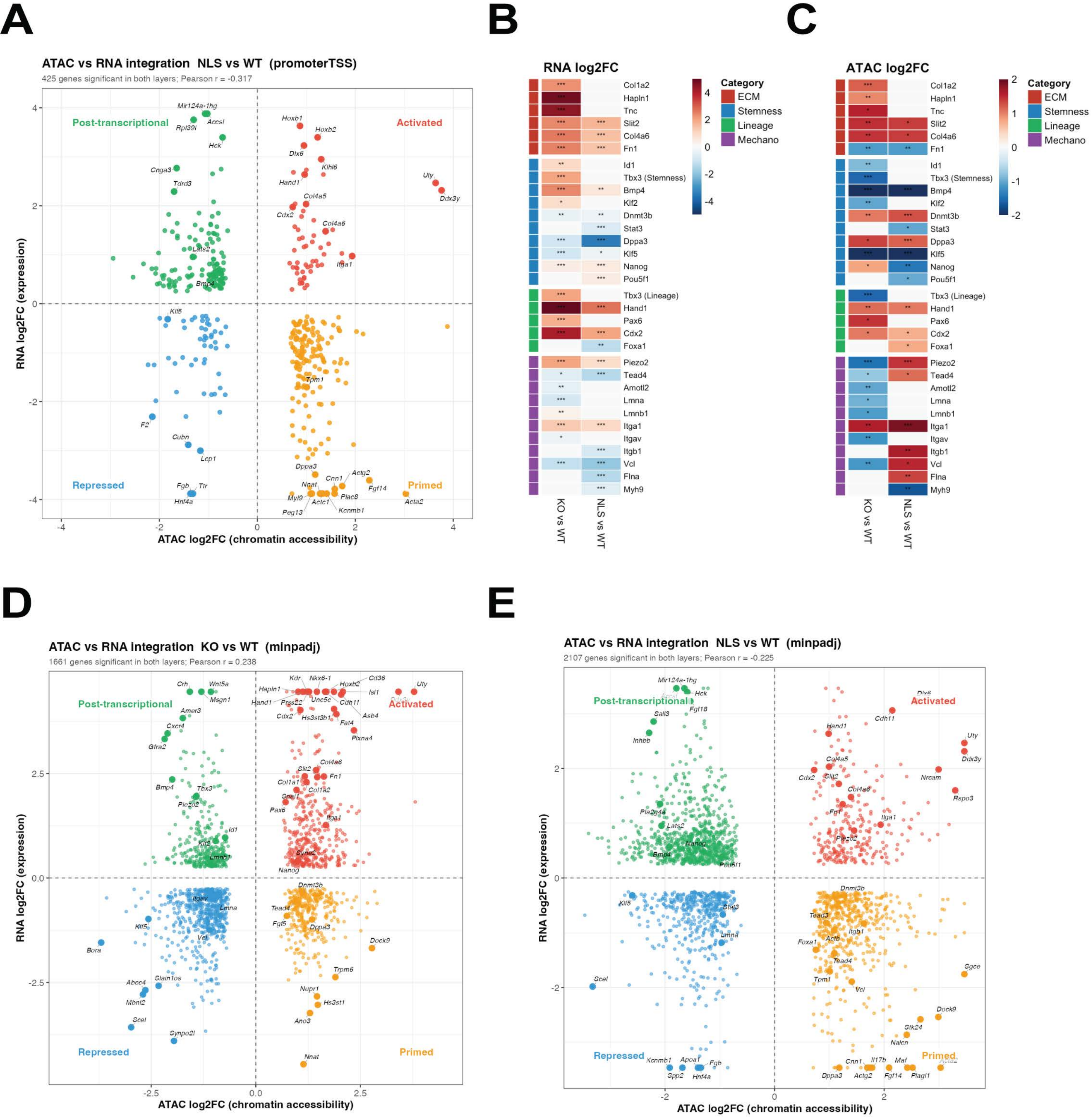
A
ATAC vs RNA integration NLS vs WT (promoterTSS)
425 genes significant in both layers; Pearson r = -0.317
Post-transcriptional
Activated
Repressed
Primed
RNA log2FC (expression)
ATAC log2FC (chromatin accessibility)
B
RNA log2FC
C
ATAC log2FC
Category
ECM
Stemness
Lineage
Mechano
Col1a2
Hapln1
Tnc
Slit2
Col4a6
Fn1
Id1
Tbx3 (Stemness)
Bmp4
Klf2
Dnmt3b
Stat3
Dppa3
Klf5
Nanog
Pou5f1
Tbx3 (Lineage)
Hand1
Pax6
Cdx2
Foxa1
Piezo2
Tead4
Amotl2
Lmna
Lmnb1
Itga1
Itgav
Itgb1
Vcl
Flna
Myh9
KO vs WT
NLS vs WT
D
ATAC vs RNA integration KO vs WT (minpadj)
1661 genes significant in both layers; Pearson r = 0.238
E
ATAC vs RNA integration NLS vs WT (minpadj)
2107 genes significant in both layers; Pearson r = -0.225

Supplementary figure 6

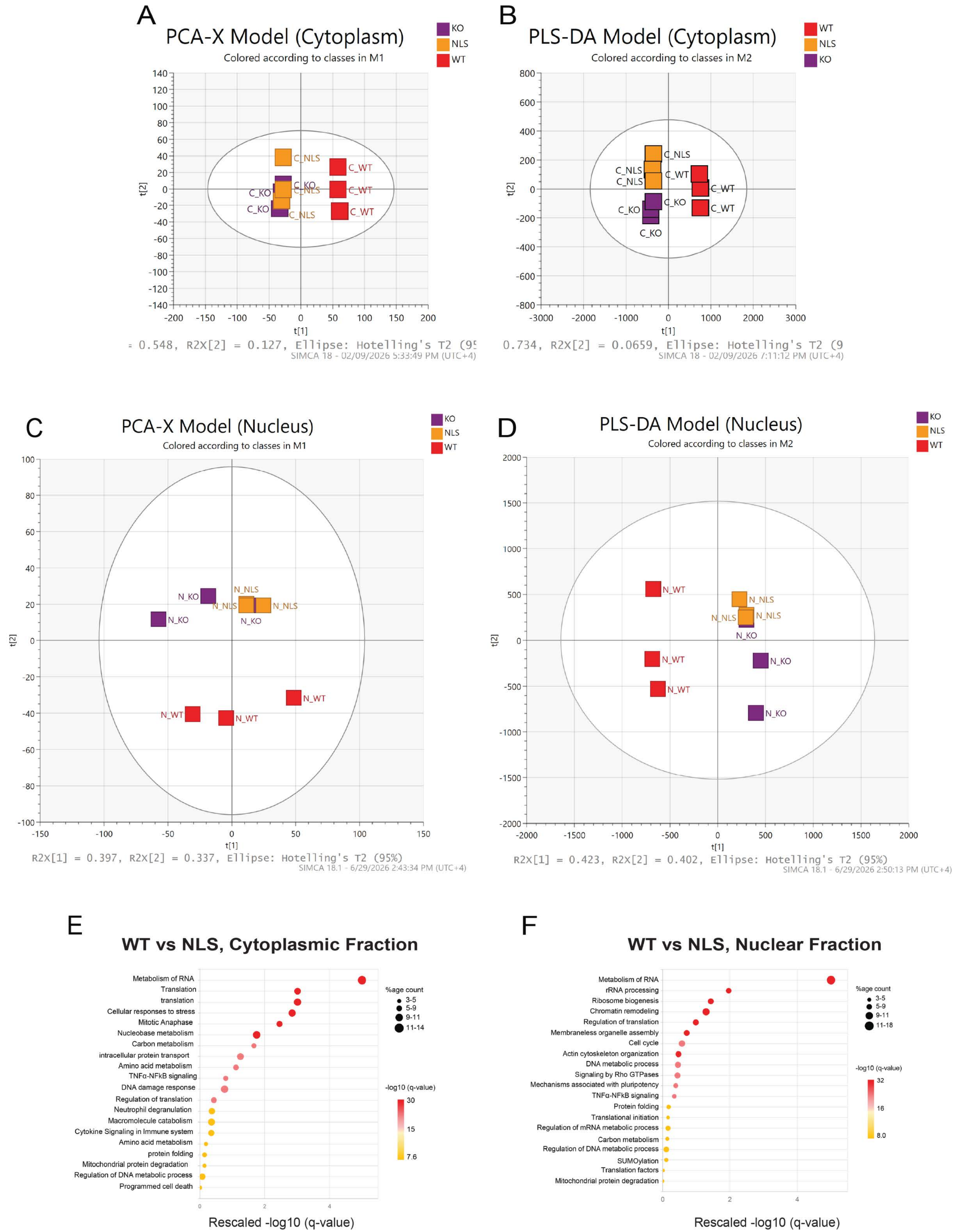

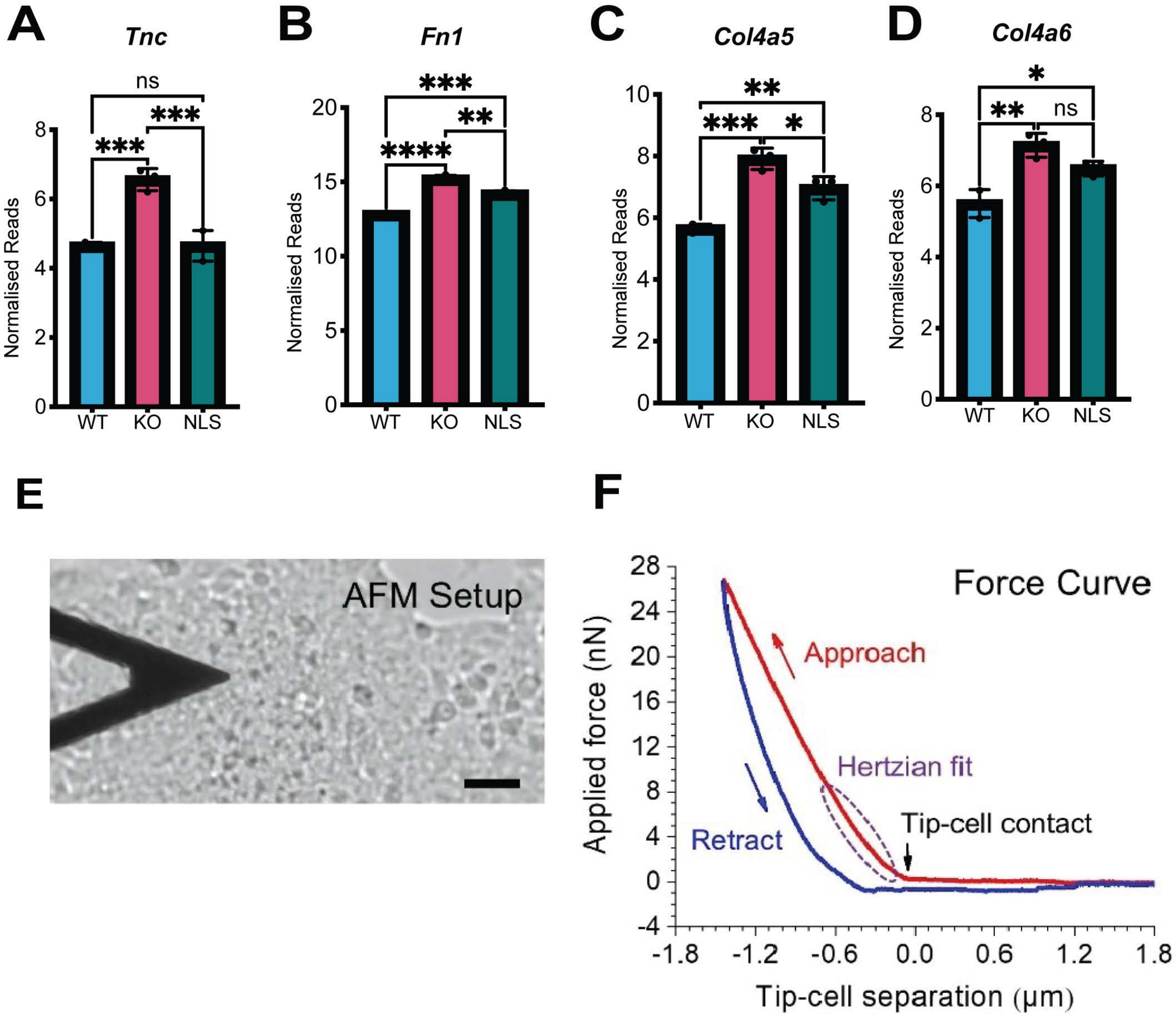
A
Tnc
ns
***
***
Normalised Reads
0
2
4
6
8
WT
KO
NLS
B
Fn1
***
**
****
Normalised Reads
0
5
10
15
20
WT
KO
NLS
C
Col4a5
**
***
*
Normalised Reads
0
2
4
6
8
10
WT
KO
NLS
D
Col4a6
*
**
ns
Normalised Reads
0
2
4
6
8
WT
KO
NLS
E
AFM Setup
F
Force Curve
Approach
Hertzian fit
Tip-cell contact
Retract
Applied force (nN)
Tip-cell separation (μm)
-4
0
4
8
12
16
20
24
28
-1.8
-1.2
-0.6
0.0
0.6
1.2
1.8

### RNA-Seq at Day 14

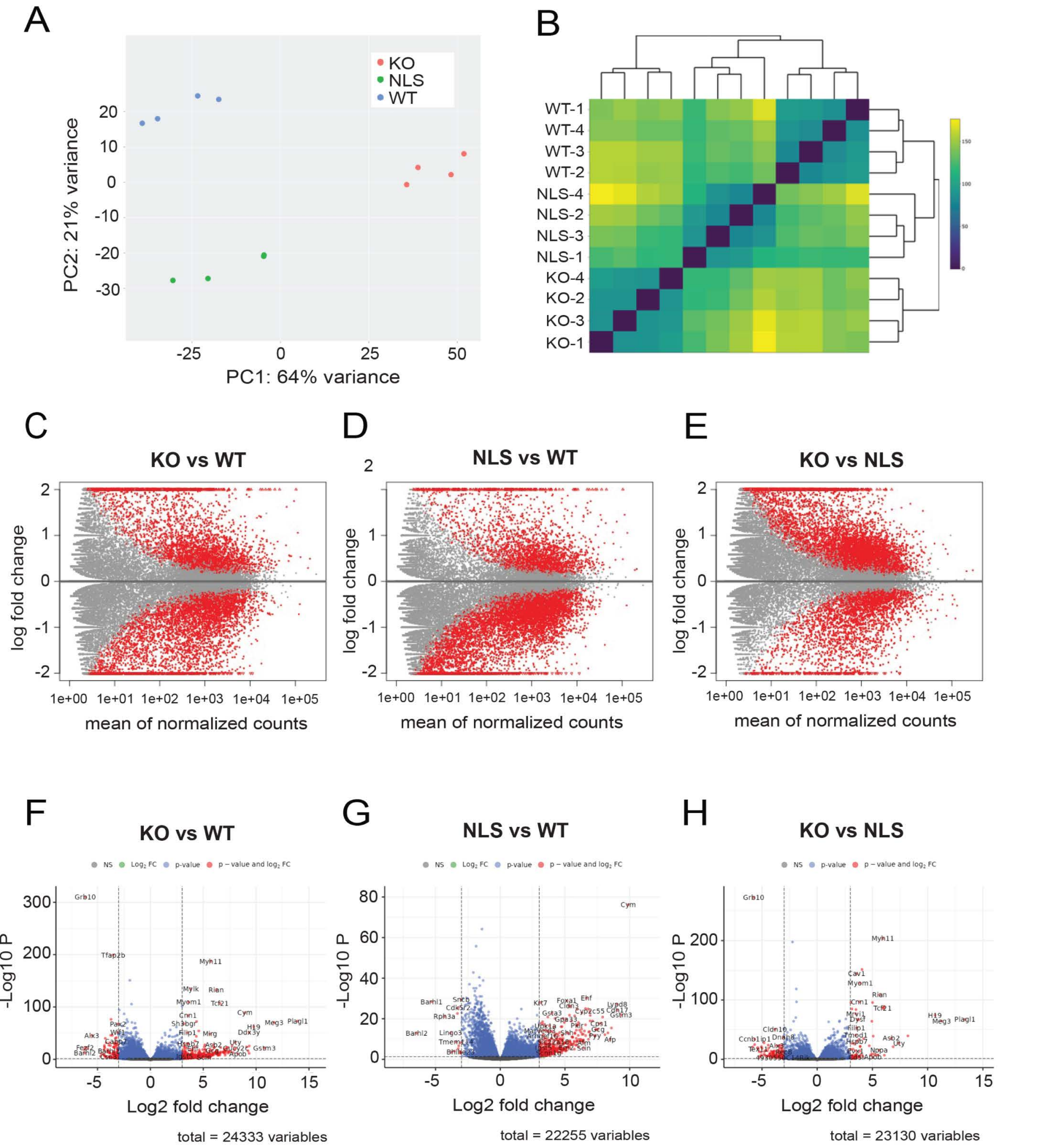

### KO mESCs vs WT mESCs, Day 14

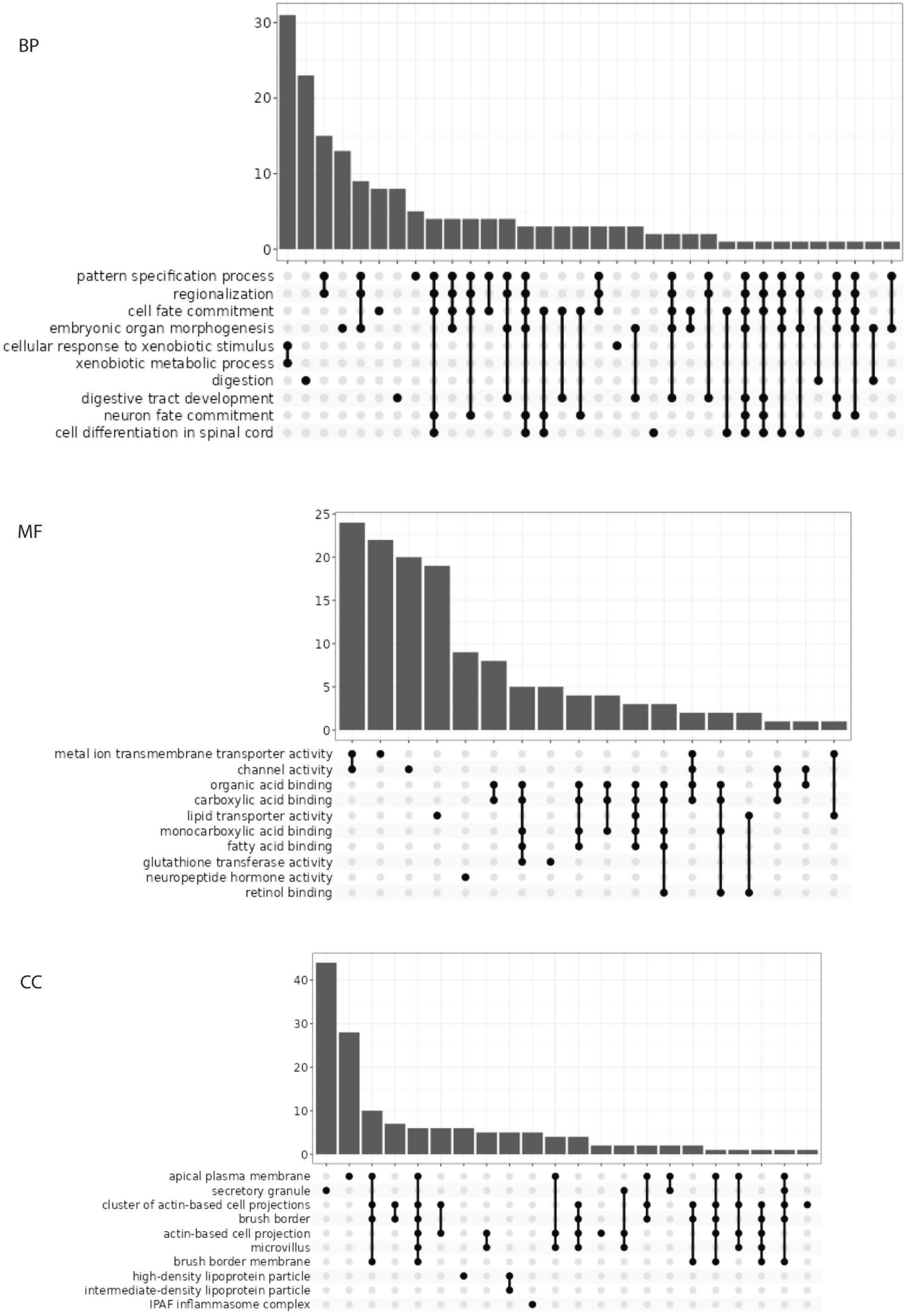

### RNA-Seq at Day 4

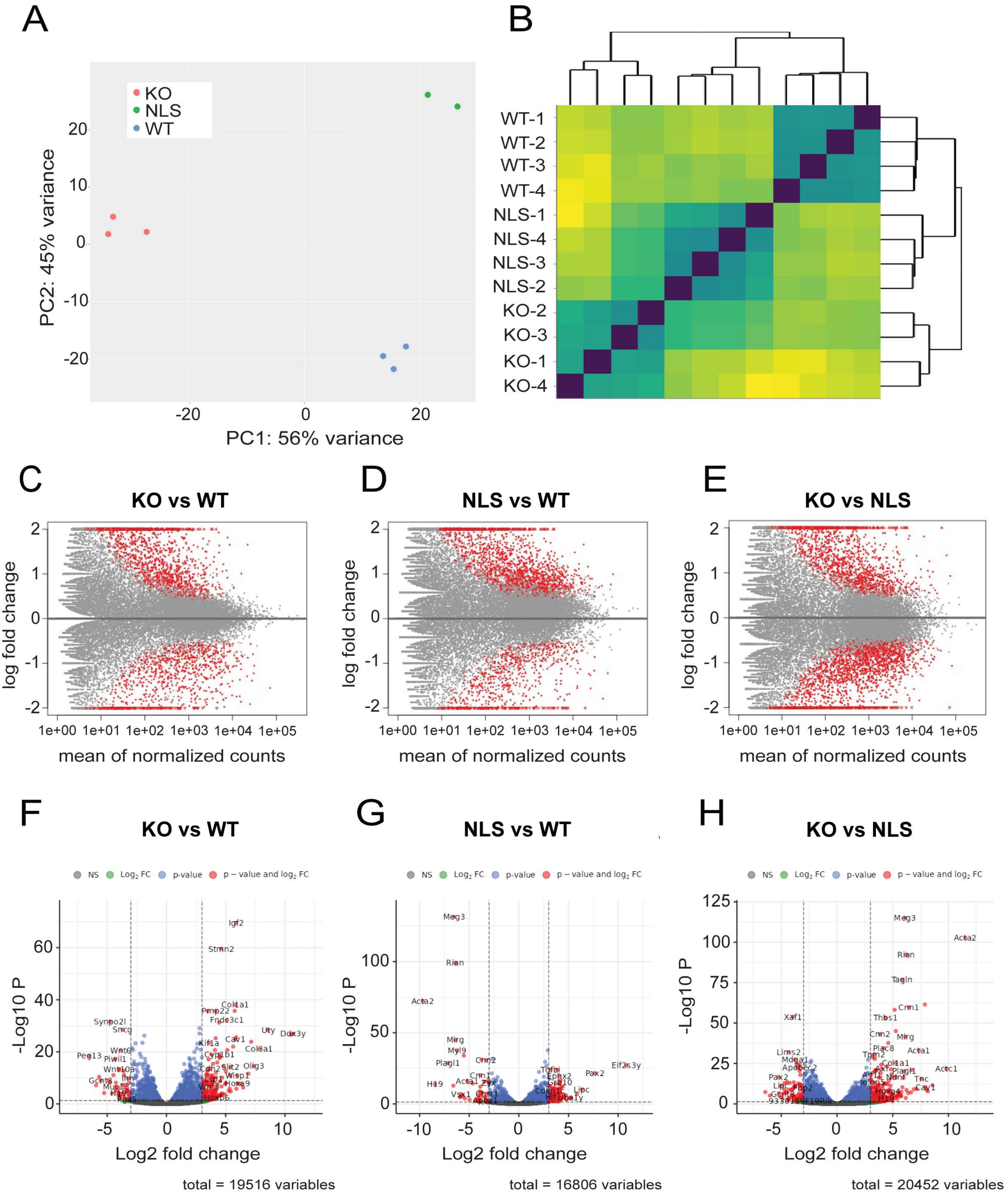

### KO mESCs vs WT mESCs, Day 4

BP

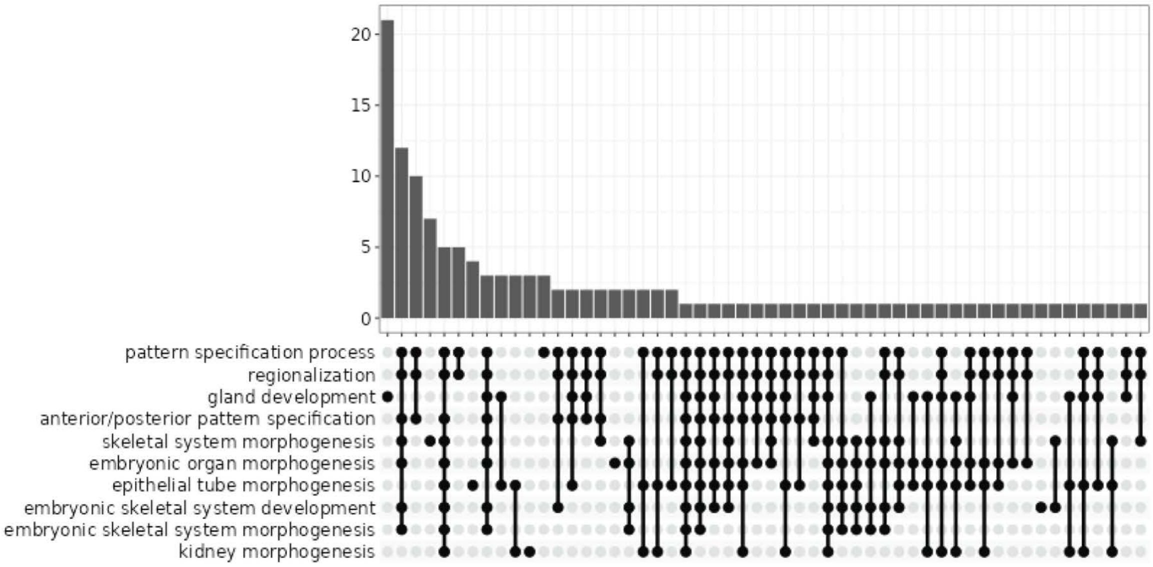


MF

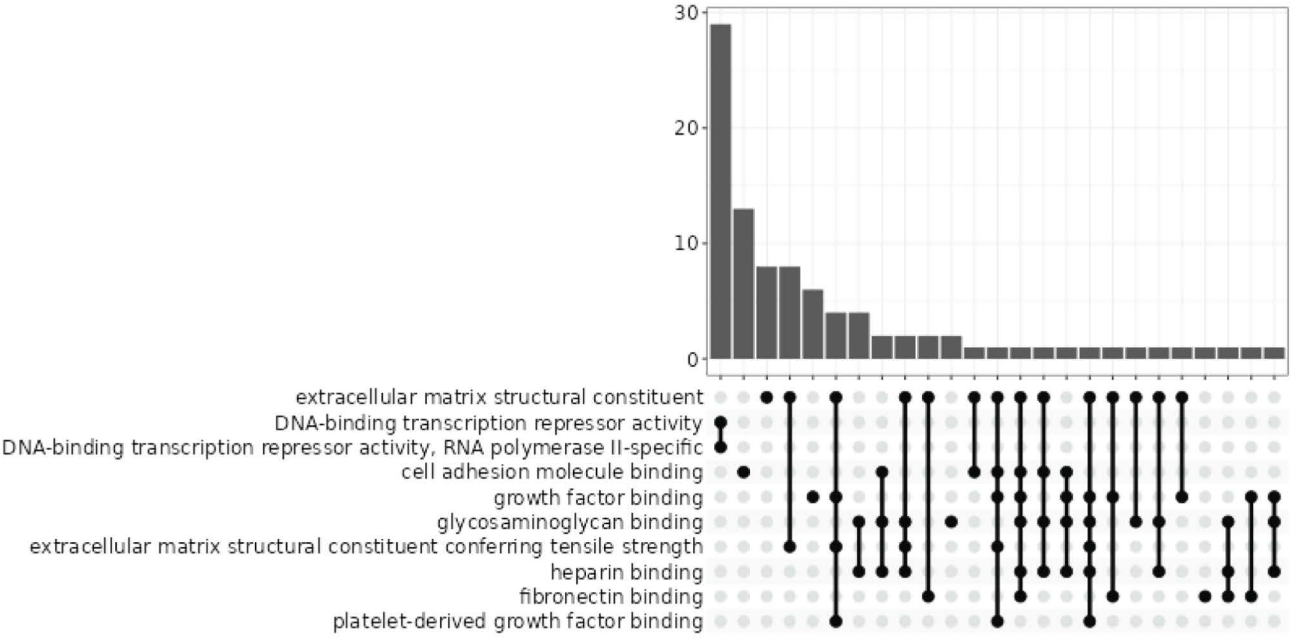


CC

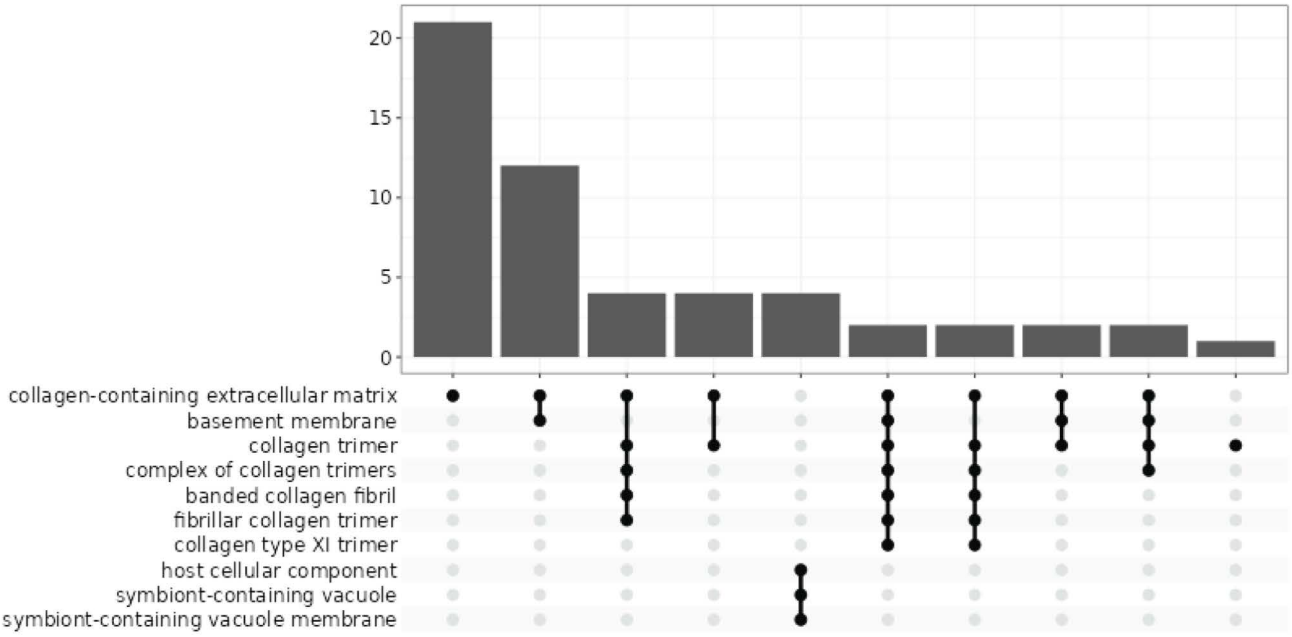

### RNA-Seq at Day 7

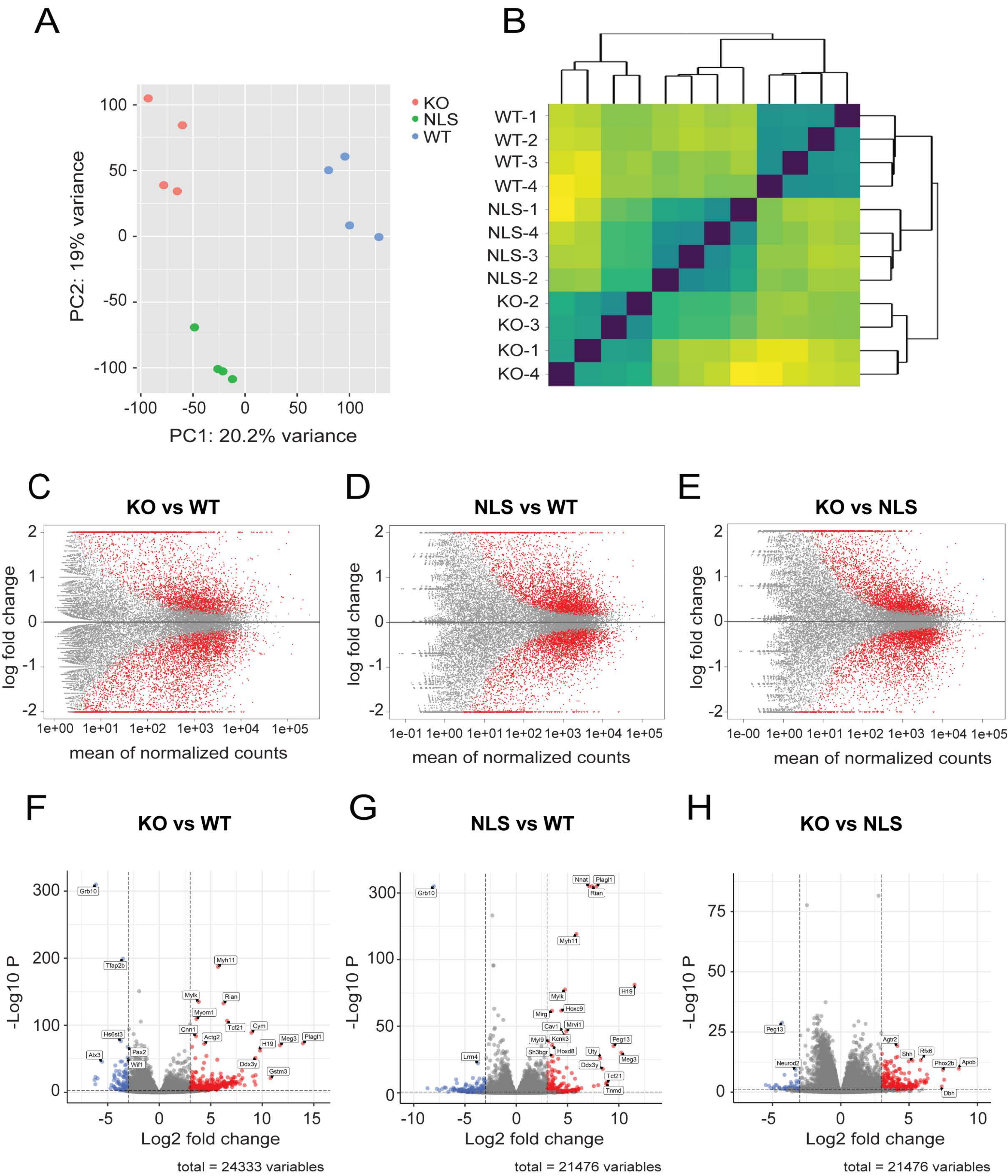

### KO mESCs vs WT mESCs, Day 7

BP

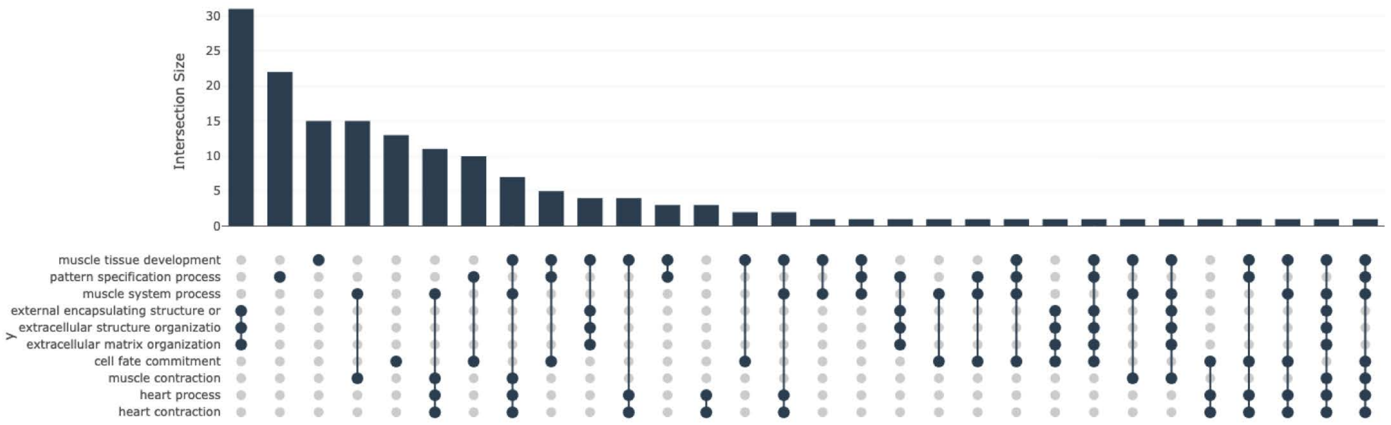


MF

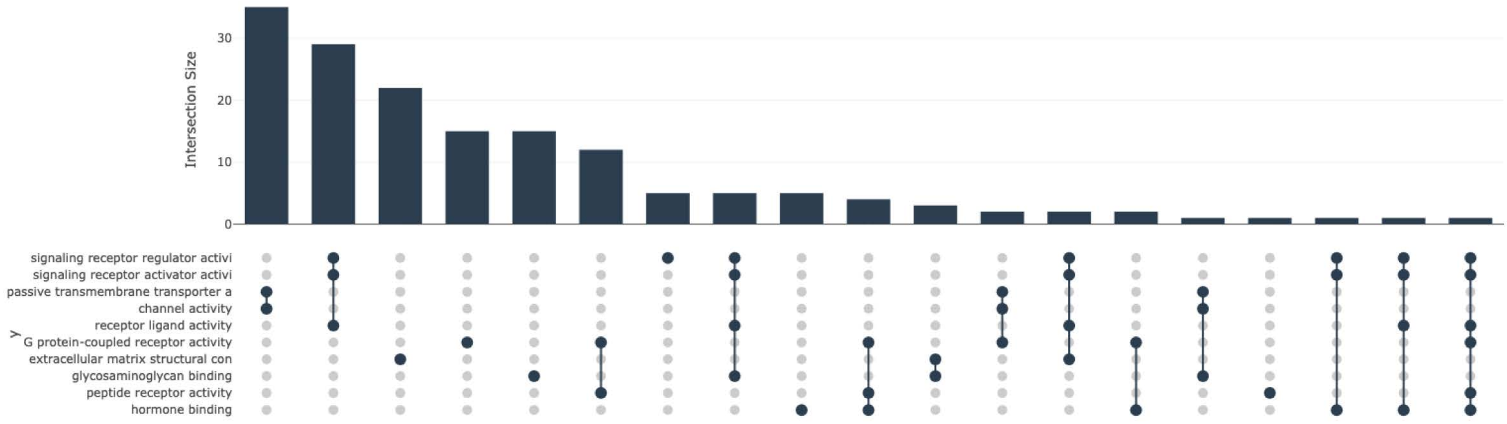


CC

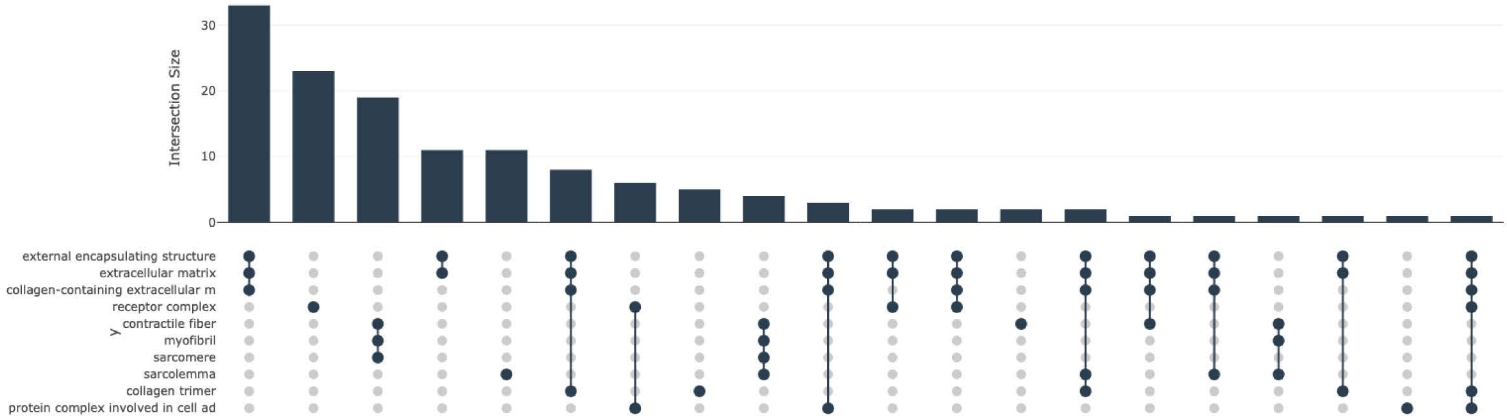
